# Preserving Native Cellulose–Xylan Architecture Enables Structure–Property Control in Holocellulose Nanofibrils and High-Performance Sustainable Materials

**DOI:** 10.64898/2026.06.07.730583

**Authors:** Parveen Kumar Deralia, Rosalie Cresswell, Yoshihisa Yoshimi, Tomohiro Kuga, Alberto Echevarría-Poza, Phil Howell, Alan Dickson, Marie Joo Le Guen, Edward Wagner, Amadeus C. S. de Alcântara, Carlos Guilherme Tissi Batista, Nadège Follain, Alyssa Miller, Michele Vendruscolo, Stefan J. Hill, Johnny Beaugrand, Munir S. Skaf, Daniel J. Cosgrove, Steven P. Brown, James A. Elliott, Ray Dupree, Paul Dupree

## Abstract

The hierarchical organization of cellulose microfibrils and their intimate interactions with hemicelluloses such as xylan underpin the exceptional mechanical performance of plant cell walls. However, translating these biological design principles into sustainable nanocellulosic materials remains limited by conventional cellulose nanofibril production routes, which rely on harsh chemical treatments that disrupt the native cellulose–hemicellulose architecture. Here, we present an optimized isolation strategy for holocellulose nanofibrils (hCNFs) that preserves native cellulose structure, xylan substitution and conformation, and cellulose–xylan interactions. Using wild-type *Arabidopsis thaliana*, a xylan glucuronidation-deficient *gux1/2* mutant, and *Brassica napus* straw as model systems, we systematically elucidate how xylan content and substitution pattern govern nanofibril isolation, interfacial interactions, and macroscopic properties. Two-dimensional ^13^C magic-angle spinning NMR demonstrates retention of native cellulose glucosyl environments, the presence of two-fold and three-fold helical xylan conformations, and cellulose-associated two-fold helical xylan. Cryogenic transmission electron microscopy reveals fibril widths of ∼3 nm, consistent with elementary cellulose Iβ microfibrils. We show that xylan glucuronidation regulates colloidal stability, hydration behavior, and interfibrillar cohesion, whereas xylan content controls nanofibrillation efficiency. These multiscale structural features translate directly into moisture sorption, thermal behavior, and mechanical performance. Notably, *Brassica napus* hCNF films exhibit exceptional strength and extensibility, surpassing many chemically modified CNF systems. This work demonstrates that preserving the native cellulose–hemicellulose architecture enables high-performance, sustainable nanocellulosic materials without chemical reconstruction.

## 1. Introduction

Lignocellulosic biomaterials and biochemicals are central to the development of a sustainable bioeconomy. Plant cell walls are composed of cellulose microfibrils embedded in a matrix of hemicelluloses and aromatic lignin polymers. Cellulose consists of β-1,4-linked glucan chains that assemble into microfibrils approximately 3–4 nm in diameter through intra- and intermolecular hydrogen bonding and stacking interactions. Hemicelluloses, including xylan and glucomannan, associate with specific cellulose microfibril surfaces in defined conformations, thereby influencing both structural organization and mechanical function.^[1–4]^

Xylan, the dominant hemicellulose in the secondary cell walls of higher plants, comprises a β-1,4-linked xylopyranosyl backbone variably substituted with acetyl, arabinofuranosyl, glucuronosyl, and feruloyl side groups, imparting substantial chemical diversity across plant species.^[5,6]^ In native cell walls, this hierarchical polymer network confers remarkable mechanical resilience and biological functionality. However, the same structural complexity poses a major challenge for isolating cellulose nanofibrils (CNFs) directly from biomass while retaining the native polymer architecture.

Conventional CNF production typically involves delignification through kraft or sulfite pulping, followed by one or more chemical, enzymatic, or mechanical treatments to achieve fibrillation.^[7–12]^ These processes inevitably disrupt cellulose and hemicellulose structures and eliminate native cellulose–xylan interactions. For example, TEMPO-mediated oxidation introduces carboxylate groups and induces cellulose depolymerization,^[10]^ whereas alkaline pretreatments remove, depolymerize, or deacetylate hemicelluloses.^[10,11]^ As a result, most CNFs differ fundamentally from native cellulose microfibrils in both structure and surface chemistry.

Developing alternative isolation strategies that preserve the native cellulose–hemicellulose architecture would enable the production of nanocellulosic materials that more closely resemble biological composites while reducing reliance on energy-intensive pulping processes. Achieving this goal requires analytical approaches capable of resolving molecular-level structure and interactions during nanofibril isolation and of correlating these features with macroscopic material properties.

Although hemicellulose content is known to influence nanofibrillation yield and enzymatic recalcitrance,^[5,13–15]^ most studies employ chemical or enzymatic pretreatments that fundamentally alter polymer structure.^[9,16–20]^ Because such treatments modify the very features of interest, the roles of hemicellulose substitution pattern and charge in fibril isolation and end-use performance remain poorly understood. Other investigations have instead removed hemicellulose entirely or adsorbed it onto chemically modified cellulose substrates,^[9,12,16–23]^ approaches that rarely recapitulate the complexity of native cell wall organization and therefore limit the production of materials with natural-like structure and function.

Genetic engineering offers a powerful means to selectively alter cell wall composition while maintaining native polymer assembly. *Arabidopsis thaliana gux1/2* mutants, which lack glucuronic acid substitutions on xylan, provide a unique system for probing how xylan fine structure governs cellulose–hemicellulose interactions.^[24]^ In parallel, *Brassica napus* represents a commercially relevant lignocellulosic residue with natural variation in xylan glucuronidation.^[25]^

Recent advances in the application of two-dimensional ^13^C magic-angle spinning solid-state NMR (2D ^13^C MAS ssNMR) have enabled direct, non-destructive characterization of cellulose and hemicellulose structure, conformation, and interactions in never-dried native cell walls.^[3,4,26,27]^ Unlike traditional techniques such as X-ray diffraction, infrared spectroscopy, and one-dimensional ^13^C MAS NMR (1D ^13^C MAS NMR), which often rely on peak deconvolution and provide only relative crystallinity estimates, 2D ^13^C MAS NMR resolves distinct glucosyl environments and directly identifies cellulose-associated hemicellulose conformations. Notably, several glucosyl residue environments that corresponding to different regions of the cellulose fibril can be resolved.^[27]^

Here, we integrate genetic variation, mild delignification, and multiscale characterization to isolate holocellulose nanofibrils (Hemicellulose-preserved CNFs are referred to as holocellulose nanofibrils (hCNFs)) that preserve native cellulose structure, xylan structure and conformation, and cellulose–xylan interactions. Using wild-type *Arabidopsis thaliana*, *gux1/2* mutants, and *Brassica napus* straw, we establish direct links between xylan content and structure, nanofibrillation efficiency, colloidal stability, hydration behavior, and the mechanical performance of the resulting nanopaper films. Our results demonstrate that preserving the native cellulose–hemicellulose architecture enables high-performance, sustainable nanocellulosic materials without chemical reconstruction.

## 2. Results

### 2.1 Holocellulose nanofibril isolation and preservation of xylan substitution pattern

#### 2.1.1 Mild delignification and mechanical processing yield partially individualized hCNFs

To examine how the native cellulose–xylan architecture governs nanofibril isolation and material properties, hCNFs were prepared from three lignocellulosic (LC) sources: *Arabidopsis thaliana* wild-type (*At* WT), the xylan glucuronidation-deficient *Arabidopsis thaliana* mutant (*At gux1/2*), and *Brassica napus* wild-type (*Bn* WT) straw (Figure 1a).

**Figure 1.**
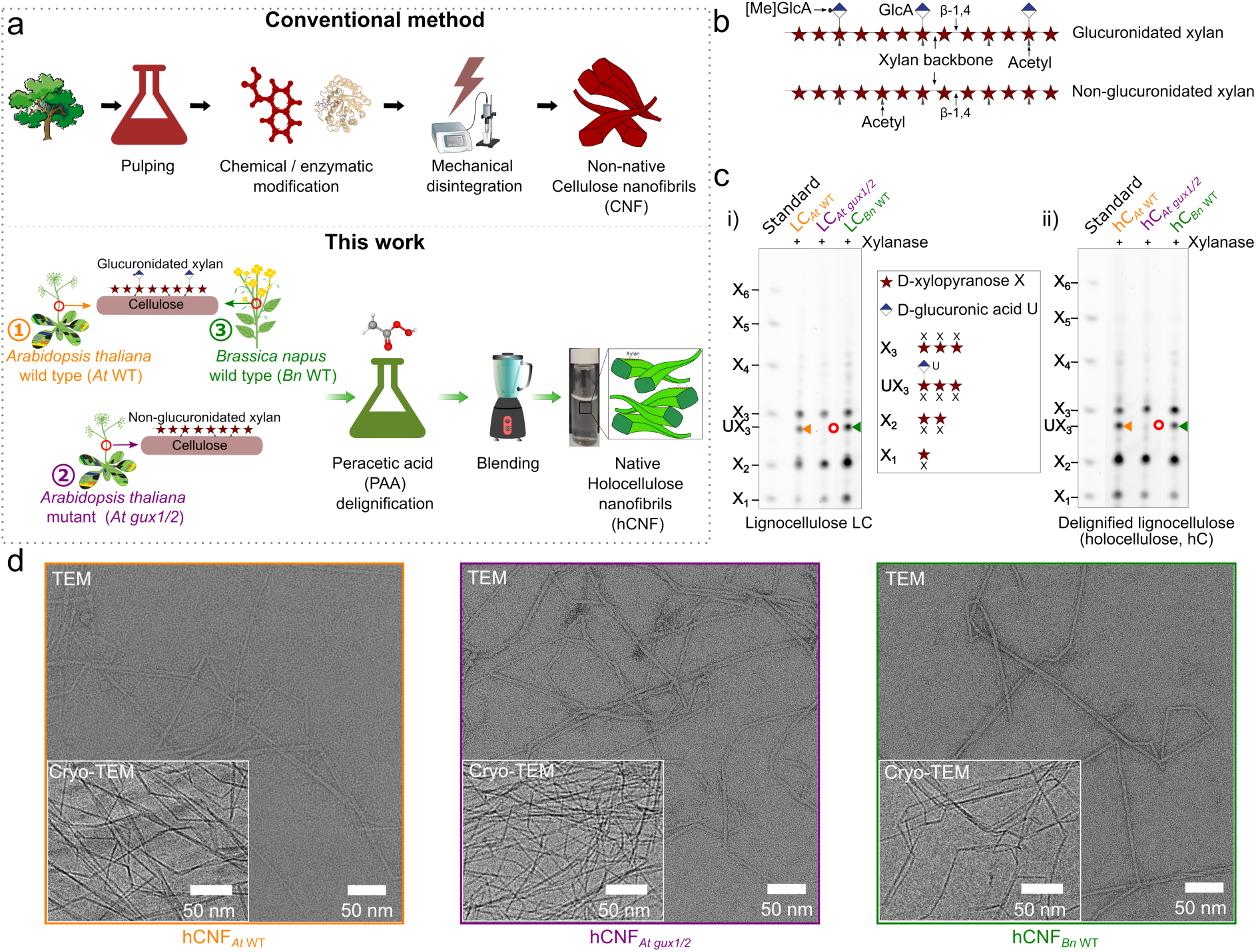
Preparation of hCNFs with preserved xylan substitution pattern. a) Schematic comparison of conventional cellulose nanofibril preparation with the hCNF isolation workflow used here for three lignocellulose (LC) sources: ① *Arabidopsis thaliana* wild-type (*At* WT, orange), ② *Arabidopsis thaliana* variant (*At gux1/2,* purple, genetically modified), and ③ *Brassica napus* wild-type (*Bn* WT, green). b) Schematic representation of glucuronidated and non-glucuronidated xylan, each consisting of a β-1,4-linked D-xylosyl backbone bearing *O*-acetyl substituents, with (glucuronidated) or without (non-glucuronidated) 4-*O*-methyl-D-glucuronic acid ([Me]GlcA) side groups; see Figure S2a for the chemical structure. c) Polysaccharide Analysis by Carbohydrate gel Electrophoresis (PACE) of *Cj*GH10 xylanase digestion products from lignocellulose (LC) and holocellulose (hC), used to assess glucuronic acid (GlcA) substitution patterns on xylan. LC and hC from *At* WT and *Bn* WT show UX_3_ digestion products (orange and green arrowheads), whereas *At gux1/2* lacks UX_3_ (red circle). X_1_–X_6_ denote xylosyl oligosaccharide standards. The xylanase action scheme and original gels are shown in Figures S2 and S4. d) Representative TEM micrograph of hCNFs; inset shows a cryo-TEM image. Individual fibrils (blue markers) and loosely bundled fibrils are shown in Figure S7.

The lignocellulosic materials, sample nomenclature, acronyms, and symbols are summarized in Tables S1–S3, and the lignin and neutral monosaccharide contents of the LCs are shown in Figure S1.

Xylan substitution patterns in the native lignocelluloses were analyzed using polysaccharide analysis by carbohydrate gel electrophoresis (PACE) following digestion with *Cj*GH10 xylanase (see Figures S2a,b for xylan chemical structure and digestion scheme). As expected, *At* WT and *Bn* WT lignocelluloses released glucuronidated UX_3_ oligosaccharides, whereas *At gux1/2* lacked these products, confirming the absence of glucuronic acid substitutions on xylan in the mutant (Figure 1c-i), consistent with previous reports.^[24,25]^ Xylan acetylation in *Bn* WT was verified by xylanase digestion of DMSO-extracted xylan (Figure S3).^[28]^

Unlike conventional CNF production routes (Figure 1a), which involve alkaline pulping and oxidative pretreatments that disrupt cellulose and hemicellulose structure,^[10,11]^ a mild peracetic acid (3% w/v) delignification was employed without alkaline washing, in contrast to previously reported methods.^[9,18]^ This approach effectively removed lignin while preserving xylan content, yielding xylan-rich holocellulose (hC). Subsequent mechanical fibrillation in a high-speed blender produced polydisperse suspensions; translucent, colloidally stable hCNF fractions were then recovered from the supernatant after centrifugation (see Figures S5–S6 for the hCNF separation scheme and images of the hCNF suspensions).

Electron microscopy (TEM and cryo-TEM; Figure 1d and Figure S7) and AFM (Figure S8) revealed long, thin, partially individualized nanofibrils, consistent with previous reports on holocellulose-derived CNFs.^[9,18,29]^ Sugar analysis confirmed the retention of xylan in both hCs and hCNFs (Figures S9–S10), consistent with previous studies.^[9,18]^ Notably, hCNFs derived from *At gux1/2* contained a higher xylan fraction than those from *At* WT, suggesting stronger xylan–cellulose association in the absence of glucuronic acid substitutions.^[30,31]^

#### 2.1.2 Xylan substitution pattern in hCs and hCNFs remains identical to that in native lignocellulose

To determine whether the xylan substitution pattern was altered during hCNF preparation, xylan glucuronidation and acetylation were examined in hCs and hCNFs by PACE following GH10 xylanase digestion. Glucuronidated UX_3_ oligosaccharides were detected in hCs from *At* WT and *Bn* WT but were absent in *At gux1/2*, confirming the preservation of native glucuronidation patterns through delignification (Figure 1c-ii).

Xylan acetylation, which plays a key role in cellulose–xylan interactions,^[3,28]^ was assessed in *Bn* WT samples by extracting acetylated xylan with DMSO prior to enzymatic digestion. The oligosaccharide profiles obtained from lignocellulose, hC, and hCNF samples were nearly identical, whereas non-acetylated xylan yielded shorter fragments (Figure S3). These results are consistent with a previous report on *Arabidopsis thaliana* acetylation.^[28]^ Together, they demonstrate that both the glucuronidation and acetylation patterns of xylan remain unchanged throughout hCNF isolation.

### 2.2 Preservation of cellulose structure and xylan helical conformation in hCNFs

#### 2.2.1 MAS NMR confirms retention of native cellulose structure in hCNFs

The preservation of cellulose structure and xylan conformation following delignification and fibrillation was investigated using 1D and 2D ^13^C MAS NMR on uniformly ^13^C labeled *At* WT lignocellulose (^13^C labeled LC*_At_* _WT_), holocellulose (^13^C labeled hC*_At_* _WT_), and hCNFs (^13^C labeled hCNF*_At_* _WT_). 1D ^13^C CP MAS spectra confirmed the complete removal of lignin, as evidenced by the disappearance of methoxyl signals at ∼56 ppm (Figures 2a and S11a). In contrast, xylan acetyl methyl signals at ∼21 ppm were retained, indicating preservation of acetylation.

**Figure 2.**
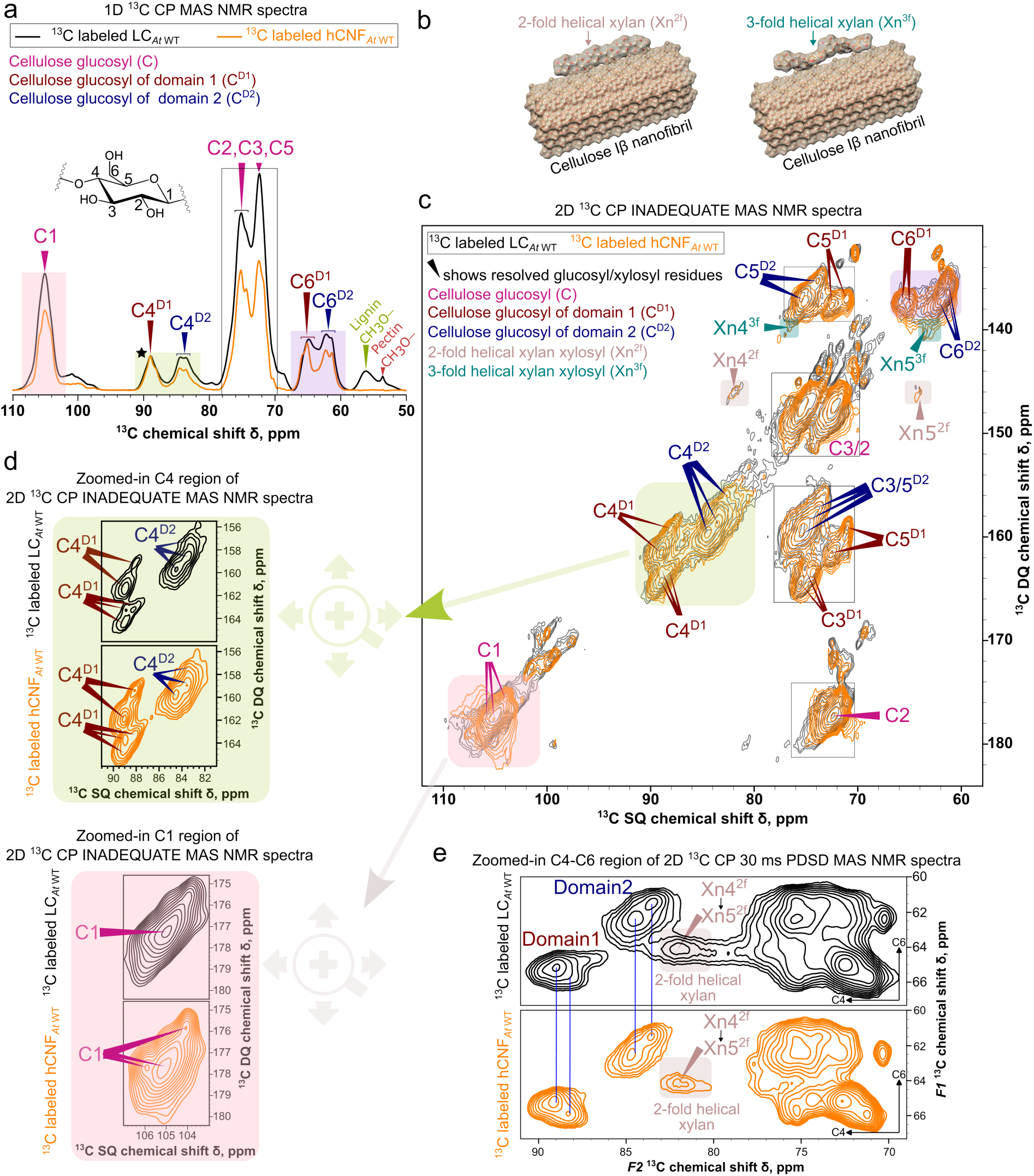
MAS NMR evidence for preservation of native cellulose structure and xylan conformations in hCNFs. a) Overlay of 1D ^13^C CP MAS NMR spectra of ^13^C-labeled LC*_At_* _WT_ and hCNF*_At_* _WT_. Resonances C1–C6 correspond to the glucosyl carbons of cellulose. Native plant cellulose exhibits two spectral domains — domain 1 (dark-red markers) and domain 2 (blue markers) — distinguished by chemical-shift differences at C4 and C6.^[26,32]^ Spectra are normalized to the C4^D1^ signal (asterisked) at ∼89 ppm. The full 0–200 ppm spectral overlay is shown in Figure S11a. b) Schematic illustrating the compatibility of 2-fold and 3-fold helical xylan conformations with the cellulose surface. c) Overlay of 2D ^13^C CP refocused INADEQUATE MAS NMR spectra of ^13^C labeled LC*_At_* _WT_ and hCNF*_At_* _WT_, showing closely matching cellulose glucosyl environments (markers). To aid comparison, shaded boxes mark the same spectral regions in the 1D and 2D spectra — the C1 (pink), C4 (olive-green), and C6 (purple) regions — illustrating how resonances that overlap in 1D are resolved in the 2D INADEQUATE spectrum. Cellulose domain 1 and domain 2 environments are indicated by dark-red and blue markers, respectively; 2-fold (Xn^2f^) and 3-fold (Xn^3f^) helical xylan peaks are highlighted in rose-brown and teal, respectively. The ^13^C chemical shifts of cellulose glucosyl carbons for LC*_At_* _WT_ and hCNF*_At_* _WT_ are listed in Table S4 for comparison. d) Zoomed-in views of the C1 and C4 regions from panel c, demonstrating improved resolution of distinct glucosyl environments in cellulose for hCNFs. e) Zoomed-in C4–C6 and 2-fold helical xylan (Xn4^2f^ → Xn5^2f^ cross peak) regions from 2D ^13^C CP 30 ms PDSD MAS NMR spectra of ^13^C labeled LC*_At_* _WT_ and hCNF*_At_* _WT_ revealing preservation of cellulose–2-fold helical xylan interactions in hCNFs. All MAS NMR experiments were performed at a ^1^H Larmor frequency of 1.0 GHz and an MAS frequency of 12.5 kHz. See Section 3.2 of the Supporting Information for further experimental details.

The carbohydrate region (50–110 ppm) showed characteristic cellulose resonances corresponding to the C1–C6 carbons (Figure 2a). The C4 (∼80–90 ppm) and C6 (∼60–65 ppm) regions exhibited the well-known bifurcation into spectral domain 1 (C4/6^D1^) and domain 2 (C4/6^D2^) signals, reflecting distinct glucosyl environments associated with various fibril core and surface regions.^[26,27,32]^ Comparison of LC, hC, and hCNF spectra revealed that delignification alone preserved the relative distribution of these domains, whereas isolation of the colloidal hCNF fraction from the blended hC suspension (see scheme in Figure S5) increased the C4^D1^ and C4^D2^ ratio (Figures 2a and S11b). This change likely reflects enrichment of more ordered fibrils and removal of less ordered material during centrifugation, rather than the modification of cellulose structure itself.

To resolve overlapping signals in the 1D spectra, 2D ^13^C CP-refocused INADEQUATE MAS NMR experiments were performed. This method correlates pairs of single-quantum (SQ) signals from directly bonded carbon atoms (e.g., C1–C2 in cellulose glucosyl units) through their double-quantum (DQ) chemical shifts, substantially enhancing the resolution of distinct glucosyl and xylosyl environments that have otherwise similar chemical shifts.^[3,4,26,27]^ In addition, cross-polarization excitation selectively enhances signals from the relatively rigid (immobile) fractions of cellulose and hemicellulose.^[33]^

The resulting spectra revealed multiple well-resolved cellulose glucosyl resonances in both LC*_At_* _WT_ and hCNF*_At_* _WT_ (indicated by markers in Figure 2c–d), with improved resolution in the hCNFs (Figure S11c). This improvement is evident from the appearance of additional resolved peaks (for example, in the zoomed-in C1 region of Figure 2d), accompanied by an increase in signal-to-noise in certain spectral regions (for example, the zoomed-in C4 region of Figure 2d). Importantly, the corresponding cellulose glucosyl ^13^C chemical shifts essentially remained unchanged, differing in nearly all cases by no more than the experimental uncertainty (0.1 ppm; Figure 2c and Table S4) — comparable to the very small shift changes reported for the isolation of poplar hCNFs^[27]^ and demonstrating that the molecular structure of cellulose is preserved after hCNF isolation from the cell wall.

#### 2.2.2 Presence of 2-fold and 3-fold helical xylan conformations in hCNFs

In native plant cell walls, xylan adopts both 2-fold and 3-fold helical conformations, with the 2-fold form specifically associated with cellulose fibril surfaces (Figure 2b).^[3]^ The 2D ¹³C CP-refocused INADEQUATE spectra revealed the characteristic xylan resonances in both LC*_At_* _WT_ and hCNF*_At_* _WT_ (Figure 2c). The carbon-4 signals appeared at ∼82.2 ppm (Xn4^2f^) and ∼72.4 ppm (Xn4^3f^), and the carbon-5 signals at ∼64 ppm (Xn5^2f^ and Xn5^3f^). Peaks of the 2-fold helical conformation (Xn^2f^) are highlighted in rose-brown and those of the 3-fold helical conformation (Xn^3f^) in teal. These signals confirm that both xylan conformations coexist after fibrillation. Their intensities, normalized to cellulose, were modestly reduced in the hCNFs, consistent with the higher glucose-to-xylose ratio measured in the hCNFs (9.3; Figure S10b) relative to the parent LC*_At_* _WT_ (4.1; Figure S1d). This reflects a relative depletion of xylan, likely due to preferential retention of cellulose-rich fibrils during the selective collection of the colloidal hCNF fraction (See the separation scheme in Figure S5).

To probe cellulose–xylan interactions, 2D ¹³C CP proton-driven spin diffusion (PDSD) MAS NMR experiments were performed with a 30 ms mixing time, which reports short-range ¹³C–¹³C proximities. Clear cross-peaks between the intramolecular xylan C4 and C5 carbons of the 2-fold helical conformation (Xn4^2f^ → Xn5^2f^) were observed in both LC and hCNF samples (Figure 2e). Because this 2-fold helical conformation is known to depend on cellulose binding,^[3]^ the cross-peak serves as an intramolecular marker of cellulose-bound 2-fold xylan — rather than a direct cellulose–xylan contact — providing indirect evidence that cellulose-associated 2-fold helical xylan persists in the hCNFs. Together, these results indicate that native xylan conformations and their interactions with cellulose surfaces are retained through delignification and nanofibrillation.

### 2.3 Cellulose crystalline organization probed by wide-angle X-ray scattering

Wide-angle X-ray scattering (WAXS) was employed to assess whether hCNF isolation altered cellulose crystalline organization. All hCNF films exhibited diffraction patterns characteristic of cellulose Iβ, with the combined 1–10/110, 200, and 004 reflections (Figure 3a). Peak positions closely resembled those reported for wood^[34–37]^ and CNFs^[18,29]^, differing from those of highly crystalline cellulose sources such as cotton, bacterial, tunicate and valonia cellulose.^[35,38]^

**Figure 3.**
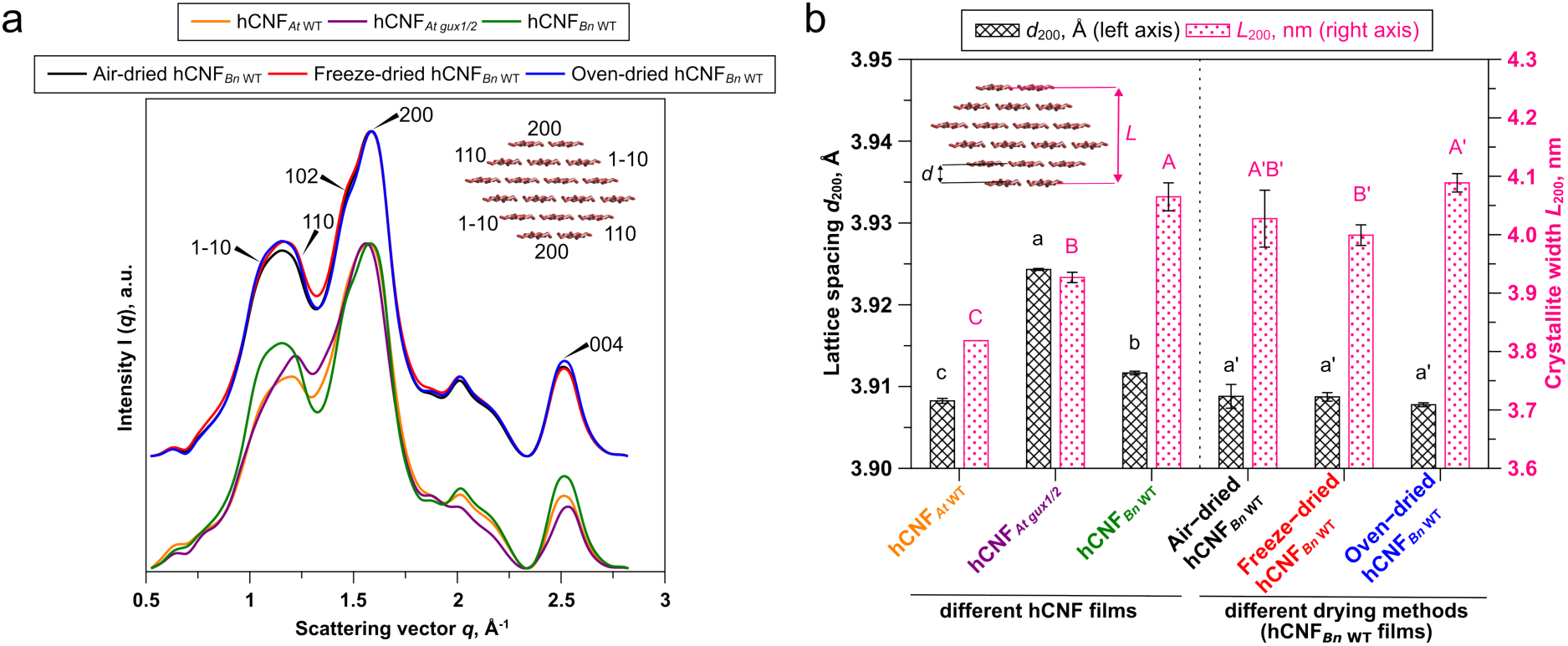
Wide-angle X-ray scattering (WAXS) analysis of fibril packing in hCNFs and the influence of drying methods. a) WAXS profiles of all hCNF samples (bottom three traces) and hCNF*_Bn_* _WT_ films prepared using different drying methods (top three traces), showing the characteristic diffraction features of cellulose Iβ (combined 1–10/110, 200, and 004 reflections). b) Lattice spacing (*d*_200_, left y-axis) and crystallite width (*L*_200_, right y-axis) derived from the 200 reflection after peak deconvolution (see Figure S12 for an example of the deconvolution). Data are presented as mean ± SD (n = 4). Columns not sharing the same letter are significantly different according to Tukey’s test at the 5% significance level.

Crystallite widths (*L*_200_) and interplanar spacings (*d*_200_) were extracted using a cellulose Iβ unit-cell model (Figure 3b).^[39]^ hCNF*_At gux1/2_* displayed a slightly larger *d*_200_ spacing than hCNF*_At_* _WT_ and hCNF*_Bn_* _WT_, potentially reflecting modified interfibrillar organization or denser xylan binding. In contrast, hCNF*_Bn_* _WT_ exhibited a modestly larger crystallite width (*L*_200_), consistent with limited interfibrillar co-crystallization or fibril fusion.^[40]^

To evaluate the influence of drying on cellulose nanofibril co-crystallization, hCNF*_Bn_* _WT_ films were prepared by air-drying, freeze-drying, and oven-drying. The drying method had negligible effects on d-spacing (*d*_200_) and only minor effects on crystallite width (*L*_200_) (Figure 3b), indicating that irreversible fibril fusion or extensive co-crystallization is limited under these conditions.

### 2.4 Fibril widths and comparison with cellulose Iβ models

#### 2.4.1 hCNF fibril widths are consistent across lignocellulosic sources

Fibril widths were quantified using both negatively stained TEM and cryo-TEM. TEM yielded average apparent widths of ∼4.8 nm (Figure 4a), whereas cryo-TEM measurements in the near-native hydrated state gave narrower widths of ∼3.0–3.2 nm, with hCNF*_At_* _WT_ being the smallest (3.07 nm). The larger mean widths measured by TEM are attributed to a systematic offset from staining artifacts, whereas the broad width distributions in both methods likely reflect intrinsic fibril-to-fibril variability and differences in fibril orientation during imaging. The TEM width values are consistent with or slightly larger than those reported for softwood hCNFs.^[18]^

**Figure 4.**
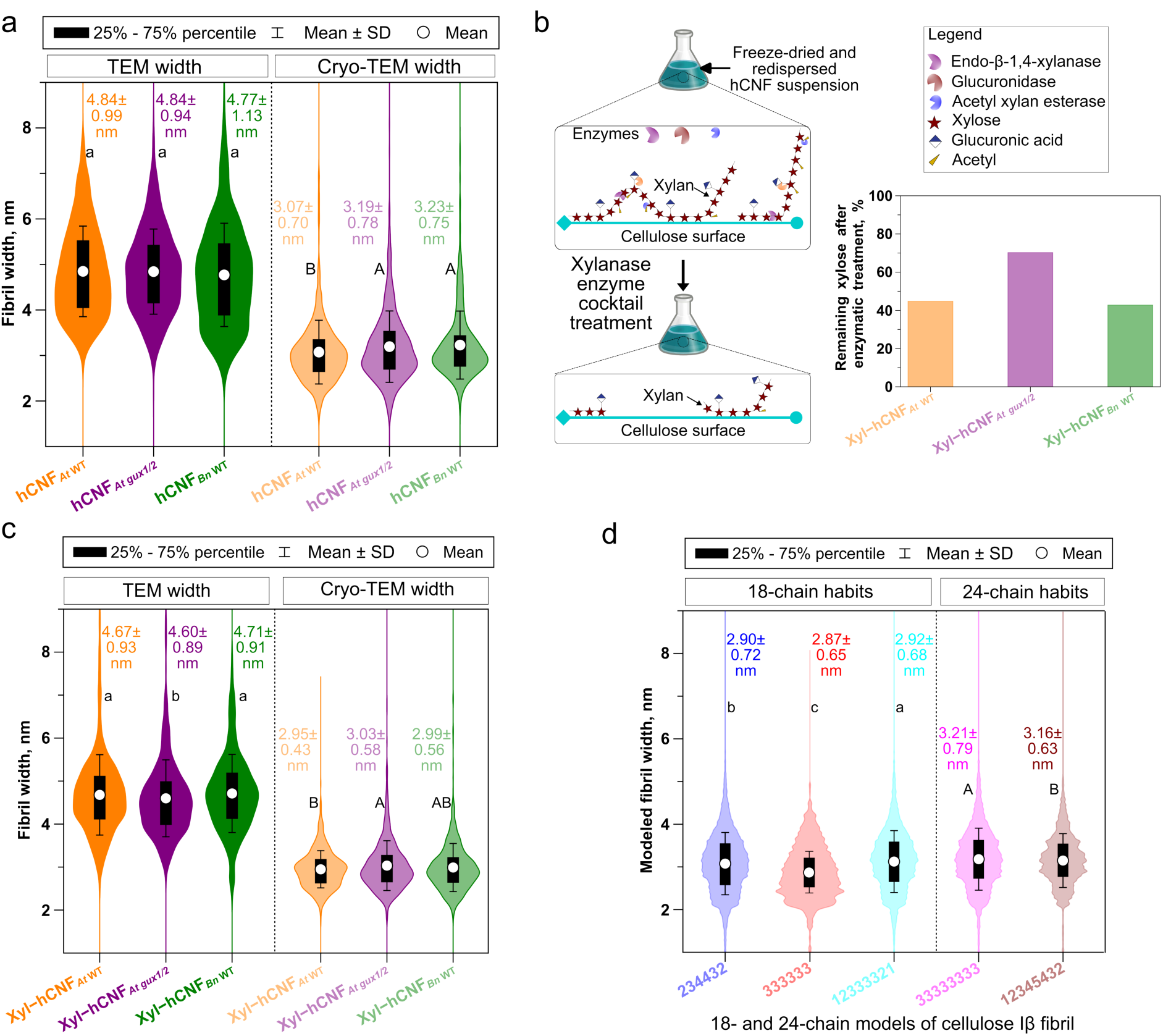
Cryo-TEM–derived fibril widths of xylanase-treated hCNFs (xyl-hCNfs) provide an experimental estimate of native xylan-bound hCNFs and agree with modeled cellulose Iβ microfibril dimensions. a) Width distributions of hCNFs measured from TEM and cryo-TEM micrographs, with corresponding mean values (n = 700–800). b) Schematic illustrating enzymatic treatment of hCNFs with a xylanase cocktail and the residual xylose remaining in xylanase-treated hCNFs (xyl-hCNFs). c) Width distribution and mean values for xyl-hCNFs determined from TEM and cryo-TEM micrographs (n = 700–800). d) Modeled fibril widths for 18- and 24-chain cellulose Iβ fibrils with different cross-sectional habits (234432, 333333, 12333321, 33333333, and 12345432). See Figures S16–S17 for details of the cellulose Iβ microfibril models and fibril size-distribution analysis. Means not sharing a common letter are significantly different as determined by Tukey’s test at the 5% significance level.

#### 2.4.2 Partial enzymatic removal of xylan modestly reduces fibril widths

To determine the contribution of xylan to the apparent fibril width, hCNFs were treated with a xylanase cocktail (GH10 xylanase, GH115 glucuronidase, CE4 acetyl esterase; Figure 4b; See Figure S13 for the xylanase-treated hCNF (xyl-hCNF) suspensions). This treatment removed ∼55–57% of xylan from *At* WT and *Bn* WT hCNFs, but only ∼30% from *At gux1/2*, indicating less accessible xylan association in the mutant. Previous studies have shown that xylanases hydrolyze only approximately 30–35% of the xylan associated with pulp-derived cellulose,^[17,41]^

similar to the results observed for At gux1/2 hCNFs. Cryo-TEM revealed small but consistent reductions in fibril width (0.1–0.2 nm) following enzymatic treatment (Figure 4c; see Figure S14 for statistical analysis), giving a genuine estimate of native naked fibril widths compared to earlier research that used either drying or processing techniques in which the sample was modified.^[18,42]^ A comparison of our fibril widths to those of previous literature supports this result (Figure S15). Together, these results indicate that partially removed xylan contributes only marginally to the apparent fibril width, and that cryo-TEM of xylanase-treated fibrils provides a more reliable estimate of the native fibril width than methods involving drying or sample processing.

#### 2.4.3 Cryo-TEM widths agree with modeled cellulose Iβ microfibrils

The experimentally measured widths of xylanase-treated hCNFs (2.95–3.03 nm) were compared with the theoretical apparent diameters of the widely accepted 18- and 24-chain cellulose Iβ microfibrils with different cross-sectional habits (Figure 4d). The experimental distributions closely matched those predicted for the 18-chain fibrils, although the 24-chain models could not be definitively excluded. These results are therefore consistent with current models of native cellulose microfibrils,^[43]^ though we cannot rule out that the fibrils have somewhat different chain numbers from the predicted 18.

### 2.5 Xylan content and charge govern fibrillation efficiency and colloidal stability

#### 2.5.1 Xylan content controls nanofibrillation efficiency

Nanofibrillation yields, surface charge, and colloidal stability were assessed to elucidate the physicochemical effects of xylan structure. The nanofibrillation yield was highest for hCNF*_At gux1/2_* (∼80%), followed by hCNF*_Bn_* _WT_ (∼76%) and hCNF*_At_* _WT_ (∼50%) (Figure 5a). The yield correlated strongly with xylan content (Figure S18), suggesting that higher xylan levels facilitate fibril liberation by reducing cellulose–cellulose cohesion.

**Figure 5.**
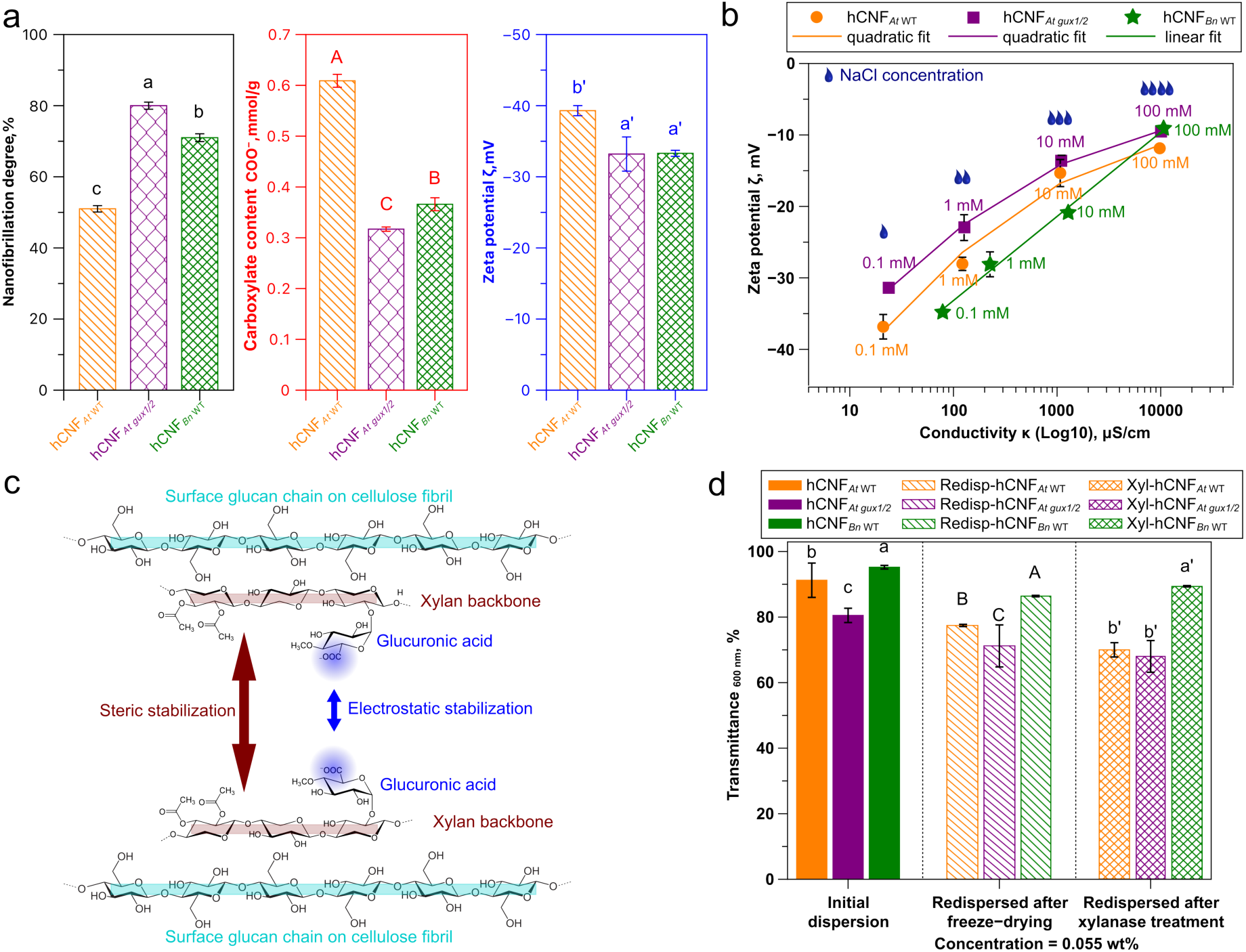
Physical characterization of hCNF colloidal suspensions links nanofibrillation efficiency to xylan content and colloidal stability to xylan charge. a) Degree of nanofibrillation, carboxylate content (conductometric titration), and zeta potential of hCNF colloidal suspensions. Nanofibrillation efficiency was assessed by comparing the dry mass of hCNFs in the supernatant with that in the polydisperse suspension after centrifugation (see separation scheme in Figure S5). b) Zeta-potential (ζ) measurements of hCNFs as a function of increasing NaCl concentration to assess the stabilizing mechanism. c) Schematic illustrating the proposed mechanisms of colloidal stabilization in different hCNF suspensions. d) Light transmittance at 600 nm of freshly prepared and redispersed hCNFs after freeze-drying, and of xylanase-treated hCNF (xyl-hCNF) suspensions. Data are presented as mean ± SD (n = 4) for (a) and (b). For (d), data represent 2–3 formulations with three technical replicates and three measurements per replicate. Mean values not sharing the same letter are significantly different according to Tukey’s test at the 5% significance level.

Carboxylate content did not correlate with fibrillation yield (Figure 5a), indicating that xylan charge is not the primary driver of fibrillation efficiency; rather, structural features such as xylan coverage and matrix porosity appear more influential. The enhanced fibrillation of LC*_At gux1/2_* may be linked to increased cell wall porosity, as evidenced by its greater saccharifiability—an indicator of disrupted lignin–carbohydrate linkages.^[30]^ Together, these findings suggest that nanofibrillation efficiency correlates better with the amount of hCNF xylan than with the xylan charge.

#### 2.5.2 Xylan charge determines colloidal stabilization mechanisms

All hCNF suspensions exhibited zeta potentials (ζ) more negative than –30 mV, indicating substantial surface charge (Figure 5a). This charge, however, arises from more than one source — glucuronidated xylan and residual pectin (present in the *At* samples but not in the *Bn* WT sample) — which complicates interpretation of the static ζ values.

To disentangle electrostatic from steric contributions to colloidal stability, suspensions were titrated with NaCl to progressively screen the surface charge (Figure 5b). For hCNF*_Bn WT_*, ζ decreased approximately linearly with NaCl concentration and colloidal stability was markedly reduced, consistent with predominantly electrostatic stabilization by glucuronidated xylan. In contrast, hCNF*_At_* _WT_ and hCNF*_At gux1/2_* showed a quadratic decrease in ζ and lost stability to a lesser extent, indicating an additional, charge-independent steric contribution — attributable to surface-associated xylan and pectin — that becomes relatively more important where glucuronidation is low or absent. Schematic representations of these proposed mechanisms are shown in Figure 5c.

We next investigated the aggregation characteristics of the hCNF suspensions using UV transmittance assays. hCNFs with glucuronidated xylan exhibited higher transmittance (%T at 600 nm) than hCNFs with non-glucuronidated xylan (Figure 5d). Figure S19a shows the transmittance of hCNFs as a function of concentration, revealing increased flocculation at higher concentrations. Optical transmittance was used as a proxy for redispersibility after freeze-drying and revealed that hCNFs with glucuronidated xylan (*At* WT, *Bn* WT) had good redispersibility and aggregation resistance, losing only 9-15% of their transmittance after freeze-drying (Figure 5d, see Figure S19b for additional transmittance data as a function of suspension concentration). Partial xylan removal had little effect on transmittance (Figures 5d and S19b-c), suggesting that two-fold helical xylan remains bound and maintains interfibrillar spacing. Furthermore, centrifugation and vortexing experiments differentiated interfibrillar behaviors: xylanase-treated *At gux1/2* hCNFs showed pronounced flocculation after centrifugation that resolved only upon vortexing, whereas glucuronidated samples (*At* WT, *Bn* WT) flocculated minimally and remained colloidally stable and visually clear throughout (Figure S20). These results demonstrate that xylan structure — specifically charge — rather than the amount of xylan, governs suspension stability. Freeze-drying had no significant effect on fibril interactions (Figure S19d).

### 2.6 Moisture sorption, thermal stability, and mechanical performance

#### 2.6.1 Water interactions are modulated by xylan charge

Dynamic vapor sorption (DVS) was employed to quantify the moisture sorption behavior of hCNFs. The isotherm profiles of hCNFs (Figure 6a) are typical of systems with strong polymer–polymer and polymer–solvent interactions.^[44]^ Low (a_w_ < 0.2), medium (0.2 ≤ a_w_ ≤ 0.7), and high (a_w_ > 0.7) water-activity regimes were identified in the DVS isotherms (Figure 6a). The hCNF*_At gux1/2_* sample, containing non-glucuronidated xylan, exhibited lower water sorption in the low and medium water-activity (a_w_) regimes than the samples containing glucuronidated xylan (hCNF*_At_* _WT_ and hCNF*_Bn_* _WT_). This behavior indicates reduced accessibility of hydrophilic binding sites and weaker water–polysaccharide interactions, consistent with the absence of glucuronic acid substituents that otherwise promote hydration and swelling. The reduced sorption in these regimes suggests more tightly associated xylan–cellulose interfaces, limiting both strongly bound water uptake and the onset of multilayer adsorption. Consistent with this, Park model fitting of the DVS isotherms yielded a lower K*_d_* value for hCNF*_At gux1/2_* (0.12) than for the samples containing glucuronidated xylan (0.15; Table S5 and Figure S21). Because K*_d_* reflects the affinity and capacity of the polymer matrix for water in the Henry’s law dissolution regime, the reduced K*_d_* confirms diminished water–polysaccharide interactions and fewer effective hydrophilic sorption sites in the non-glucuronidated xylan sample. By analogy, previous studies have shown that enzymatically produced CNFs possess lower surface charge densities than carboxymethylated CNFs, leading to reduced water sorption.^[45]^

**Figure 6.**
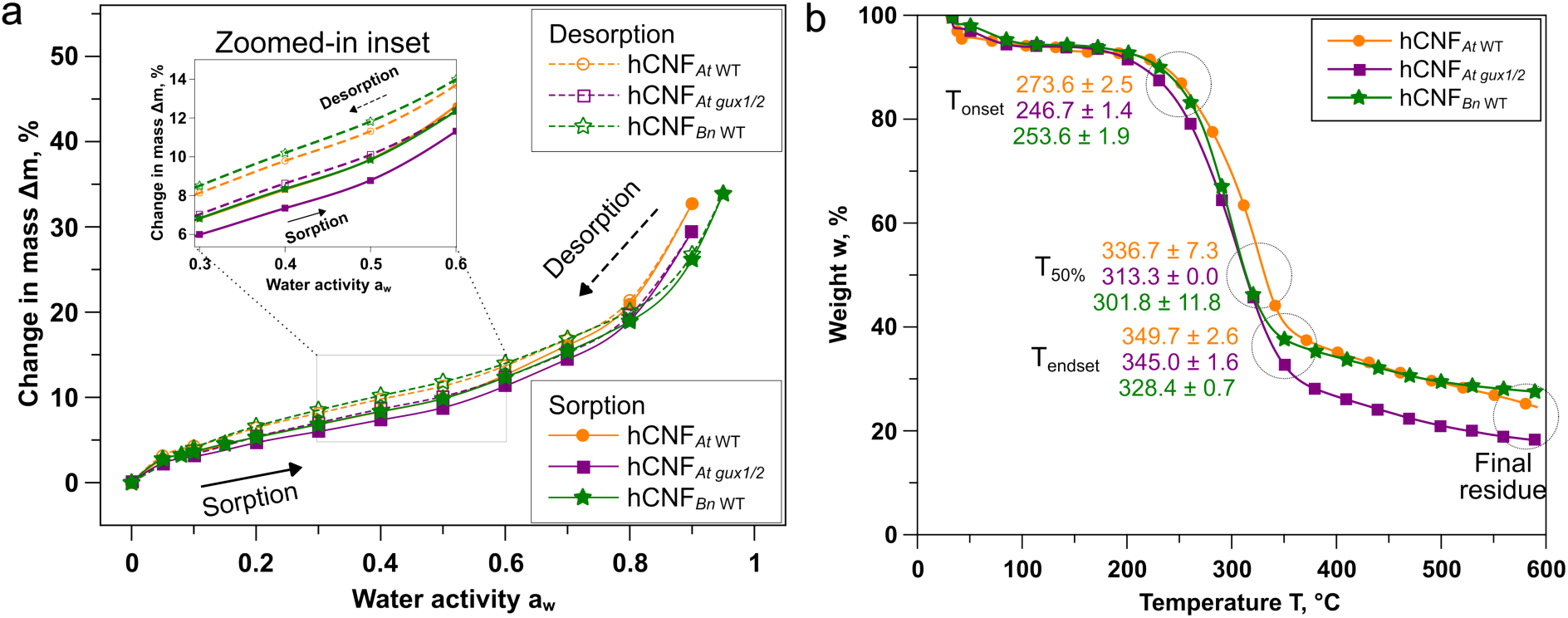
Water–activity–dependent sorption behavior and thermal stability of hCNFs. a) Dynamic vapor sorption (DVS) isotherms of hCNFs, showing characteristic water-uptake regimes at low (a_w_ < 0.2), medium (0.2 ≤ a_w_ ≤ 0.7), and high (a_w_ > 0.7) water activity. The hCNF*_At gux1/2_* sample, which lacked glucuronidated xylan, exhibited reduced water sorption (inset) and a lower K*_d_* value (Table S5 and Figure S21) relative to hCNFs containing glucuronidated xylan, indicating fewer hydrophilic binding sites and weaker water–polysaccharide interactions. b) Representative thermal decomposition profiles of hCNFs, highlighting differences in decomposition temperatures. Decomposition temperatures and statistical comparisons are summarized in Table S6. Data are presented as mean ± SD (n = 3).

#### 2.6.2 Thermal stability reflects polymer composition

The thermal stability of hCNFs was assessed by thermogravimetric analysis (TGA). All samples displayed the characteristic single-step thermal decomposition profile typical of cellulosic materials (Figure 6b).^[46]^ The maximum decomposition temperatures (T_50%_ and T_endset_) followed the order hCNF*_At_* _WT_ > hCNF*_At gux1/2_* > hCNF*_Bn_* _WT_, although the onset decomposition temperature (T_onset_) of hCNF*_At gux1/2_* was slightly lower than that of hCNF*_Bn_* _WT_ (Figure 6b and Table S6). This order broadly correlates with the glucose-to-xylose ratio (Figure S10b). hCNF*_At gux1/2_* yielded the lowest final residue, likely reflecting differences in xylan substitution such as carboxylic acids have been reported to influence residue formation.^[47,48]^ Overall, these results suggest that the thermal stability of hCNFs is broadly correlated with the glucose-to-xylose ratio, whereas the final residue is affected by xylan charge.

#### 2.6.3 Mechanical performance of hCNF films

All hCNF films exhibited high transparency when placed directly on a printed logo (Figure 7a). At a distance of 1.5 cm, hCNF*_Bn_* _WT_ films were the most transparent, followed by hCNF*_At_* _WT_ and hCNF*_At gux1/2_* (Figures 7a and S22), reflecting differences in fibril dispersion, bundling, and interfibrillar interactions.

**Figure 7.**
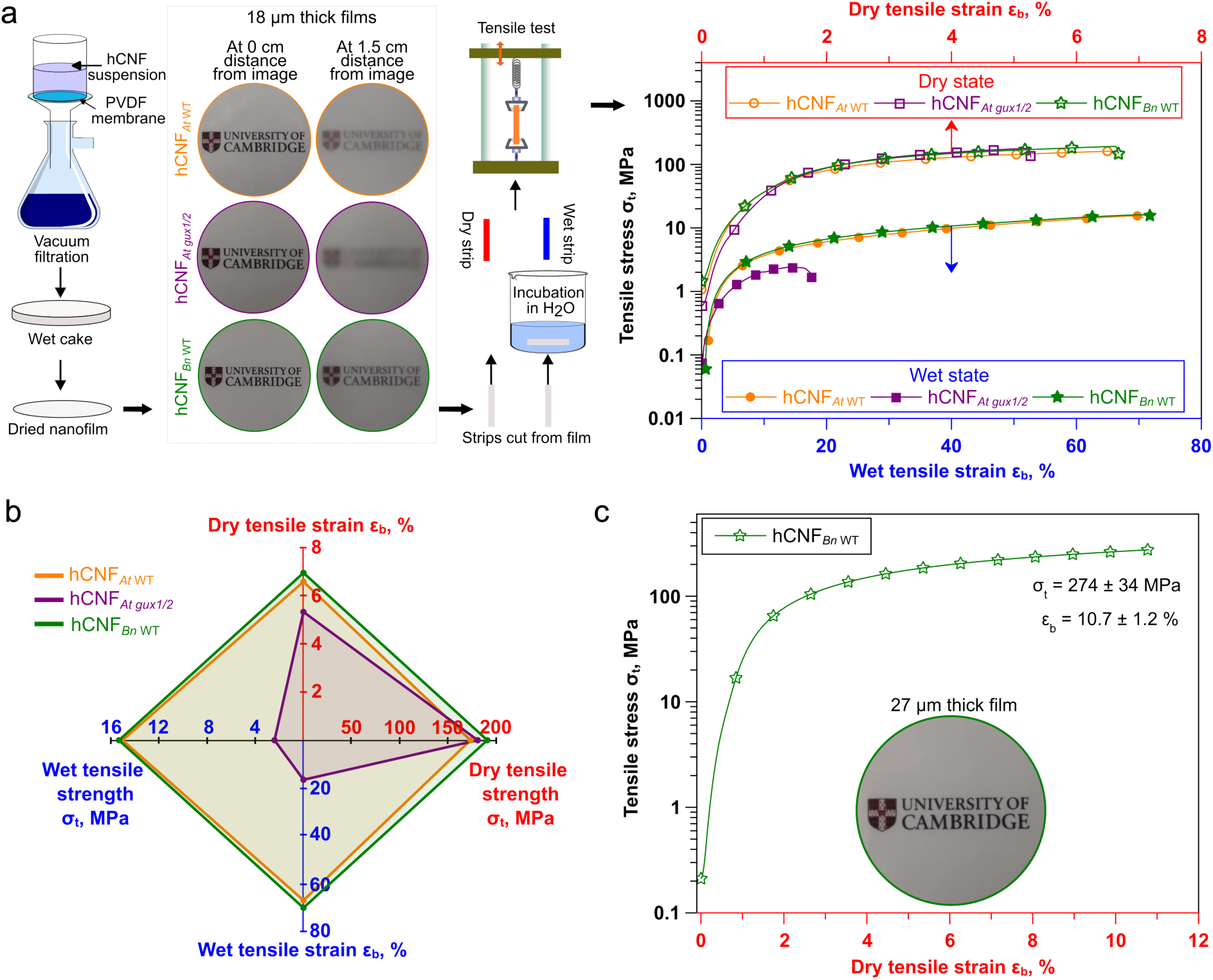
Nanopaper film fabrication, visual transparency, and mechanical properties of hCNF films. a) Schematic illustrating hCNF nanofilm/paper fabrication and tensile testing under dry and wet conditions, together with photographs demonstrating film transparency. Images were captured at 0 and 1.5 cm distance from a printed logo (see additional images in Figure S22). Representative stress–strain curves of hCNF films were measured in the dry state (top x-axis) and wet state (bottom x-axis). Quantitative tensile data are summarized in Table S7 (mean ± SD, n = 8–9). b) Radar plot comparing the tensile properties of hCNF films, revealing that the hCNF*_At gux1/2_* film exhibited significantly reduced stress and strain in the wet state, whereas all films showed comparable behavior in the dry state. c) Representative stress–strain curve of hCNF*_Bn_* _WT_ nanopaper films (27 μm thickness) derived from *Brassica napus* under dry conditions, included to benchmark mechanical performance against previously reported lignocellulosic nanopapers (see Figure S23). Values are presented as mean ± SD (n = 4).

Mechanical testing under dry conditions revealed similar maximum tensile strength (σ_t_) and Young’s modulus (E) across all films (Figure 7a and Table S7). The tensile strain at break (ε_b_) followed the order hCNF*_Bn_* _WT_ (6.9 ± 1.1%) > hCNF*_At_* _WT_ (6.6 ± 1.0%) > hCNF*_At gux1/2_* (5.3 ± 0.6%) (Figure 7b and Table S7). The lower strain observed for hCNF*_At gux1/2_* may be related to its higher xylan content and the absence of glucuronic acid substituents, consistent with previous reports that films composed of short polymers such as xylan tend to be more brittle and less extensible.^[46,49]^

In the wet state, the two films with xylan glucuronidation (from *At* WT and *Bn* WT) retained only about 8–9% of their dry-state tensile strength (σ_t_) (Figures 7a–b and Table S7), consistent with previous observations that pulp-based paper sheets retain less than 10% of their original strength upon wetting.^[50]^ In all films, the tensile strain at break (ε_b_) increased upon wetting, reflecting plasticization by water, although the magnitude of this increase varied: ε_b_ rose roughly tenfold for the two films with xylan glucuronidation but only about threefold for hCNF*_At gux1/2_*. Indeed, hCNF*_At gux1/2_* was a notable exception across all measures, performing markedly worse: it retained only ∼1% of its dry σ_t_, and its wet σ_t_ and ε_b_ were approximately 6-and 4-fold lower, respectively, than those of the other two films, with a correspondingly lower wet Young’s modulus.

This poorer wet performance may reflect weaker interfibrillar cohesion, potentially arising from the absence of glucuronic acid substituents on the xylan, since surface charges in papermaking fibers are known to enhance wet strength.^[51]^ In line with this, atomistic simulations indicate that water can act as a molecular lubricant at cellulose–hemicellulose interfaces, weakening interpolymer interactions upon hydration^[52]^ — a mechanism that would amplify the wet-strength loss seen here, particularly in the charge-deficient hCNF*_At gux1/2_*. In addition, the tendency of these hCNFs to aggregate could alter film porosity. Consistent with this, WAXS analysis of the hCNF*_At gux1/2_* film indicated less dense fibril packing (increased *d*_200_; Figure 3b), which would facilitate water penetration upon wetting.

Taken together, these results show that the influence of xylan structure on hCNF film mechanical properties depends strongly on hydration state. In the dry state, all films behaved similarly, with only modest differences in extensibility that tracked xylan content and the loss of glucuronic acid substituents in hCNF*_At gux1/2_*. Upon wetting, however, these compositional differences became pronounced: although all films lost most of their strength and became more extensible through plasticization, hCNF*_At gux1/2_* was disproportionately weakened, indicating that charged xylan substituents and dense fibril packing are key determinants of wet interfibrillar cohesion. Overall, xylan structure exerts its greatest influence on film mechanical properties most strongly under wet conditions, and retaining native glucuronic acid substitution on xylan therefore appears important for producing hCNF materials with useful wet-state mechanical performance.

#### 2.6.4 Preserving xylan improves mechanical properties in hCNF films

To evaluate whether preservation of the native cellulose and xylan structures leads to improved film performance relative to chemically modified systems, the mechanical properties of hCNF*_Bn_* _WT_ (27 μm thickness) derived from *Brassica napus*, a commercially relevant lignocellulosic source, were compared with those of previously reported CNF films (Figure S23). These thicker hCNF*_Bn_* _WT_ films exhibited a high tensile strength (σ_t_) of 274 ± 34 MPa and a tensile strain at break (ε_b_) of 10.7 ± 1.2% (Figure 7c). The resulting combined stress–strain performance exceeded that reported for nearly all CNF films produced via chemical modification, the sole exception being a film with nematic or anisotropic ordering (data point 13, Figure S23), whose directional alignment confers both high strength and enhanced extensibility. Overall, these results suggest that reassembly of native hCNF building blocks into films can yield mechanical properties that are comparable or superior to those of films fabricated from chemically modified CNFs, highlighting the potential of structurally preserved hCNFs for high-performance, sustainable materials.

## 3 Discussion

The central motivation of this work was to address a fundamental limitation in current cellulose nanofibril (CNF) production strategies: the loss of native cellulose–hemicellulose structure and interactions caused by harsh chemical and mechanical processing.^[10,12]^ Conventional pulping and chemical modification enable effective fibrillation but fundamentally alter the native cellulose–hemicellulose structure,^[7–9]^ disrupting the molecular architecture that supports the remarkable mechanical and functional capabilities of plant cell walls. Consequently, although nanocellulose materials can be manufactured on a large scale, their structures and properties frequently differ markedly from those of natural biomass, and the opportunity to exploit natural biomass variation is thereby lost. This study addresses that challenge by showing that holocellulose nanofibrils (hCNFs) can be isolated while preserving the native cellulose structure, xylan fine structure (substitution and conformation), and cellulose–xylan interactions, thereby enabling a more faithful application of biological design principles in engineered materials. We combined mild delignification, genetic variation in xylan structure, and multiscale characterization to demonstrate that the retained structures and interactions underpin behavior across molecular, colloidal, and macroscopic length scales.

### 3.1 Conservation of native cellulose–xylan structures as a basis for structure–property correlations

Retaining the natural polymer organization throughout processing is essential for establishing meaningful structure–property correlations. The integrated PACE and MAS NMR investigations (Figures 1c and 2a,c) indicate that the glucuronidation and acetylation patterns of xylan, together with the cellulose structure, remained fundamentally unaltered throughout delignification and hCNF isolation. Indeed, we recently extended the approach to poplar hCNFs, describing in detail the cellulose Iβ glucosyl environments.^[27]^ The persistence of both two-fold and three-fold helical xylan conformations (Figures 2c,e) confirms that native cellulose–xylan interactions survived delignification and mechanical fibrillation. This preservation distinguishes the present hCNFs from conventional CNFs generated through one/more pulping and chemical functionalization-treatments, in which hemicellulose removal or chemical alteration markedly modifies fibril surface chemistry and arrangement.^[7–11]^

The ^13^C MAS NMR spectra of the isolated hCNFs were better resolved than those of the intact lignocellulose (Figures 2c–d, Figure S11c). Critically, this resolution gain reflects fractionation rather than structural change: because the cellulose ^13^C chemical shifts were unaltered (Table S4), the sharper spectra report a cleaner population of native fibrils rather than a processing-induced polymorph — direct molecular evidence that isolation selects, rather than reconstructs, native cellulose. The same fractionation that sharpened the spectra should also narrow the fibril population, a prediction borne out by the tight width distributions measured by TEM and cryo-TEM (Figures 4a,c), with no sign of recrystallization or polymorphic transition (Figure 3a,b). Together, these molecular-level observations establish the structural foundation for all subsequent findings and confirm the isolation procedure as one that maintains, rather than disrupts, the native nanofibril architecture.

### 3.2 Correlating molecular structure with fibril morphology and crystalline organization

The nanoscale morphology obtained by TEM and cryo-TEM (Figures 4 a,c) aligns well with the conserved cellulose molecular structure indicated by MAS NMR. The cryo-TEM widths of fibrils in the near-native hydrated state are around 3 nm (Figure 4c) and closely match with the predicted apparent diameters of 18-and 24-chain cellulose Iβ elementary fibrils^[43]^ with different cross-sectional habits (Figure 4d). Beyond experimental error in interpreting low-contrast EM images, the partial xylan coating and the viewing of fibrils from different angles (owing to fibril twisting and interaction with the grid) may account for some of the spread in measured widths. The partial enzymatic removal of accessible xylan produced only a small decrease in fibril width (Figures 4a,c), suggesting that xylan forms a thin, conformationally defined interfacial layer^[3,4]^ rather than a thick, disordered surface coating. A substantial fraction of xylan persisted unhydrolyzed (Figure 4b), likely inaccessible to the enzyme. This residual xylan may consist predominantly of two-fold helical xylan bound to cellulose surfaces, which renders it inaccessible to the xylanase — an enzyme that requires a free xylan backbone for cleavage.^[53]^ Alternatively, steric hindrance may impede xylanase access to xylan within some hCNF bundles (Figure S7). This interpretation is consistent with the MAS NMR detection of surface-bound two-fold helical xylan (Figures 2c,e) and helps explain why fibril widths remained tightly constrained across different plant sources (Figures 4 a,c).

WAXS measurements (Figures 3a,b) further reinforce this picture. Despite differences in xylan content and substitution, all hCNFs exhibited the characteristic cellulose Iβ diffraction pattern with only modest variations in crystallite width and interplanar spacing (Figures 3a,b), in contrast to the substantial increase in crystallite lateral dimensions reported after hydrothermal pretreatment.^[40]^ These small differences are consistent with xylan-mediated modulation of interfibrillar packing rather than disruption of the cellulose crystalline core (Figure 3b). The absence of strong drying effects on crystallite dimensions (Figure 3b) indicates that co-crystallization and irreversible fibril fusion are limited under the processing conditions used, preserving the individuality of the elementary fibrils. Notably, the cellulose Iβ diffraction patterns of the hCNFs reflect contributions from both individual and loosely bundled fibrils, as seen in TEM and AFM images (Figures S7–8).

### 3.3 Distinct roles of xylan content and xylan structure in nanofibrillation and colloidal behavior

Using tailored, genetically modified biomass, we show that nanofibrillation efficiency and colloidal stability reflect the distinct roles of xylan content and xylan structure, respectively. Nanofibrillation efficiency correlated strongly with xylan content, with higher xylan levels facilitating fibril liberation during mechanical processing (Figures 5a and S18). This behavior is consistent with xylan acting as a spacer within the cell wall, reducing cellulose–cellulose cohesion^[5,13–15]^ and enabling more efficient fibrillation. Cell wall porosity may contribute alongside xylan content: LC_At gux1/2_ undergoes markedly more saccharification than the wild type, likely owing to disrupted lignin linkages,^[30,31]^ which could further ease fibril liberation. Xylan charge, by contrast, did not correlate with fibrillation efficiency (Figure 5a), indicating that these structural and architectural factors — rather than electrostatics — govern nanofibrillation.

In contrast, the colloidal stability and interaction behavior of the resulting hCNFs were governed primarily by xylan glucuronidation. Glucuronidated xylan conferred electrostatic stabilization,^[21]^ yielding colloidally stable, redispersible suspensions that resisted aggregation even after freeze-drying (Figures 5a–d and S20). Non-glucuronidated xylan, although promoting fibrillation through its higher content (Figure S18), provided predominantly steric stabilization and was more tightly associated with cellulose surfaces, as evidenced by reduced enzymatic accessibility (Figure 4b) and enhanced aggregation under centrifugation (Figure S20). These contrasting stabilization mechanisms (Figure 5c) are directly traceable to the preserved xylan helical conformations and substitution patterns identified by MAS NMR (Figures 2c,e) and PACE (Figures 1c-ii and S3), respectively, illustrating how subtle structural distinctions shape macroscopic suspension behavior. Together, these results reveal a trade-off between fibrillation efficiency and suspension stability that is finely tuned by the chemical structure of xylan — an insight relevant to the industrial processing and formulation of nanocellulose products.

### 3.4 Water interactions and thermal response reflect preserved cellulose–xylan interactions

Differences in cellulose–xylan interactions also govern macroscopic functional properties, including water sorption and thermal behavior. The reduced moisture uptake of hCNFs containing non-glucuronidated xylan (Figure 6a and Table S5) reflects fewer accessible hydrophilic binding sites and tighter cellulose–xylan association, consistent with the steric stabilization and lower enzymatic accessibility of these samples. In contrast, glucuronidated xylan enhanced hydration capacity, consistent with its electrostatic role in colloidal stability. These hydration tendencies are mirrored in the thermal decomposition behavior: differences in decomposition temperature and char yield tracked the glucose-to-xylose ratio and xylan glucuronidation (Figures 6b and S10b) rather than any alteration in cellulose structure, reinforcing the conclusion that the cellulose Iβ organization is preserved (WAXS data, Figure 3). Collectively, these findings highlight the role of xylan in regulating polymer–water interactions — and, through them, secondary physicochemical properties — while the cellulose structural framework remains intact (MAS NMR data; Figure 2c). In other words, the molecular design of hemicellulose can be used to tune moisture and thermal response without compromising the cellulose structural framework.

### 3.5 Translation of native nanoscale architecture into macroscopic film performance

All the preceding structural and physicochemical differences converge in the mechanical performance of hCNF films. The similar dry stiffness and strength across all films (Figure 7b and Table S7) reflect the conserved cellulose structure and fibril dimensions demonstrated by MAS NMR, WAXS, and TEM/cryo-TEM — the dry response is set by the cellulose structural framework, which is common to all three sources. The differences in extensibility, wet strength, and transparency (Figures 7a–c and Table S7), by contrast, arise from variations in interfibrillar cohesion, packing density, and hydration — properties shaped by the xylan structural features identified in the physicochemical analyses. Films containing glucuronidated xylan showed better wet mechanical performance, traceable to the stronger electrostatic interactions and denser fibril networks that mitigate water-induced weakening; conversely, the poorer wet performance of non-glucuronidated films followed from their lower electrostatic stability, greater tendency to aggregate, and looser packing (WAXS, Figure 3b). The wet state thus discriminates between sources that the dry state does not, precisely because it is governed by the xylan-dependent interfibrillar properties rather than the conserved cellulose core.

This architecture-to-property mapping is what allows native biomass variation to be exploited directly. Thicker films from *Brassica napus* hCNFs exhibited remarkable combined strength and extensibility, surpassing nearly all chemically modified CNF systems reported in the literature,^[18–21,23]^ with the sole exception of an anisotropically ordered film (Figure S23). That a structurally preserved, minimally processed material can match or exceed chemically engineered systems underscores the central benefit of this approach: preserving native cellulose–hemicellulose interactions, rather than compensating for their absence through post-synthetic chemical modification.

## 4 Conclusions

We demonstrate that hemicellulose is not merely an obstacle to processing but an essential structural element that influences nanofibril isolation, interaction, and performance. Xylan content governs the efficiency of hCNF nanofibril liberation, whereas the fine structure of xylan governs interactions among the nanofibrils, their interaction with water, and their behavior under mechanical stress. By maintaining the native cellulose–xylan architecture from isolation through assembly, this study uncovers direct, multiscale structure–property connections that are typically obscured in conventional CNF systems.

Our findings point toward a shift in nanocellulose processing — away from routes that rely on chemical deconstruction followed by re-adsorption of hemicellulose onto isolated cellulose, and toward strategies that preserve and exploit the inherent design of plant cell walls. Combining genetic engineering of hemicellulose architecture with mild extraction offers a robust route to sustainable, high-performance materials that more closely approach the efficiency and function of natural biomass. More broadly, these results establish hemicellulose fine structure as a tunable design lever: because xylan substitution can be altered genetically without disrupting cellulose assembly, properties such as colloidal stability, hydration, and wet-state mechanical performance can be tuned at the level of the living plant, before any processing begins. This reframes the plant cell wall not as feedstock to be deconstructed and rebuilt, but as a biosynthetic platform whose native architecture can be engineered directly — a principle we expect to extend to other cell wall polymers and biomass sources in the design of next-generation bio-based materials.

## Experimental Section

Experimental details are provided in the Supporting information.

## Acknowledgements

This work was supported mainly by the UK Research and Innovation (UKRI) project “Smart Sustainable Plastic Packaging from Plants” (S^2^UPPlant) [grant number NE/V010565/1] and partially by the EPSRC grant “Bioderived and Bioinspired Advanced Materials for Sustainable Industries” (VALUED) [EP/W031019/1]. T. Kuga received support from JSPS KAKENHI (22J12566), and A. Echevarría-Poza acknowledges the Herchel Smith PhD Scholarship from the University of Cambridge. E. Wagner was supported by US Department of Energy grant DE-FG2-84ER13179, and D. J. Cosgrove was supported by NSF Science and Technology Center grant agreement no. CMMI 15-48571. We thank the EPSRC Underpinning Multi-User Equipment Call (EP/P030467/1) and EPSRC/BBSRC (EP/T015063/1; EP/R029946/1 for the 1 GHz instrument) for access to the TEM imaging facility and the UK High-Field Solid-State NMR Facility, respectively. We thank H. Greer for TEM data, and D. Chirgadze, S. Hardwick, and L. Cooper for cryo-TEM data, at Cambridge. WAXS data were collected at the Australian Synchrotron and funded by the New Zealand Ministry of Business, Innovation and Employment [C04X1703] under the Advanced Nanocellulose project’s Strategic Science Investment Fund – Manufactured Products from Trees platform. We thank Nigel Kirby for WAXS data collection at the Australian Synchrotron, and Dinu Iuga and W. Trent Franks for MAS NMR data.

## Author Contributions

PKD and PD conceptualized the project and designed the experiments. RC and RD collected and processed the MAS NMR spectra. PKD analyzed the MAS NMR data in collaboration with YY, RC, SPB, and RD. YY prepared the monosaccharide composition samples and analyzed the data together with PKD. TK acquired the cryo-TEM images. AEP grew ^13^C labeled plants for ssNMR studies, and PH grew *Arabidopsis* plants. MJLG and SD collected the X-ray diffraction data, which were processed by AD; the data were analyzed by PKD, AD, and SD. AFM imaging was performed by AM under the supervision of MV. ND and JB performed the DVS experiments, with data analysis by ND, JB, and PKD. ACSdeA and CGTB conducted computational modeling of cellulose Iβ fibril widths, supervised by MSS. EW performed the wet tensile testing, with data analyzed by PKD, EW, and DJC. JAE was responsible for funding acquisition and project leadership (S^2^UPPlant) and contributed to the analysis of the dry tensile test data. All other data collection and analysis were carried out by PKD. PKD drafted the manuscript. All authors edited, reviewed, and approved the final version of the manuscript.

## References

1. S.-Y. Ding, and M. E. Himmel, “The Maize Primary Cell Wall Microfibril: A New Model Derived from Direct Visualization,” Journal of Agricultural and Food Chemistry 54, no. 3 (2006): 597–606, 10.1021/jf051851z.

2. L. Donaldson, “Cellulose Microfibril Aggregates and Their Size Variation with Cell Wall Type,” Wood Science and Technology 41, no. 5 (2007): 443–460, 10.1007/s00226-006-0121-6.

3. T. J. Simmons, J. C. Mortimer, O. D. Bernardinelli, et al., “Folding of Xylan onto Cellulose Fibrils in Plant Cell Walls Revealed by Solid-State NMR,” Nature Communications 7, no. 1 (2016): 13902, 10.1038/ncomms13902.

4. O. M. Terrett, J. J. Lyczakowski, L. Yu, et al., “Molecular Architecture of Softwood Revealed by Solid-State NMR,” Nature Communications 10, no. 1 (2019), 10.1038/s41467-019-12979-9.

5. M. J. Selig, L. G. Thygesen, C. Felby, and E. R. Master, “Debranching of Soluble Wheat Arabinoxylan Dramatically Enhances Recalcitrant Binding to Cellulose,” Biotechnology Letters 37, no. 3 (2015): 633–641, 10.1007/s10529-014-1705-0.

6. P. K. Deralia, A. K. Sonker, A. Lund, A. Larsson, A. Ström, and G. Westman, “Side Chains Affect the Melt Processing and Stretchability of Arabinoxylan Biomass-Based Thermoplastic Films,” Chemosphere 294 (2022): 133618, 10.1016/j.chemosphere.2022.133618.

7. S. F. Plappert, J.-M. Nedelec, H. Rennhofer, H. C. Lichtenegger, and F. W. Liebner, “Strain Hardening and Pore Size Harmonization by Uniaxial Densification: A Facile Approach toward Superinsulating Aerogels from Nematic Nanofibrillated 2,3-Dicarboxyl Cellulose,” Chemistry of Materials 29, no. 16 (2017): 6630–6641, 10.1021/acs.chemmater.7b00787.

8. A. Isogai, and L. Bergström, “Preparation of Cellulose Nanofibers Using Green and Sustainable Chemistry,” Current Opinion in Green and Sustainable Chemistry 12 (2018): 15–21, 10.1016/j.cogsc.2018.04.008.

9. S. Galland, F. Berthold, K. Prakobna, and L. A. Berglund, “Holocellulose Nanofibers of High Molar Mass and Small Diameter for High-Strength Nanopaper,” Biomacromolecules 16, no. 8 (2015): 2427–2435, 10.1021/acs.biomac.5b00678.

10. T. Saito, S. Kimura, Y. Nishiyama, and A. Isogai, “Cellulose Nanofibers Prepared by TEMPO-Mediated Oxidation of Native Cellulose,” Biomacromolecules 8, no. 8 (2007): 2485–2491, 10.1021/bm0703970.

11. M. Zanuttini, V. Marzocchi, P. Mocchiutti, and M. Inalbon, “Deacetylation Consequences in Pulping Processes,” Holz als Roh - und Werkstoff 63, no. 2 (2005): 149–153, 10.1007/s00107-004-0557-z.

12. M. Beaumont, B. L. Tardy, G. Reyes, et al., “Assembling Native Elementary Cellulose Nanofibrils via a Reversible and Regioselective Surface Functionalization,” Journal of the American Chemical Society 143, no. 41 (2021): 17040–17046, 10.1021/jacs.1c06502.

13. M. A. Kabel, H. van den Borne, J. P. Vincken, A. G. J. Voragen, and H. A. Schols, “Structural Differences of Xylans Affect Their Interaction with Cellulose,” Carbohydrate Polymers 69, no. 1 (2007): 94–105, 10.1016/j.carbpol.2006.09.006.

14. M. A. Kabel, G. Bos, J. Zeevalking, A. G. J. Voragen, and H. A. Schols, “Effect of Pretreatment Severity on Xylan Solubility and Enzymatic Breakdown of the Remaining Cellulose from Wheat Straw,” Bioresource Technology 98, no. 10 (2007): 2034–2042, 10.1016/j.biortech.2006.08.006.

15. P. K. Deralia, A. Jensen, C. Felby, and L. G. Thygesen, “Chemistry of Lignin and Hemicellulose Structures Interacts with Hydrothermal Pretreatment Severity and Affects Cellulose Conversion,” Biotechnology Progress 37, no. 5 (2021): e3189, 10.1002/btpr.3189.

16. S. Iwamoto, K. Abe, and H. Yano, “The Effect of Hemicelluloses on Wood Pulp Nanofibrillation and Nanofiber Network Characteristics,” Biomacromolecules 9, no. 3 (2008): 1022–1026, 10.1021/bm701157n.

17. S. Arola, J. M. Malho, P. Laaksonen, M. Lille, and M. B. Linder, “The Role of Hemicellulose in Nanofibrillated Cellulose Networks,” Soft Matter 9, no. 4 (2013): 1319–1326, 10.1039/c2sm26932e.

18. X. Yang, M. S. Reid, P. Olsén, and L. A. Berglund, “Eco-Friendly Cellulose Nanofibrils Designed by Nature: Effects from Preserving Native State,” ACS Nano 14, no. 1 (2020): 724–735, 10.1021/acsnano.9b07659.

19. F. Chen, W. Xiang, D. Sawada, et al., “Exploring Large Ductility in Cellulose Nanopaper Combining High Toughness and Strength,” ACS Nano 14, no. 9 (2020): 11150–11159, 10.1021/acsnano.0c02302.

20. D. M. de Carvalho, C. Moser, M. E. Lindström, and O. Sevastyanova, “Impact of the Chemical Composition of Cellulosic Materials on the Nanofibrillation Process and Nanopaper Properties,” Industrial Crops and Products 127 (2019): 203–211, 10.1016/j.indcrop.2018.10.052.

21. P. Eronen, M. Österberg, S. Heikkinen, M. Tenkanen, and J. Laine, “Interactions of Structurally Different Hemicelluloses with Nanofibrillar Cellulose,” Carbohydrate Polymers 86, no. 3 (2011): 1281–1290, 10.1016/j.carbpol.2011.06.031.

22. M. J. Selig, L. G. Thygesen, and C. Felby, “Correlating the Ability of Lignocellulosic Polymers to Constrain Water with the Potential to Inhibit Cellulose Saccharification.,” Biotechnology for biofuels 7, no. 1 (2014): 159, 10.1186/s13068-014-0159-x.

23. X. Yang, and L. A. Berglund, “Structural and Ecofriendly Holocellulose Materials from Wood: Microscale Fibers and Nanoscale Fibrils,” Advanced Materials 33, no. 28 (2021): 2001118, 10.1002/adma.202001118.

24. J. C. Mortimer, G. P. Miles, D. M. Brown, et al., “Absence of Branches from Xylan in Arabidopsis Gux Mutants Reveals Potential for Simplification of Lignocellulosic Biomass,” Proceedings of the National Academy of Sciences 107, no. 40 (2010): 17409–17414, 10.1073/pnas.1005456107.

25. I. P. Wood, B. M. Pearson, E. Garcia-Gutierrez, et al., “Carbohydrate Microarrays and Their Use for the Identification of Molecular Markers for Plant Cell Wall Composition,” Proceedings of the National Academy of Sciences 114, no. 26 (2017): 6860–6865, 10.1073/pnas.1619033114.

26. T. Wang, H. Yang, J. D. Kubicki, and M. Hong, “Cellulose Structural Polymorphism in Plant Primary Cell Walls Investigated by High-Field 2D Solid-State NMR Spectroscopy and Density Functional Theory Calculations,” Biomacromolecules 17, no. 6 (2016): 2210–2222, 10.1021/acs.biomac.6b00441.

27. R. Cresswell, P. K. Deralia, Y. Yoshimi, et al., “Using Solid-State NMR to Understand the Structure of Plant Cellulose,” Journal of the American Chemical Society 147, no. 51 (2025): 47223–47236, 10.1021/jacs.5c14452.

28. M. Busse-Wicher, T. C. F. Gomes, T. Tryfona, et al., “The Pattern of Xylan Acetylation Suggests Xylan May Interact with Cellulose Microfibrils as a Twofold Helical Screw in the Secondary Plant Cell Wall of Arabidopsis Thaliana,” Plant Journal 79, no. 3 (2014): 492–506, 10.1111/tpj.12575.

29. S. Koskela, S. Wang, D. Xu, et al., “Lytic Polysaccharide Monooxygenase (LPMO) Mediated Production of Ultra-Fine Cellulose Nanofibres from Delignified Softwood Fibres,” Green Chemistry 21, no. 21 (2019): 5924–5933, 10.1039/C9GC02808K.

30. O. M. Terrett, and P. Dupree, “Lignin-Xylan Interactions in Arabidopsis Stem Affect the Assembly, Porosity and Recalcitrance of the Cell Wall.”

31. O. M. Terrett, and P. Dupree, “Covalent Interactions between Lignin and Hemicelluloses in Plant Secondary Cell Walls,” Current Opinion in Biotechnology 56 (2019): 97–104, 10.1016/j.copbio.2018.10.010.

32. R. Dupree, T. J. Simmons, J. C. Mortimer, et al., “Probing the Molecular Architecture of *Arabidopsis Thaliana* Secondary Cell Walls Using Two- and Three-Dimensional ^13^ C Solid State Nuclear Magnetic Resonance Spectroscopy,” Biochemistry 54, no. 14 (2015): 2335–2345, 10.1021/bi501552k.

33. T. Wang, O. Zabotina, and M. Hong, “Pectin–Cellulose Interactions in the Arabidopsis Primary Cell Wall from Two-Dimensional Magic-Angle-Spinning Solid-State Nuclear Magnetic Resonance,” Biochemistry 51, no. 49 (2012): 9846–9856, 10.1021/bi3015532.

34. Y. Wang, K. Leppänen, S. Andersson, R. Serimaa, H. Ren, and B. Fei, “Studies on the Nanostructure of the Cell Wall of Bamboo Using X-Ray Scattering,” Wood Science and Technology 46, no. 1 (2012): 317–332, 10.1007/s00226-011-0405-3.

35. A. N. Fernandes, L. H. Thomas, C. M. Altaner, et al., “Nanostructure of Cellulose Microfibrils in Spruce Wood,” Proceedings of the National Academy of Sciences 108, no. 47 (2011): E1195–E1203, 10.1073/pnas.1108942108.

36. L. H. Thomas, V. T. Forsyth, A. Martel, I. Grillo, C. M. Altaner, and M. C. Jarvis, “Diffraction Evidence for the Structure of Cellulose Microfibrils in Bamboo, a Model for Grass and Cereal Celluloses,” BMC Plant Biology 15, no. 1 (2015): 153, 10.1186/s12870-015-0538-x.

37. Y. Ono, M. Takeuchi, Y. Zhou, and A. Isogai, “Characterization of Cellulose and TEMPO-Oxidized Celluloses Prepared from Eucalyptus Globulus,” Holzforschung 76, no. 2 (2022): 169–178, 10.1515/hf-2021-0159.

38. Y. Okita, T. Saito, and A. Isogai, “Entire Surface Oxidation of Various Cellulose Microfibrils by TEMPO-Mediated Oxidation,” Biomacromolecules 11, no. 6 (2010): 1696–1700, 10.1021/bm100214b.

39. Y. Nishiyama, P. Langan, and H. Chanzy, “Crystal Structure and Hydrogen-Bonding System in Cellulose Iβ from Synchrotron X-Ray and Neutron Fiber Diffraction,” Journal of the American Chemical Society 124, no. 31 (2002): 9074–9082, 10.1021/ja0257319.

40. C. Driemeier, F. M. Mendes, B. S. Santucci, and M. T. B. Pimenta, “Cellulose Co-Crystallization and Related Phenomena Occurring in Hydrothermal Treatment of Sugarcane Bagasse,” Cellulose 22, no. 4 (2015): 2183–2195, 10.1007/s10570-015-0638-7.

41. T. Oksanen, J. Buchert, and L. Viikari, “The Role of Hemicelluloses in the Hornification of Bleached Kraft Pulps,” Holzforschung 51, no. 4 (1997): 355–360, 10.1515/hfsg.1997.51.4.355.

42. R. Kuramae, T. Saito, and A. Isogai, “TEMPO-Oxidized Cellulose Nanofibrils Prepared from Various Plant Holocelluloses,” Reactive and Functional Polymers 85 (2014): 126–133, 10.1016/j.reactfunctpolym.2014.06.011.

43. D. J. Cosgrove, P. Dupree, E. D. Gomez, C. H. Haigler, J. D. Kubicki, and J. Zimmer, “How Many Glucan Chains Form Plant Cellulose Microfibrils? A Mini Review,” Biomacromolecules 25, no. 10 (2024): 6357–6366, 10.1021/acs.biomac.4c00995.

44. R. Kohler, R. Alex, R. Brielmann, and B. Ausperger, “A New Kinetic Model for Water Sorption Isotherms of Cellulosic Materials,” Macromolecular Symposia 244, no. 1 (2006): 89–96, 10.1002/masy.200651208.

45. E. Brännvall, and C. Aulin, “CNFs from Softwood Pulp Fibers Containing Hemicellulose and Lignin,” Cellulose 29, no. 9 (2022): 4961–4976, 10.1007/s10570-022-04585-8.

46. P. K. Deralia, A. M. Du Poset, A. Lund, A. Larsson, A. Strom, and G. Westman, “Oxidation Level and Glycidyl Ether Structure Determine Thermal Processability and Thermomechanical Properties of Arabinoxylan-Derived Thermoplastics,” ACS Applied Bio Materials 4, no. 4 (2021): 3133–3144, 10.1021/acsabm.0c01550.

47. K. Lichtenstein, and N. Lavoine, “Toward a Deeper Understanding of the Thermal Degradation Mechanism of Nanocellulose,” Polymer Degradation and Stability 146 (2017): 53–60, 10.1016/j.polymdegradstab.2017.09.018.

48. H. Fukuzumi, T. Saito, Y. Okita, and A. Isogai, “Thermal Stabilization of TEMPO-Oxidized Cellulose,” Polymer Degradation and Stability 95, no. 9 (2010): 1502–1508, 10.1016/j.polymdegradstab.2010.06.015.

49. P. K. Deralia, A. M. du Poset, A. Lund, A. Larsson, A. Ström, and G. Westman, “Hydrophobization of Arabinoxylan with N-Butyl Glycidyl Ether Yields Stretchable Thermoplastic Materials,” International Journal of Biological Macromolecules 188 (2021): 491–500, 10.1016/j.ijbiomac.2021.08.041.

50. H. H. Espy, “The Mechanism of Wet-Strength Development in Paper: A Review,” 78, no. 4 (1995).

51. J. Laine, T. Lindström, G. G. Nordmark, and G. Risinger, “Studies on Topochemical Modification of Cellulosic Fibres: Part 3. The Effect of Carboxymethyl Cellulose Attachment on Wet-Strength Development by Alkaline-Curing Polyamide-Amine Epichlorohydrin Resins,” Nordic Pulp & Paper Research Journal 17, no. 1 (2002): 57–60, 10.3183/npprj-2002-17-01-p057-060.

52. L. N. Trentin, A. C. S. Alcântara, C. G. T. Batista, and M. S. Skaf, “Unraveling the Mechanical Behavior of Softwood Secondary Cell Walls through Atomistic Simulations,” Biomacromolecules 26, no. 6 (2025): 3395–3409, 10.1021/acs.biomac.4c01806.

53. H. S. Overkleeft, G. J. Davies, and S. J. Williams, “Catalyzing Carbohydrate Cleavage: Glycosidases and Their Mechanisms,” Chemical Reviews 126, no. 5 (2026): 3287–3323, 10.1021/acs.chemrev.5c00803.

