## Supplemental file for "Preserving Native Cellulose–Xylan Architecture Enables Structure–Property Control in Holocellulose Nanofibrils and High-Performance Sustainable Materials"

Y. Yoshimi
[$] Now at Department of Science and Technology on Food Safety, Faculty of Biology-Oriented Science and Technology, Kindai University, 930 Nishimitani, Kinokawa, Wakayama, 649-64693, Japan

T. Kuga
Department of Biomaterials Sciences, Graduate School of Agricultural Life Sciences, University of Tokyo, 1-1-1 Yayoi, Bunkyo, Tokyo, 113-8657, Japan

R. Cresswell, S.P. Brown, R. Dupree
Department of Physics, University of Warwick, Coventry, CV4 7AL, United Kingdom

P. Howell
Niab, Park Farm, Villa Road, Histon, Cambridge CB24 9NZ, United Kingdom

A. Dickson, M.J. Le Guen, S. J. Hill
New Zealand Institute for Bioeconomy Science Limited, Private Bag 3020, Rotorua, New Zealand

E. Wagner, D. J. Cosgrove
Department of Biology, Pennsylvania State University, University Park, PA 16802, USA

A.C.S. de Alcântara, C.G.T. Batista, M.S. Skaf
Institute of Chemistry, University of Campinas, São Paulo 13083-861, Brazil

N. Follain
Univ Rouen Normandie, INSA Rouen Normandie, CNRS, Normandie Univ, PBS UMR 6270, F-76000 Rouen, France

A. Miller, M. Vendruscolo
Yusuf Hamied Department of Chemistry, University of Cambridge, Lensfield Road, Cambridge CB2 1EW, United Kingdom

J. Beaugrand
UR1268 BIA, INRAE Nantes, France

J. A. Elliott
Department of Materials Science & Metallurgy, University of Cambridge, 27 Charles Babbage Road, Cambridge, CB3 0FS, United Kingdom

### Deceased on 17 June 2025

### 1. Chemicals, enzymes, materials, sample nomenclature, and acronyms

#### *1.1 Chemicals, enzymes, and materials*

All chemicals were purchased from Sigma-Aldrich and used as received unless stated otherwise. Deionized (DI) water was used throughout, including for the preparing aqueous suspensions.

GH10 endo-β-1,4-xylanase (*Cj*GH10) from *Cellvibrio japonicus* was provided by Novozymes (Denmark). GH115A α-glucuronidase (*Bo*GH115A) from *Bacteroides ovatus* and CE4 acetyl xylan esterase 4A (*Ct*Axe4A) from *Clostridium thermocellum* were purchased from NZYtech (Portugal).

#### *1.2 Lignocelluloses*

*Arabidopsis thaliana* (Col-0 ecotype, wild type; *At* WT) and the GUX1/GUX2-deficient mutant lacking glucuronic acid (GlcA) substitutions on xylan (*At gux1/2*) were grown at Niab Park Farm (Cambridge, UK) and harvested at 11 weeks. Plants were stored undried at −20 °C. Only the bottom stem segments, which are enriched in secondary cell walls, were used. Chlorophyll was removed prior to use by incubation in 96% (v/v) ethanol at 70 °C for 30 min, followed by washing with DI water.

*Brassica napus* (wild type; *Bn* WT), known as oilseed rape straw, was provided by Professor James Elliott (University of Cambridge, UK) as dried bales sourced locally. The material was washed with water and cut into smaller pieces with scissor prior to use.

#### *1.3 Information on lignocellulose, sample nomenclature, acronyms*

**Table S1.** Lignocellulosic biomass samples used in this study.

| Serial number | Lignocellulose plant | Acronym | Growth information | Remark |
|---|---|---|---|---|
| *1* | *Arabidopsis* | *At* | - | Hardwood model plant, only bottom stems used |
| *1.1* | *Arabidopsis thaliana* wild type | *At* WT | Green house grown | Plant having xylan with 4-*O*-methyl-D-glucuronic acid (MeGlcA) |
| *1.2* | *Arabidopsis thaliana gux1/2* | *At gux1/2* | Green house grown | Plant having xylan without 4-*O*-methyl-D-glucuronic acid ([Me]GlcA), [Me]- methylated glucuronic acid and non-methylated glucuronic acid together |
| *2* | *Brassica napus* | *Bn* | - | Common name- rapeseed straw |
| *2.1* | *Brassica napus* wild type | *Bn* WT | Field grown | Commercial-relevant lignocellulose; used to benchmark *Arabidopsis* |

**Table S2.** Sample nomenclature used in this study

| | | Sample | Acronym | Remark |
|---|---|---|---|---|
| 1 | | Lignocellulose | LC | Plant biomass |
| | 1.1 | *Arabidopsis thaliana* wild type | $LC_{\mathit{At}\ \mathrm{WT}}$ | |
| | 1.2 | *Arabidopsis thaliana gux1/2* | $LC_{\mathit{At\ gux1/2}}$ | |
| | 1.3 | *Brassica napus* wild type | $LC_{\mathit{Bn}\ \mathrm{WT}}$ | |
| 2 | | Holocellulose | hC | Peracetic acid delignified lignocellulose |
| | 2.1 | *Arabidopsis thaliana* wild type holocellulose | $hC_{\mathit{At}\ \mathrm{WT}}$ | |
| | 2.2 | *Arabidopsis thaliana gux1/2* holocellulose | $hC_{\mathit{At\ gux1/2}}$ | |
| | 2.3 | *Brassica napus* wild type holocellulose | $hC_{\mathit{Bn}\ \mathrm{WT}}$ | |
| 3 | | Holocellulose nanofibrils | hCNF | Fibrils separated from blended hC suspension; Hemicellulose-preserved CNFs are referred to as holocellulose nanofibrils (hCNFs |
| | 3.1 | *Arabidopsis thaliana* wild type holocellulose nanofibrils | $hCNF_{\mathit{At}\ \mathrm{WT}}$ | |
| | 3.2 | *Arabidopsis thaliana gux1/2* holocellulose nanofibrils | $hCNF_{\mathit{At\ gux1/2}}$ | |
| | 3.3 | *Brassica napus* wild type holocellulose nanofibrils | $hCNF_{\mathit{Bn}\ \mathrm{WT}}$ | |
| 4 | | Xylanase treated holocellulose nanofibrils | Xyl-hCNF | Fibrils made from undigested hCNFs after xylanase-cocktail treatment |
| | 4.1 | Xylanase treated holocellulose nanofibrils from *Arabidopsis thaliana* wild type | Xyl-$hCNF_{\mathit{At}\ \mathrm{WT}}$ | |
| | 4.2 | Xylanase treated holocellulose nanofibrils from *Arabidopsis thaliana gux1/2* | Xyl-$hCNF_{\mathit{At\ gux1/2}}$ | |
| | 4.3 | Xylanase treated holocellulose nanofibrils from *Brassica napus* wild type | Xyl-$hCNF_{\mathit{Bn}\ \mathrm{WT}}$ | |

**Table S3.** List of acronyms and symbols

| Acronym | Definition |
|---|---|
| AFM | Atomic force microscopy |
| *At* | *Arabidopsis thaliana* |
| At *gux1/2* | *Arabidopsis thaliana* xylan glucuronidation-deficient mutant |
| *At* WT | *Arabidopsis thaliana* wild type |
| $a_w$ | Water activity |
| *Bn* | *Brassica napus* |
| *Bn* WT | *Brassica napus* wild type |
| $C4^{D1}/C6^{D1}$ | Cellulose glucosyl of domain 1 of carbon4/6 at ~89 ppm and ~65 ppm |
| $C4^{D2}/C6^{D2}$ | Cellulose glucosyl of domain 2 of carbon4/6 at ~84 ppm and ~62 ppm |
| CE4 | Carbohydrate esterase family 4 (acetyl esterase) |
| CNF | Cellulose nanofibril, used interchangeably with cellulose microfibril |
| CP | Cross-polarization |
| Cryo-TEM | Cryogenic transmission electron microscopy |
| D1 | Spectral domain 1 ($C4^{D1}/C6^{D1}$ $^{13}C$ chemical shifts of ~89 ppm and ~65 ppm) |
| D2 | Spectral domain 2 ($C4^{D2}/C6^{D2}$ $^{13}C$ chemical shifts of ~84 ppm and ~62 ppm) |
| domain 1 / domain 2 | Ratio of distinct cellulose glucosyl environments resolved at C4 and C6 in MAS NMR |
| $d_{200}$ | Interplanar (lattice) spacing of the 200 reflection |
| DMSO | Dimethyl sulfoxide |
| DQ | Double quantum |
| DVS | Dynamic vapor sorption |
| E | Young's modulus |
| GH10 | Glycoside hydrolase family 10 (xylanase) |
| GH115 | Glycoside hydrolase family 115 (α-glucuronidase) |
| GlcA | Glucuronic acid |
| *gux* | GlucUronic acid substitution of Xylan (mutant) |
| *gux1/2* | gux double mutant (glucuronic acid unsubstituted xylan) |
| hC | Holocellulose (delignified lignocellulose) |
| hCNF | Holocellulose nanofibril (Hemicellulose-preserved CNFs are referred to as holocellulose nanofibrils (hCNFs) |
| INADEQUATE | Incredible natural-abundance double-quantum transfer experiment |
| $K_d$ | Park-model dissolution (Henry's-law) coefficient |
| $L_{200}$ | Crystallite width derived from the 200 reflection |

| | |
|---|---|
| LC | Lignocellulose / lignocellulosic |
| MAS | Magic-angle spinning |
| [Me]GlcA | [4-*O*-methyl]-D-glucuronic acid, [Me] denotes both methylated and non-methylated glucuronic acid |
| NMR | Nuclear magnetic resonance |
| PACE | Polysaccharide analysis by carbohydrate gel electrophoresis |
| PDSD | Proton-driven spin diffusion |
| SQ | Single quantum |
| ssNMR | Solid-state nuclear magnetic resonance |
| TEM | Transmission electron microscopy |
| TEMPO | (2,2,6,6-Tetramethylpiperidin-1-yl)oxyl |
| $T_{endset}$ | Endset decomposition temperature |
| TGA | Thermogravimetric analysis |
| $T_{onset}$ | Onset decomposition temperature |
| $T_{50\%}$ | Temperature at 50% mass loss |
| $UX_3$ | Glucuronosyl-substituted xylotriose digestion product |
| WAXS | Wide-angle X-ray scattering |
| $Xn4^{2f}$ / $Xn4^{3f}$ | Xylan C4 carbon in two-fold / three-fold helical conformation |
| $Xn5^{2f}$ / $Xn5^{3f}$ | Xylan C5 carbon in two-fold / three-fold helical conformation |
| xyl-hCNF | Xylanase-treated holocellulose nanofibril |
| $\zeta$ | Zeta potential |
| $\sigma_t$ | Maximum tensile strength |
| $\varepsilon_b$ | Tensile strain at break |

#### 2. Holocellulose nanofibril (hCNF) isolation and xylan structural characterization

##### *2.1 Monosaccharide and lignin content and xylan substitution pattern*

###### 2.1.1 Preparation of alcohol insoluble residue (AIR)

Alcohol-insoluble residues (AIR) from the different lignocellulosic materials, used for PACE analysis and for determination of monosaccharide compositional and lignin content, were prepared as described previously.[1] Plant material was incubated in 96% (v/v) ethanol at 70 °C for 30 min and homogenized using a ball mill (MM400, Retsch). The homogenate was centrifuged at 4000 g for 15 min, and the resulting solid residue was washed sequentially with

100% (v/v) ethanol; chloroform:methanol (2:3, v/v; twice); and 65%, 80%, and 100% (v/v) ethanol.

2.1.2 Monosaccharide compositional analysis

Trifluoroacetic acid (TFA) hydrolysis was used to determine the composition of non-cellulosic polysaccharides and was performed as described previously.[1,2] Briefly, alcohol-insoluble residue (AIR; 1 mg) was incubated in 2 M TFA (400 μL) at 121 °C for 1 h. After cooling to room temperature, the samples were centrifuged, and the supernatant was dried *in vacuo* and retained for subsequent HPLC analysis.

The pellet remaining after TFA hydrolysis was subjected to a two-stage sulfuric acid hydrolysis to quantify glucose derived from cellulose and xylose from the TFA-resistant xylan fraction. The residue was first hydrolyzed with 72% (w/w) sulfuric acid at 24 °C for 1 h. The hydrolysate was then diluted to 1 M sulfuric acid and heated at 100 °C for 3 h. After hydrolysis, the samples were neutralized with barium carbonate and prepared for HPLC analysis.

Total cell-wall monosaccharides were quantified by HPAEC–PAD following established protocols.[1] Solubilized neutral monosaccharides were quantified using a Dionex ICS3000 system equipped with a PA20 analytical column, a PA20 guard column, and a borate trap (Dionex, Surrey, UK), as described previously.[1,3]

Acidic monosaccharides, including galacturonic acid (GalA) and glucuronic acid (GlcA), were separated using a linear gradient of 20–200 mM ammonium acetate in 100 mM NaOH over 10 min, followed by a 10-min isocratic step at 200 mM ammonium acetate in 100 mM NaOH.[3]

2.1.3 Lignin content measurement

Total lignin content in LC, hC, hCNFs: Total lignin content, including both acid-insoluble and acid-soluble fractions, was determined using a spectroscopic method following a previously reported procedure.[4] Alcohol-insoluble residue (AIR; 1 mg) was placed in a screw-cap microcentrifuge tube, and 1 mL of freshly prepared cysteine solution (0.1 g mL$^{-1}$ in 72% sulfuric acid) was added. The samples were incubated in a thermomixer at 24 °C and 600 rpm, with vortex mixing every 10 min. After complete dissolution (1 h), the reaction mixture was diluted to a final volume of 50 mL with deionized water.

UV–vis absorbance of the diluted solution was recorded between 200 and 400 nm using a Shimadzu UV–vis spectrophotometer (UV-1800). Measurements were performed in quartz cuvettes with a 1 cm path length. Hydrolysis was carried out in triplicate, and three UV measurements were collected for each sample. Data are reported as the mean ± standard deviation.

Total lignin content was calculated according to the following equation:

$$Total\ lignin\ content, \% = \frac{abs \times V}{\varepsilon \times m \times L} \times 100 \quad (1)$$

where *abs* is the UV absorbance at $\lambda = 283$ nm, $V$ is the total volume of the diluted solution (L), $m$ is the dry mass of the sample (g), $L$ is the cuvette path length (cm), and $\varepsilon$ is the lignin absorption coefficient at 283 nm (g$^{-1}$ L cm$^{-1}$).

Klason (acid-insoluble) lignin content was determined using a previously reported protocol.[5]

##### 2.1.4 Extraction of acetylated xylan from lignocellulose, holocellulose, and holocellulose nanofibrils from *Brassica napus* (*Bn* WT)

Alcohol-insoluble residue (AIR) from *Brassica napus* wild-type lignocellulose ($LC_{\mathit{Bn}\ \mathrm{WT}}$) was delignified using 11 wt% peracetic acid at 85 °C for 30 min. Acetylated xylan was then extracted from the delignified lignocellulose ($LC_{\mathit{Bn}\ \mathrm{WT}}$), holocellulose ($hC_{\mathit{Bn}\ \mathrm{WT}}$), and holocellulose nanofibrils ($hCNF_{\mathit{Bn}\ \mathrm{WT}}$), by treatment with dimethyl sulfoxide (DMSO) at 60 °C for 24 h. The extraction was repeated three times; after each 24 h cycle, the DMSO extract was collected by centrifugation and replaced with fresh DMSO.

Acetylated xylan was precipitated from the combined DMSO extracts by adding four volumes of ethanol (EtOH), followed by acidification with formic acid to pH 3.5. The mixture was incubated at 4 °C for 48 h to ensure complete precipitation and then centrifuged. The precipitate was washed with excess methanol (five volumes relative to the DMSO extract), recovered by centrifugation, resuspended in deionized water, and dried using a centrifugal evaporator at 30 °C.

##### 2.1.5 Xylan substitution analysis by Polysaccharide Analysis by Carbohydrate gel Electrophoresis (PACE)

Xylan substitution patterns, specifically the presence of methylglucuronic acid (MeGlcA) substituents, were analyzed by polysaccharide analysis using carbohydrate gel electrophoresis (PACE) of xylanase-generated oligosaccharides.

To obtain non-acetylated xylan, alcohol-insoluble residue (AIR; 1 mg) was treated with 20 µL of 4 M NaOH for 1 h and neutralized with 1 M HCl. Both non-acetylated and acetylated xylan samples were then incubated in 0.1 M ammonium acetate buffer (pH 6.0) with an excess of GH10 xylanase and digested overnight at 21 °C to ensure complete hydrolysis. Control reactions lacking either substrate or enzyme were performed in parallel to identify any nonspecific bands.

The released oligosaccharides were dried and derivatized with 8-aminonaphthalene-1,3,6-trisulfonic acid (ANTS; Invitrogen). ANTS derivatization, electrophoretic separation, and visualization were carried out as described previously.[2,6]

2.1.6 Supporting data for neutral monosaccharide and lignin of lignocelluloses

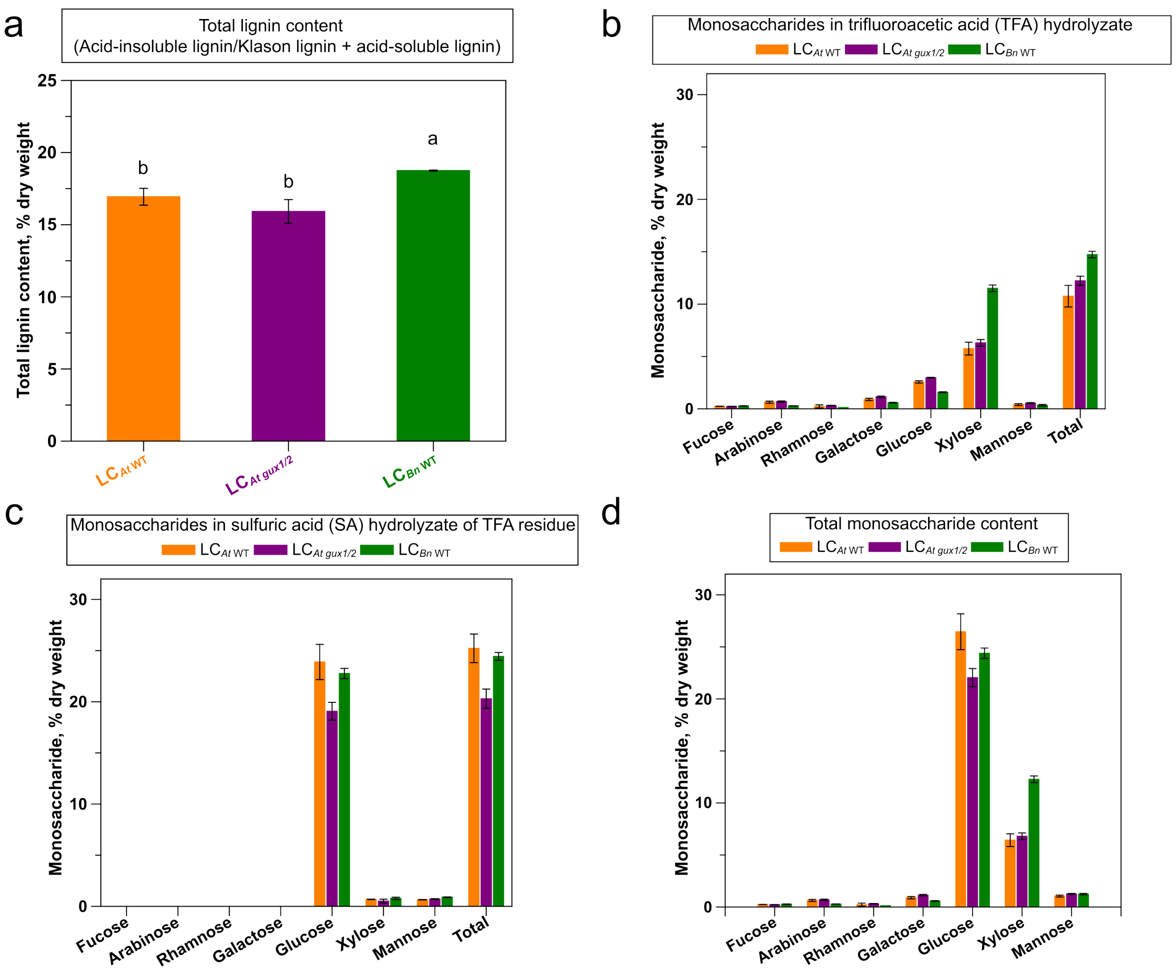

**Figure S1.** Compositional analysis of lignocellulosic substrates. a) Total lignin content in different lignocelluloses, determined by UV–vis spectroscopy following cysteine–sulfuric acid hydrolysis.[4] Mean values not sharing the same letter are significantly different according to Tukey's test at the 5% significance level. Corresponding $p$ values are: LC$_{At\ \mathrm{WT}}$ vs LC$_{At\ gux1/2}$ =0.1619; LC$_{At\ \mathrm{WT}}$ vs LC$_{Bn\ \mathrm{WT}}$ =0.0209; LC$_{At\ gux1/2}$ vs LC$_{Bn\ \mathrm{WT}}$ =0.0025. b) Neutral monosaccharide composition of hemicelluloses released by trifluoroacetic acid (TFA) hydrolysis. c) Neutral monosaccharide composition of the TFA-insoluble residue, determined after two-step sulfuric acid hydrolysis.[5] d) Total monosaccharide content of the lignocellulosic samples. The glucose-to-xylose ratio are 3.5, 3.2 and 1.82 in LC$_{At\ \mathrm{WT}}$, LC$_{At\ gux1/2}$, and LC$_{Bn\ \mathrm{WT}}$. Data are presented as mean ± SD (n = 3).

2.1.7 Supporting data for PACE gels

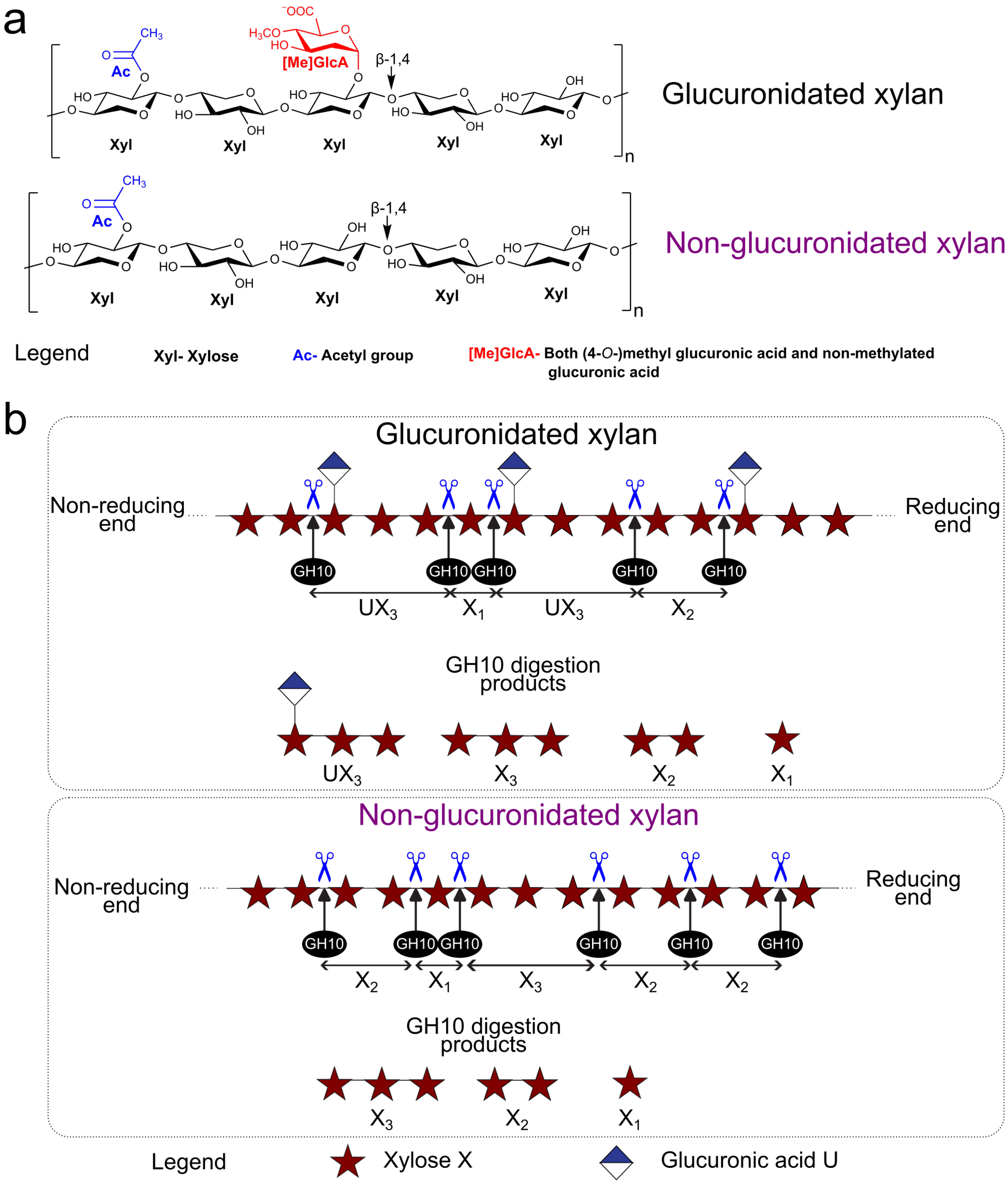

**Figure S2.** Chemical structure of glucuronidated and non-glucuronidated xylan and their digestion by *Cj*GH10 xylanase. a) Chemical structure of glucuronidated and non-glucuronidated xylan, each consisting of a β-1,4-linked D-xylosyl backbone bearing *O*-acetyl substituents, with (glucuronidated) or without (non-glucuronidated) 4-*O*-methyl-D-glucuronic acid ([Me]GlcA) side groups. b) Action of *Cj*GH10 xylanase on non-acetylated xylan and the resulting oligosaccharides. Schematic illustration of GH10 xylanase cleavage patterns on two xylan substrates. Glucuronidated xylan yields $X_1$, $X_2$, $X_3$, and the diagnostic glucuronidated oligosaccharide $UX_3$, whereas non-glucuronidated xylan is hydrolyzed to $X_1$, $X_2$, and $X_3$ only.

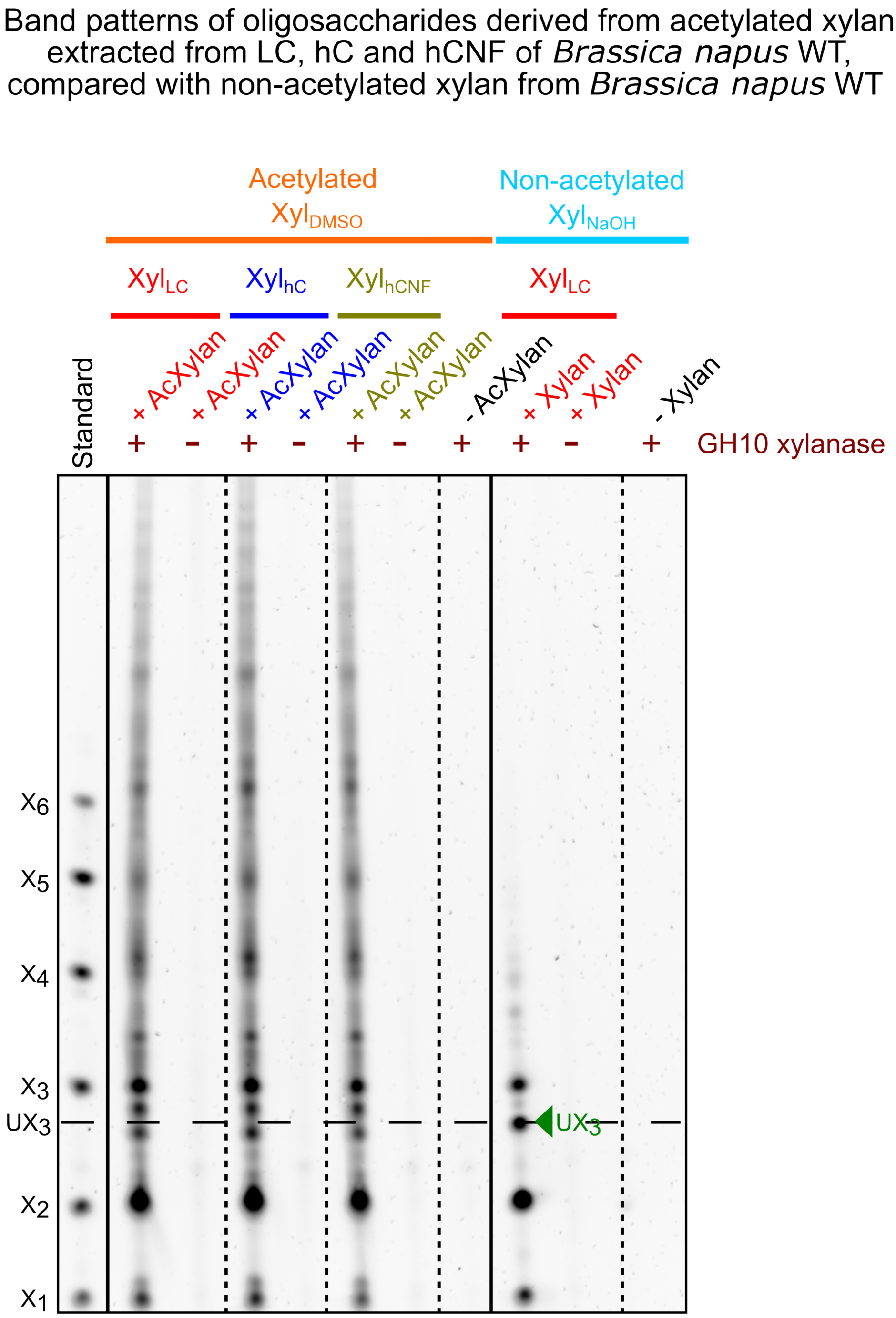

**Figure S3.** PACE analysis of GH10 xylanase digestion products from acetylated and non-acetylated xylan. PACE gels highlight distinct oligosaccharide fingerprints following GH10 xylanase treatment. Acetylated xylan is only partially hydrolyzed, owing to steric hindrance from acetyl substituents that limits enzyme access, whereas non-acetylated xylan is efficiently digested, yielding predominantly short oligosaccharides ($X_1$, $X_2$, $X_3$, and $UX_3$; green arrowhead). $X_1$–$X_6$ denote xylosyl oligosaccharide standards.

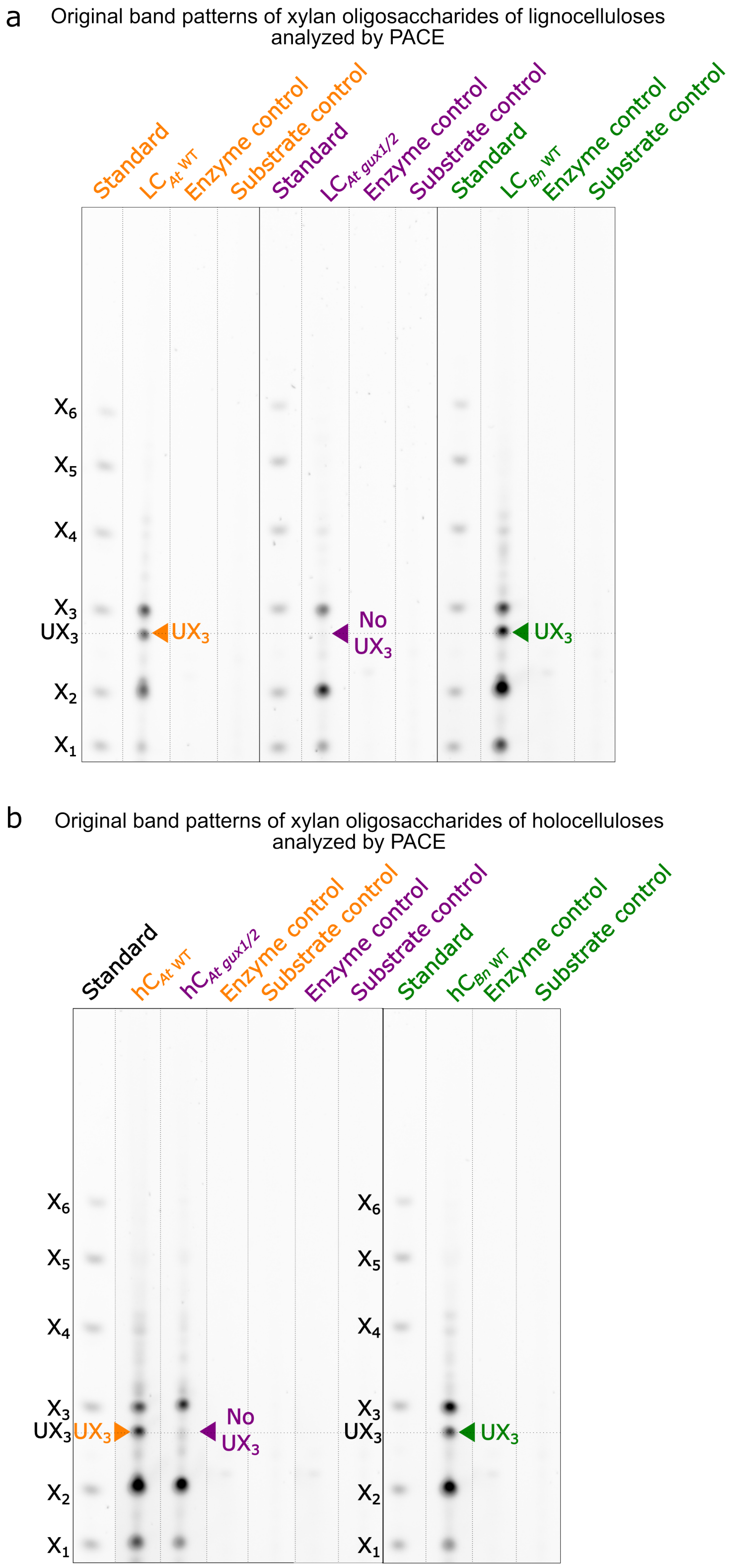

**Figure S4.** Polysaccharide analysis by carbohydrate gel electrophoresis (PACE) of GH10 xylanase digestion products, with controls. Original PACE gel corresponding to Figure 1c in the main text, shown together with substrate-only and enzyme-only controls. PACE gels of GH10 xylanase digestion products from lignocellulose (LC) and holocellulose (hC) was used to assess xylan glucuronidation. i) LC and ii) hC from *Arabidopsis thaliana* wild type (*At* WT) and *Brassica napus* wild type (*Bn* WT) display glucuronidated $UX_3$ digestion products (orange and green arrowheads), whereas the *Arabidopsis thaliana* mutant (*At gux1/2*) lacks $UX_3$ (red arrowhead). $X_1$–$X_6$ denote xylosyl oligosaccharide standards.

*2.2 Holocellulose preparation, nanofibrillation, and microscopy*

2.2.1 Holocellulose (hC) preparation

Holocelluloses (hCs) were prepared as described previously,[7] with minor modifications. Briefly, lignocellulosic materials were delignified using 3 wt% peracetic acid (PAA; 0.32–0.35 g pure PAA $g^{-1}$ dry material, pH 4.8) at 85 °C for 45 min without stirring. The delignification was repeated for three further cycles, with decantation of the spent PAA, one intermediate water wash between cycles, and replenishment with fresh 3 wt% PAA. After the final treatment, the delignified material was washed thoroughly with deionized water until the conductivity of the wash water fell below 10 µS $cm^{-1}$. The resulting holocellulose was then blended for 2 min to obtain a homogeneous slurry and stored at 4 °C prior to further processing.

Dilute holocellulose (hC) dispersions (0.10–0.12 wt%) were mechanically fibrillated using a Vitamix A3500i blender for a total of 30 min, applied as fifteen 2 min blending cycles with interim storage at 4 °C for at least 30 min between cycles to minimize sample heating. The resulting polydisperse suspension was centrifuged at 5000 rpm (4470 g) for 15 min to separate the fibrillated fraction. The supernatant, containing holocellulose nanofibrils (hCNFs), was collected for subsequent analyses (see schematic in Figure S5).

2.2.2 Holocellulose (hC) fibrillation

The unfibrillated material recovered as the centrifugation pellet is hereafter referred to as the sediment (see Figure S5). The polydisperse suspension, hCNF supernatant, and sediment fractions were stored at 4 °C prior to further characterization. Photographs of the hCNF suspensions are presented in Figure S6.

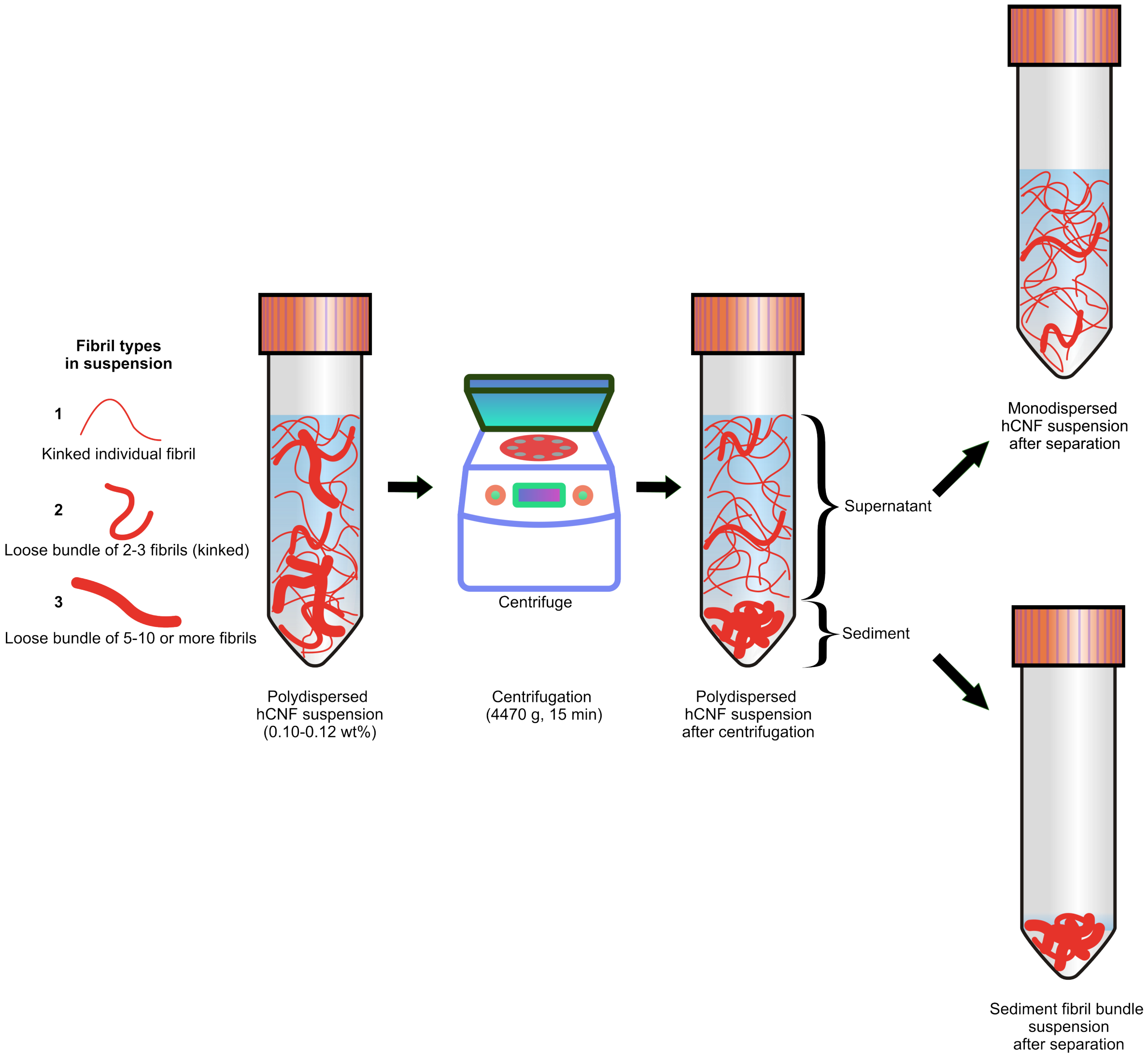

**Figure S5.** Schematic illustration of the fractionation of blended holocellulose suspensions. A simplified scheme depicting the separation of fibrillated, colloidally stable nanofibrils (hCNFs) from unfibrillated, colloidally unstable fractions (sediment) in a polydisperse holocellulose suspension following mechanical blending.

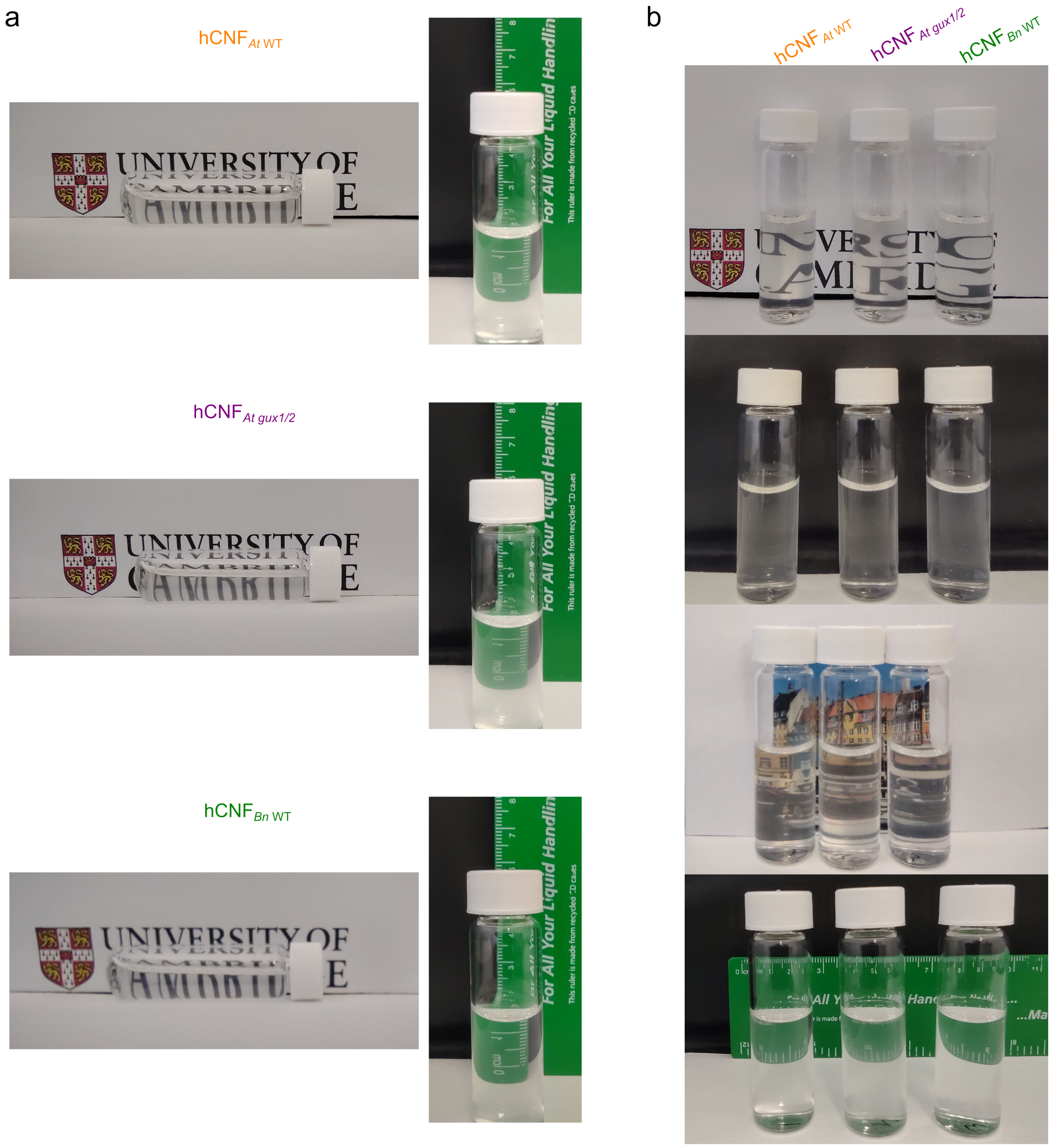

**Figure S6.** Visual comparison of hCNF colloidal suspensions. a) Photographs of individual hCNF suspension types. b) hCNF suspensions displayed side by side against different backgrounds to facilitate qualitative comparison of transparency.

2.2.3 Microscopy

Transmission electron microscopy (TEM) imaging of hCNFs and xyl-hCNFs: An aliquot of sample (2.5 µL) was deposited onto a freshly glow-discharged carbon-coated grid and allowed to adsorb for 40 s. Excess liquid was removed by manual blotting with filter paper, after which the grid was negatively stained with 2% (w/v) uranyl acetate (2.5 µL) for 40 s. The grids were air-dried at ambient temperature for 3 min prior to imaging. TEM images were acquired using a Talos F200X G2 microscope (Thermo Fisher Scientific) operated in scanning transmission electron microscopy mode at an accelerating voltage of 200 kV.

Cryogenic transmission electron microscopy (cryo-TEM) imaging of hCNFs and xyl-hCNFs: All samples were diluted to 0.01 wt% with distilled water prior to grid preparation. Aliquots (3.2 µL) were applied to glow-discharged holey carbon grids (Quantifoil Cu R1.2/1.3, 300 mesh), blotted for 3 s with a blot force of 0, and vitrified by plunge-freezing into liquid ethane using a Vitrobot Mark IV (Thermo Fisher Scientific) operated at 4 °C and 95% relative humidity. Cryo-TEM imaging was performed using a Titan Krios microscope (Thermo Fisher Scientific) equipped with a Falcon 4i direct electron detector at the Department of Biochemistry, University of Cambridge.

TEM and cryo-TEM image data analysis: TEM and cryo-TEM micrographs were analyzed using Fiji software.[8] Fibril widths were measured manually from a total of 700–800 measurements obtained from at least two to three TEM micrographs and one cryo-TEM micrograph per sample. The resulting data were imported into QtiPlot for visualization and statistical analysis. One-way analysis of variance (ANOVA) was performed in QtiPlot to assess statistically significant differences between mean hCNF widths.

Atomic force microscopy (AFM) imaging of hCNFs: hCNF suspensions (10 µL) were deposited onto freshly cleaved mica substrates functionalized with 0.5% (w/v) (3-aminopropyl)triethoxysilane (APTES) and incubated for 5 min. Excess sample was removed by rinsing with 1 mL of Milli-Q water, followed by drying under a gentle nitrogen flow. AFM imaging was conducted using an NX10 AFM system (Park Systems) operated in non-contact mode under ambient conditions. Imaging was performed in a constant phase-shift regime to minimize sample deformation. A silicon nitride cantilever (PPP-NCHR) with a nominal tip radius <10 nm and a spring constant of 5 N $m^{-1}$ was used. Image processing was carried out using SPIP software (Image Metrology), and images were flattened using first-order correction.

2.2.4 Supporting microscopy images

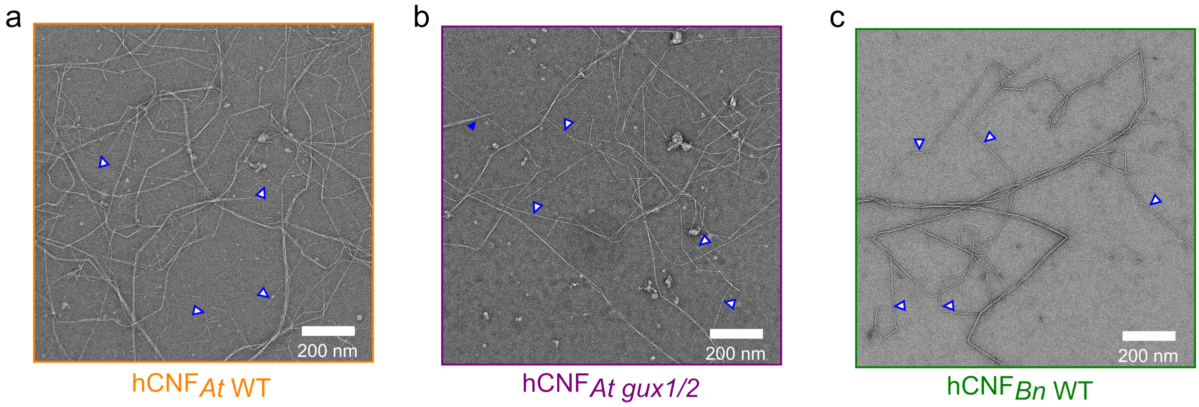

**Figure S7.** Transmission electron microscopy (TEM) images of hCNFs highlighting fibril morphology. TEM images of hCNFs of a) *At* WT, b) *At gux1/2*, and c) *Bn* WT highlighting the presence of individual fibrils (indicated by blue markers) and loosely bundled fibrils.

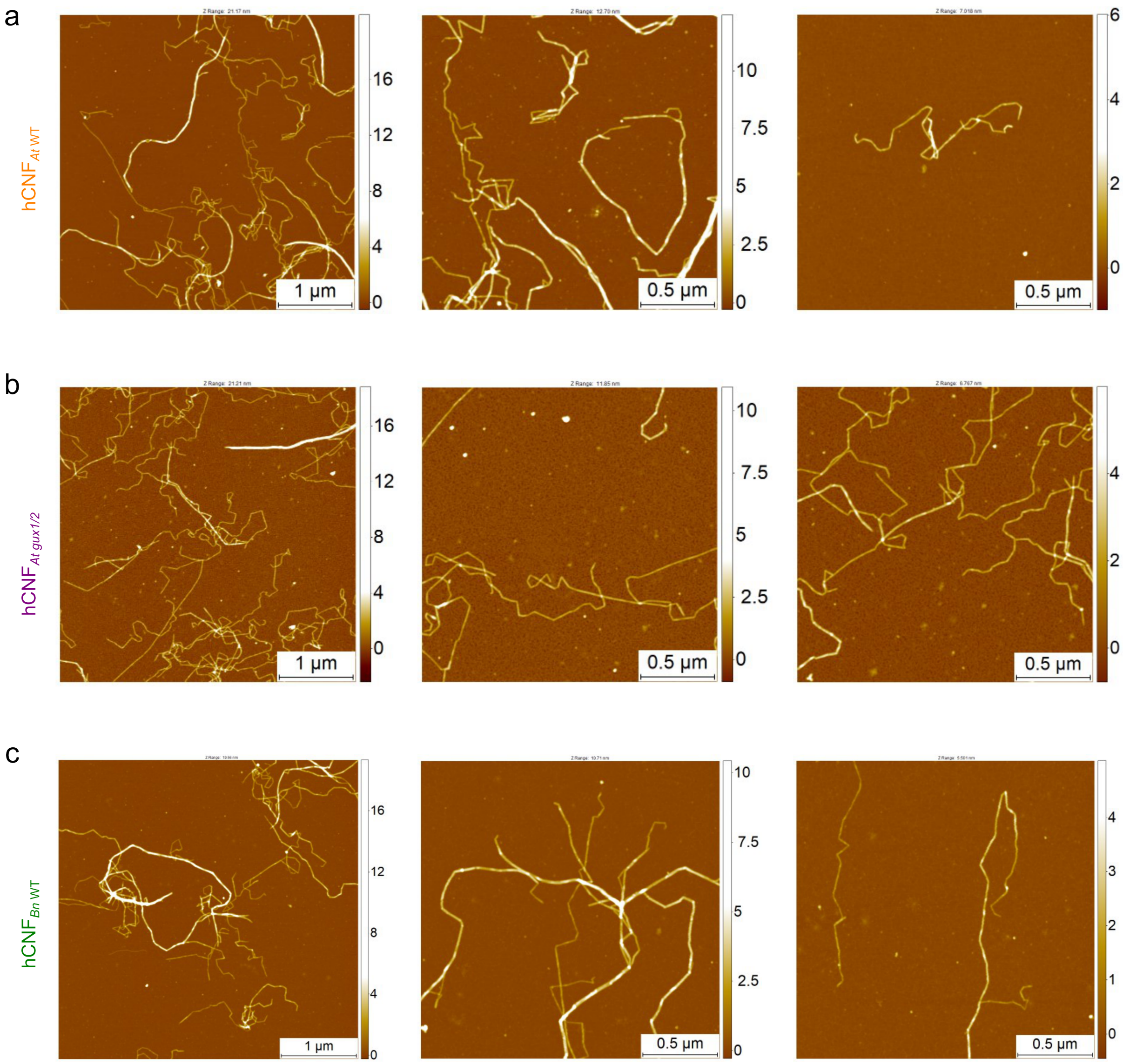

**Figure S8.** Atomic force microscopy (AFM) images of hCNFs. AFM images of hCNFs of a) *At* WT, b) *At gux1/2*, and c) *Bn* WT.

2.2.5 Supporting lignin and neutral monosaccharide data for holocellulose and holocellulose nanofibrils

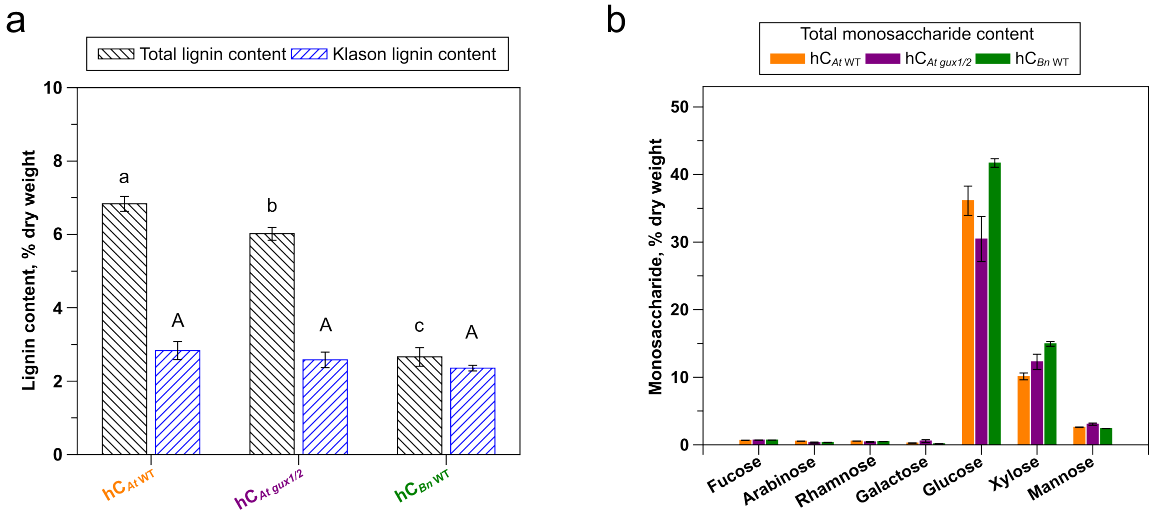

**Figure S9.** Compositional analysis of holocellulose (hC) substrates. a) Total lignin and Klason lignin contents of different holocelluloses, determined by UV–vis spectroscopy after cysteine–sulfuric acid hydrolysis[4] and by gravimetric analysis following sulfuric acid hydrolysis,[5] respectively. For comparison, the total lignin contents of the corresponding lignocelluloses (LCs) before delignification were 16.9, 15.9, and 18.8% dry weight for *At* WT, *At gux1/2*, and *Bn* WT, respectively (Figure S1). Mean values not sharing the same letter are significantly different according to Tukey's test at the 5% significance level. b) Total neutral monosaccharide content of the holocellulose substrates. Data are presented as mean ± SD (n = 3).

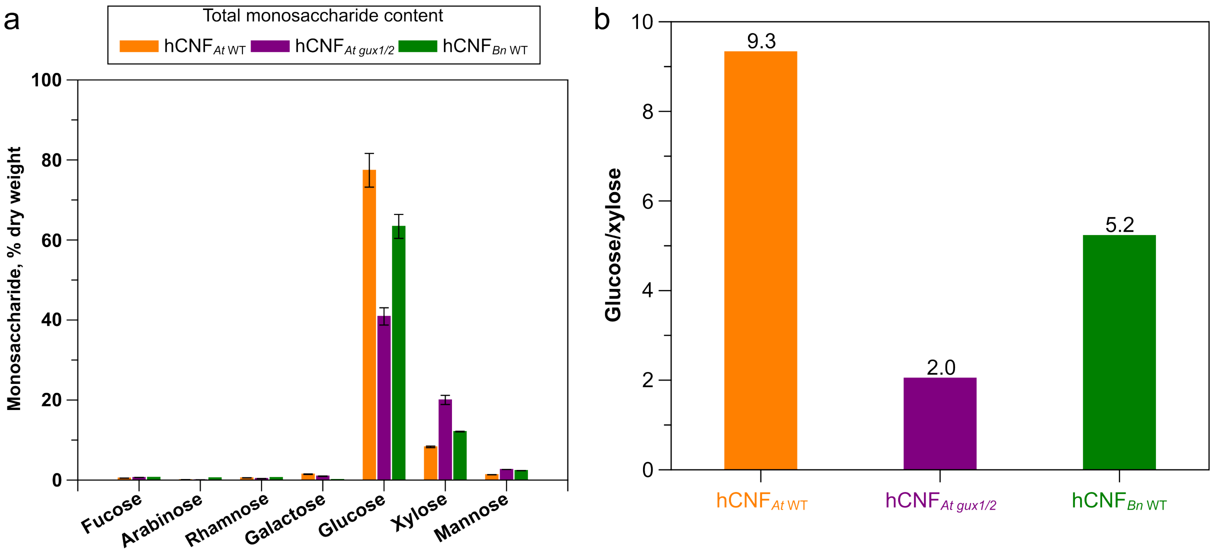

**Figure S10.** Neutral monosaccharide composition of hCNFs. a) Total neutral monosaccharide content in holocellulose nanofibrils (hCNFs). Data are presented as mean ± SD (n = 3). b) Glucose-to-xylose ratios of the different hCNF samples.

#### 3. Magic-angle spinning NMR (MAS NMR) and wide-angle X-ray scattering (WAXS)

##### *3.1 $^{13}$C-enriched Arabidopsis thaliana* wild type *(At* WT*) plant used for MAS NMR*

$^{13}$C-labeled *Arabidopsis thaliana* wild-type plants ($^{13}$C-labeled $LC_{At\ WT}$) were grown and isotopically enriched using $^{13}CO_2$ in a controlled growth chamber, as described previously.[9] Harvested plants were stored at −80 °C prior to use. For all experiments, only the bottom stem segments were selected, as they are enriched in secondary cell walls. The never-dried bottom stems were processed following chlorophyll removal by incubation in 96% ethanol at 70 °C for 30 min, followed by extensive washing with deionized water.

##### *3.2 Solid-state magic-angle spinning nuclear magnetic resonance (MAS NMR) spectroscopy*

All solid-state NMR experiments were performed on a Bruker 1 GHz AVANCE NEO solid-state NMR spectrometer (Bruker), operating at $^{1}$H and $^{13}$C Larmor frequencies of 1000.4 and 251.6 MHz, respectively. Samples were packed into 3.2 mm rotors, and spectra were acquired using a 3.2 mm $E^{Free}$ triple-resonance magic-angle spinning (MAS) probe. All measurements were conducted at 10 °C with a MAS frequency of 12.5 kHz. $^{13}$C chemical shifts were externally referenced to the carbonyl resonance of L-alanine at 177.8 ppm relative to tetramethylsilane. Typical 90° pulse lengths were 3.1 µs for $^{1}$H and 4.4 µs for $^{13}$C. All $^{1}$H–$^{13}$C cross-polarization (CP)-based experiments employed a ramped[10] $^{1}$H radiofrequency field (70–100%), a contact time of 1 ms, and a recycle delay of 2 s. During signal acquisition, SPINAL-64 heteronuclear decoupling[11] was applied with a $^{1}$H nutation frequency of 70–80 kHz.

To resolve distinct cellulose glucosyl environments within the two cellulose domains, CP-refocused INADEQUATE experiments were performed on $^{13}$C-labeled *Arabidopsis thaliana* wild-type stems ($^{13}$C labeled $LC_{At\ WT}$), $^{13}$C-labeled *Arabidopsis thaliana* wild-type holocellulose ($^{13}$C labeled $hC_{At\ WT}$), and $^{13}$C-labeled *Arabidopsis thaliana* wild-type holocellulose nanofibrils ($^{13}$C labeled $hCNF_{At\ WT}$). The CP-refocused INADEQUATE experiment correlates directly bonded carbon nuclei within the same glucan ring through a double-quantum dimension.[12,13] The indirect-dimension acquisition time was 5.1–6.0 ms, with a spectral width of 50 kHz and 128 acquisitions per $t_1$ FID. The spin-echo delay (τ) was set to 2.24 ms, corresponding to a total spin-echo duration of 8.96 ms.

Intramolecular $^{13}$C–$^{13}$C correlations were probed for all samples using two-dimensional $^{13}$C–$^{13}$C proton driven spin diffusion (PDSD) experiments with a mixing time of 30 ms.[14] For CP-PDSD experiments, the indirect-dimension acquisition time ($t_1$) was 5.1–6.1 ms, with spectral widths of 37.5–50 kHz and 48 acquisitions per $t_1$ FID.

All two-dimensional spectra were processed using Bruker TopSpin software (v3.6) with Fourier transformation into 8k ($F_2$) × 2k ($F_1$) data points. Exponential line broadening of 20 Hz was applied in the direct dimension ($F_2$), and a shifted sine-bell (QSine = 3) window function was used for processing in the indirect dimension ($F_1$).

*3.3 Supporting data for MAS NMR*

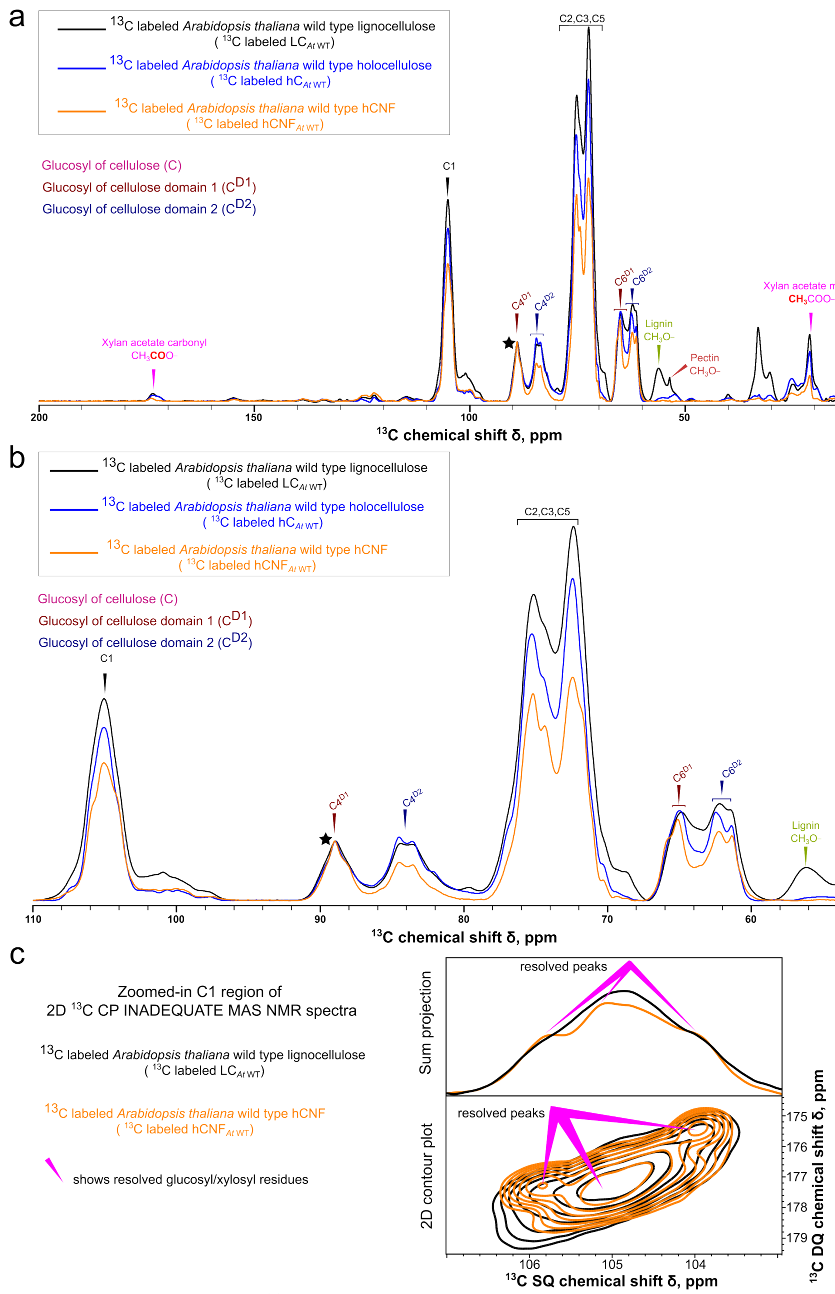

**Figure S11.** Comparison of 1D $^{13}$C CP MAS NMR spectra of stem, holocellulose, and hCNFs of $^{13}$C labeled *Arabidopsis thaliana* wild type ($^{13}$C labeled LC$_{At\ \mathrm{WT}}$) plants, and evidence of better resolution in hCNFs. a) Overlay of 1D $^{13}$C CP MAS NMR spectra over the 0–200 ppm range for never-dried stem (black), holocellulose (blue), and hCNFs (orange). b) Overlay of the expanded 50–100 ppm carbohydrate region of 1D $^{13}$C CP MAS NMR spectra for the same samples. Spectra are normalized to the C4$^{D1}$ signal (asterisked) at ~89ppm. c) Comparison of the cellulose C1 region of 2D $^{13}$C CP refocused INADEQUATE MAS NMR spectra of $^{13}$C-labelled wild-type *Arabidopsis thaliana* lignocellulose (LC$_{At\ \mathrm{WT}}$, black) and the corresponding hCNFs (hCNF$_{At\ \mathrm{WT}}$, orange). The 2D contour plots (bottom) show a small increase in the apparent resolution of glucosyl and xylosyl C1 environments for the hCNFs (magenta markers); the sum projections onto the SQ ($^{13}$C) axis are shown above (top). The hCNF spectrum shows better-resolved C1 environments than the parent lignocellulose, evident as a more pronounced shoulder/split in the sum projection and clearer separation of glucosyl and xylosyl correlations in the 2D contour plot.

**Table S4.** Comparison of $^{13}$C chemical shifts for glucosyl residues in cellulose $^{13}$C labeled LC$_{At\ \mathrm{WT}}$ and hCNF$_{At\ \mathrm{WT}}$, as determined by 2D $^{13}$C CP refocused INADEQUATE and CP 30 ms PDSD MAS NMR.

| Carbon/Domain | Lignocellulose ($^{13}$C labeled LC$_{At\ \mathrm{WT}}$) chemical shift, ppm | | | Isolated hCNF ($^{13}$C labeled hCNF$_{At\ \mathrm{WT}}$) chemical shift, ppm | | |
|---|---|---|---|---|---|---|
| C1 | | | | | | |
| Domain1/2 | - | 105.2 | - | 105.8* | 105.2 | 104.1* |
| C2 | | | | | | |
| Domain1/2 | n.a. | 72.5 | n.a. | n.a. | 72.4 | n.a. |
| C3 | | | | | | |
| Domain1/2 | 75.2 | 74.3 | n.a. | 75.4 | 74.3 | n.a. |
| C4 | | | | | | |
| Domain1 | 88.9 | 89.1 | 88.1 | 89.0 | 89.1 | 88.1 |
| Domain2 | 84.5 | 83.5 | 83.3 | 84.5 | 83.5 | 83.3 |
| C5 | | | | | | |
| Domain1 | 72.5 | 71.2 | n.a. | 72.5 | 71.2 | n.a. |
| Domain2 | 75.3 | 75.2 | 73.8 | 75.3 | 75.2 | 73.8 |
| C6 | | | | | | |
| Domain1 | 65.8 | 65.1 | n.a. | 65.8 | 65.1 | n.a. |
| Domain2 | 62.4 | 61.4 | 61.3 | 62.4 | 61.4 | 61.3 |

Domain 1/2 indicates unresolved spectral domains 1 and 2 for C1, C2, and C3; "–" denotes an unresolved peak in LC$_{At\ \mathrm{WT}}$; * indicates peaks resolved in the hCNFs; n.a., not applicable (unresolved peak in both the LC and hCNF samples).

*3.4 Wide angle X-ray scattering (WAXS)*

Thick films (70–115 µm) were prepared for wide-angle X-ray scattering (WAXS) measurements. A degassed 0.1 wt% dispersion was vacuum-filtered through a 0.45 µm PVDF membrane to form a wet cake. The cake was covered with a second PVDF membrane and sandwiched between filter papers (two on each side). Films were dried under vacuum at 80 °C for 10 min using a gel dryer. For comparison, additional samples were dried by air-drying, freeze-drying, or oven-drying at 45 °C.

WAXS measurements were conducted using an X-ray beam energy of 15 keV (λ = 0.8266 Å) and a camera length of 693.9 mm. Diffraction patterns were collected with a Pilatus3S 2M detector (1475 × 1679 pixels; pixel size 172 µm × 172 µm; Dectris, Switzerland). One-second exposures were recorded, with four diffractograms collected per sample in a 2 × 2 matrix and averaged prior to peak deconvolution.

Diffraction profiles were fitted assuming a cellulose Iβ crystal structure. After deconvolution, reflections corresponding to cellulose Iβ were assigned to the d-spacings (*d*) of $d_{1\text{-}10}$, $d_{110}$, $d_{102}$, $d_{200}$, and $d_{004}$. The peak maxima position, expressed as the scattering vector (*q*), is directly related to the lattice spacing (*d*), which was calculated according to:

$$d = \frac{2\pi}{q} \quad (2)$$

Changes in crystal packing were evaluated from shifts in peak maxima. Crystallite size (L) was estimated from the full width at half maximum (FWHM, $\Delta q$) using:

$$L = \frac{2\pi}{\Delta q} \quad (3)$$

Raw X-ray scattergrams (32-bit, 1475 × 1679 pixels) were processed using ImageJ (v1.54f).[15] Images were centered using silver behenate as a calibration standard. Calibrated intensity profiles (in q space) were converted to polar coordinates (radius = q) and further processed in R (v4.2.1). Data were summarized to three decimal places in q and smoothed using a rolling mean.

Baseline correction was performed by fitting a convex hull to plots of intensity versus q. The q range and baseline were interactively adjusted to prevent peak truncation. Inflection points in the intensity–q profiles were identified from peaks in the second derivative and used to determine the number and initial positions of peaks for deconvolution. Peak fitting was carried out using a pseudo-Voigt mixture model via the spect_em_pvmm function from the EMpeaksR package (v0.3.1). The q range, number of peaks, and initial peak positions were standardized across all samples. An example of the peak deconvolution procedure is shown in Figure S12.

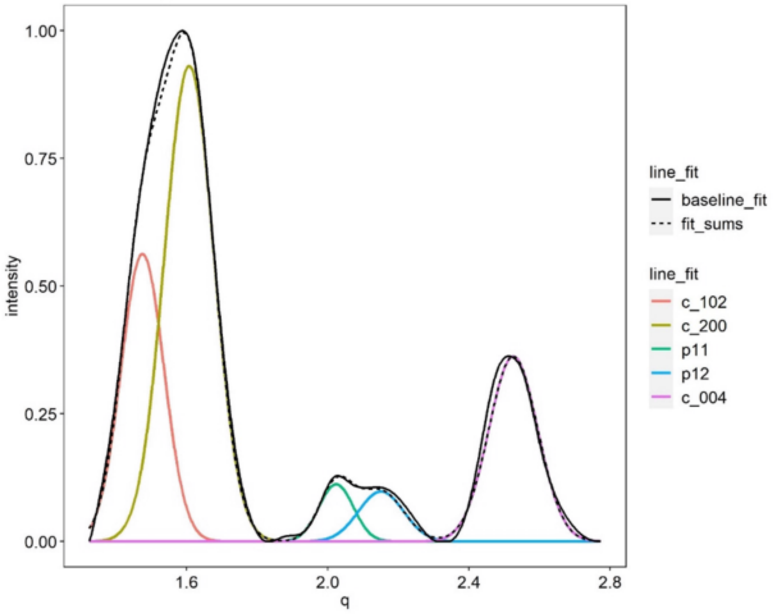

**Figure S12.** Representative example of peak deconvolution applied to WAXS data.

#### 4. Xylanase treatment and modeling of cellulose Iβ fibrils

##### *4.1 Xylanase cocktail digestion of hCNFs*

hCNF suspensions were enzymatically treated with GH10 xylanase, CE4 acetyl xylan esterase, and GH115 α-glucuronidase on an orbital shaker (120 rpm) at 30 °C in 0.1 M ammonium acetate buffer (pH 6.0) at a solids content of 0.075 wt%. After 20 h, enzymatic activity was quenched by heating the reaction mixture to 100 °C for 10 min, followed by centrifugation at 10000 rpm for 20 min at 4 °C. The resulting supernatant was filtered through a 0.1 μm PVDF membrane.

The centrifugation pellet and, where present, the membrane retentate were combined and redispersed in 65% (v/v) ethanol–water. This centrifugation–filtration procedure was repeated once for the ethanol–water mixture and then three times with deionized water. The final undigested fraction was resuspended in water at 0.10–0.12 wt% and mechanically dispersed by blending for 2 min using a Vitamix A3500i blender. The resulting material is hereafter referred to as xylanase-treated holocellulose nanofibrils (xyl-hCNFs). The xyl-hCNF suspensions were stored at 4 °C prior to characterization.

Control samples were prepared in parallel in the absence of enzymes and processed under identical conditions. Unlike for hCNFs, no attempt was made to separate unfibrillated material by centrifugation. Representative photographs of xyl-hCNF suspensions are shown in Figure S13.

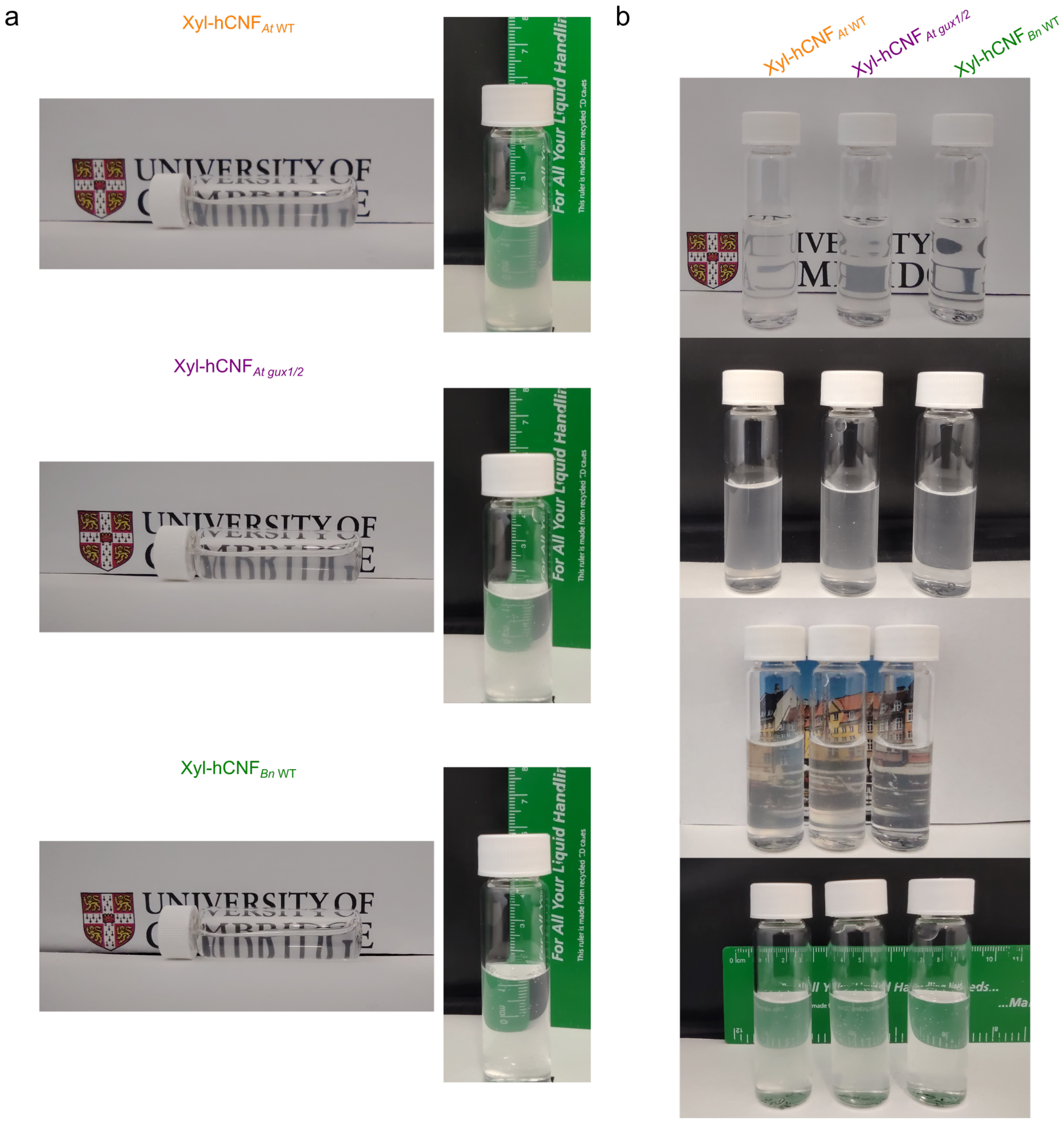

**Figure S13.** Visual comparison of xylanase-treated hCNF (xyl-hCNF) suspensions. a) Photographs of the xyl-hCNF suspension types. b) Comparison of the transparency of three xyl-hCNF suspensions placed side by side against different backgrounds.

*4.2 Supporting data for fibril width measurement*

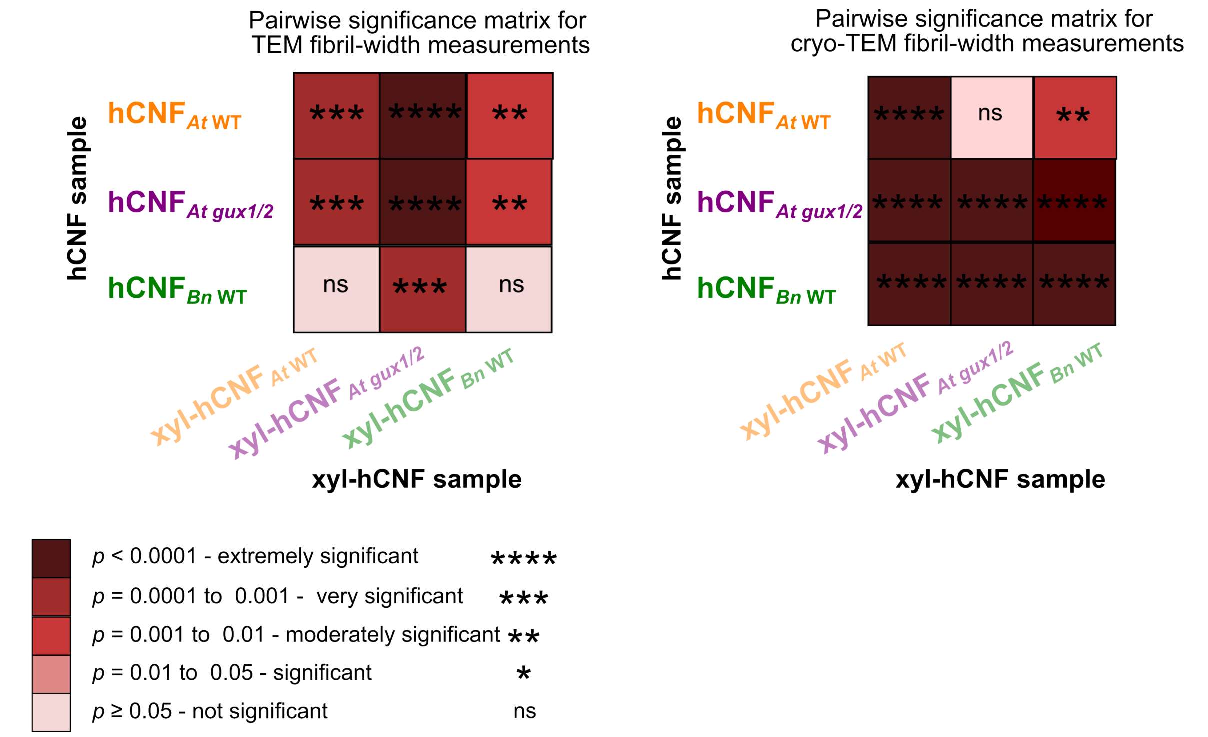

**Figure S14**. Pairwise statistical comparison of fibril widths measured by TEM and cryo-TEM for hCNFs and xyl-hCNFs. Statistical significance was assessed by one-way analysis of variance (ANOVA) followed by Tukey's post hoc test for multiple comparisons.

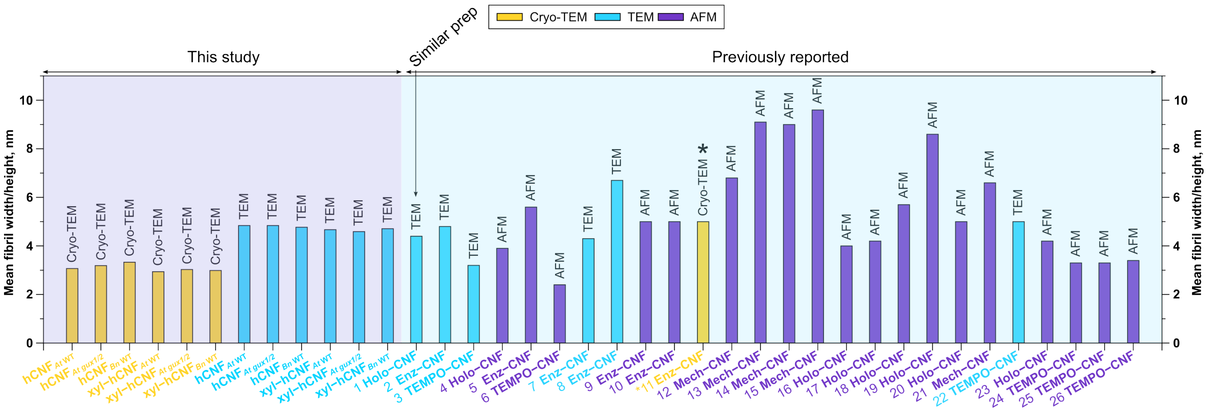

**Figure S15.** Comparison of fibril width measurements by TEM, cryo-TEM, and AFM across studies. The figure compares fibril widths obtained in this work with values reported in the literature using TEM, cryo-TEM, and AFM. Cryo-TEM consistently yields smaller fibril widths than TEM and AFM measurements. Although TEM and AFM are the most widely used techniques, cryo-TEM avoids drying and negative staining and may therefore more accurately reflect native fibril diameters. Literature data points are indexed as follows: 1-6: [7]; 7-8: [15]; 9: [16]; 10-11: [17]; 12-15: [18]; 16-19: [19]; 20: [20]; 21: [21]; 22: [22]; 23-24: [23]; 25: [24]; 26: [25]

*4.3 Computation of theoretical cellulose microfibril dimensions*

Cellulose Iβ microfibril models were constructed using the Cellulose-Builder toolkit,[26] generating structures composed of eight anhydroglucose units per chain. These chains were assembled into microfibrils containing either 18 or 24 chains to represent the habits 234432, 333333, 12333321, 33333333, and 12345432 (Figure S16).[27–29] The resulting models provided full atomic coordinates and atom types for each configuration.

Van der Waals (vdW) surfaces were calculated for each model using the volmap function implemented in VMD (Visual Molecular Dynamics).[30] Volumetric maps were generated with a grid resolution of 0.6 Å and a cutoff distance of 4.0 Å, and the vdW surfaces were exported as coordinate point clouds. These surfaces define the effective microfibril boundaries in a manner analogous to the surrounding electron density and represent the union of atomic spheres defined by vdW radii from the carbohydrate CHARMM36 force field (Figure S16a).[31]
Microfibril diameters were determined by calculating the distance between the centroid (geometric center) of the structure and points on the vdW surface, yielding direction-dependent radii. To focus on cross-sectional dimensions perpendicular to the longitudinal fibrils axis, radii were computed at multiple positions along the z-axis. The mean radius and corresponding standard deviation were calculated for each cross-section. Radial dimensions were visualized by plotting the radii as a function of polar angle in polar coordinates (Figure S17).

We mapped the cross-sectional region as a function of the polar angle (θ), scanning the fibril along the z-axis and identifying the azimuthal angle (φ) out of plane. The resulting spherical-coordinate data were used to estimate the fibril diameter and its distribution for each architecture. The scripts used for these calculations are provided below.

Computing the cellulose fibril's average radius

Input (for each fibril architecture)
- atom positions and types
- van der Waals radii for each atom type

Compute
- centroid, geometric center
- boundary (matlab function) - returns a vector of point indices representing a single conforming 2-D boundary around the points (x,y). The points (x(k),y(k)) form the boundary. Unlike the convex hull, this boundary can shrink toward the interior of the hull to envelop the points more tightly.
- radii (distances from centroid to boundary points)

The calculation is performed in 2-D, since the modeled cellulose is uniform along its longitudinal axis.

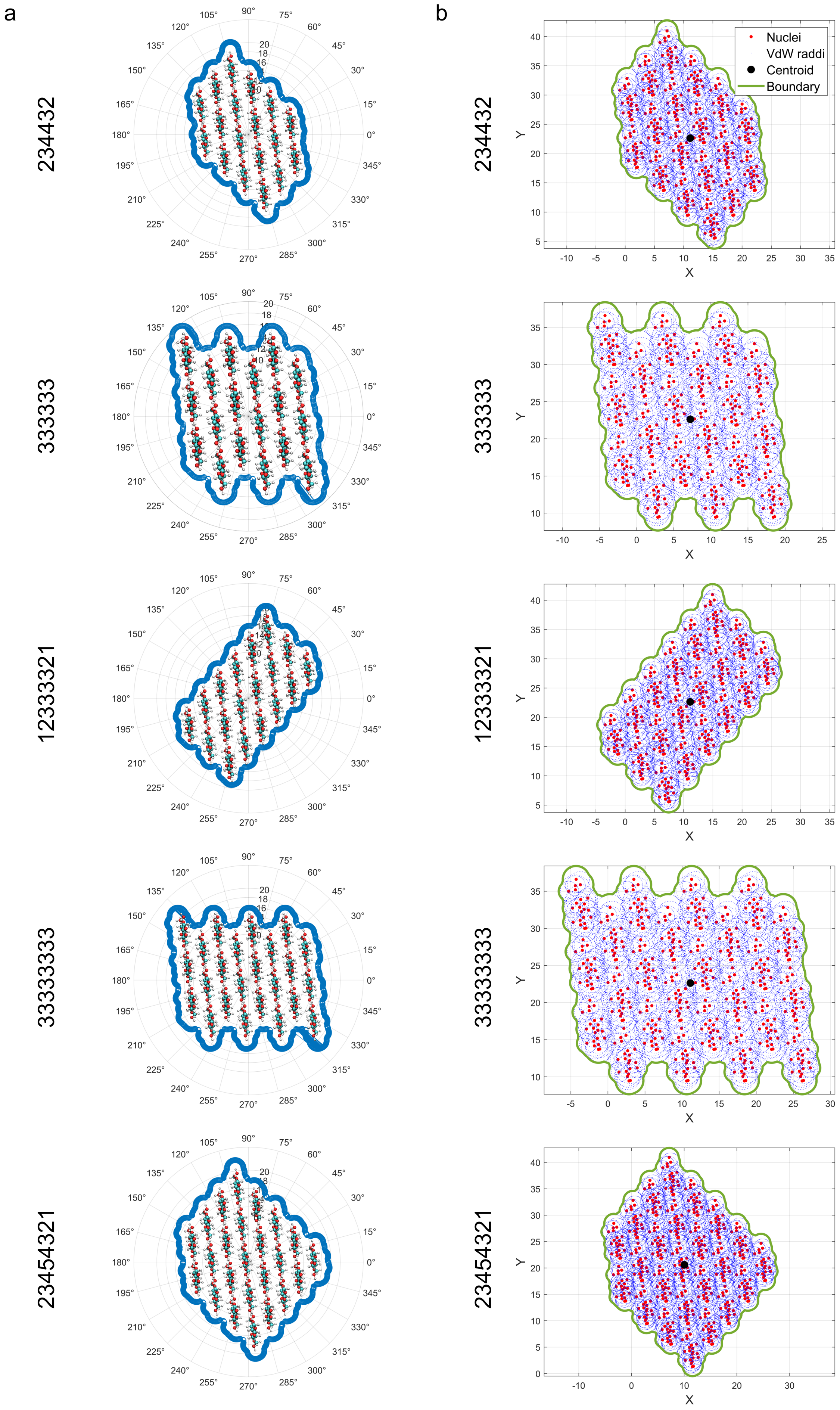
a
b
234432
333333
12333321
33333333
23454321
Nuclei
VdW raddi
Centroid
Boundary
X
Y

**Figure S16.** Cross-sectional models of cellulose Iβ microfibrils. a) Cross-sectional views of 18-chain and 24-chain cellulose Iβ models, illustrating chain arrangement and angular distribution around the fibril center. Van der Waals (vdW) radii were combined to generate vdW surfaces, which were used to calculate the fibril radii. b) Cross-sectional view of a cellulose microfibril showing the combined vdW spheres for the first glucose residue. The mean apparent diameters calculated per residue are 30.0 ± 6.1 Å (12333321), 29.3 ± 5.2 Å (234432), 29.4 ± 3.9 Å (333333), 33.3 ± 4.5 Å (33333333), and 33.4 ± 4.0 Å (23454321).

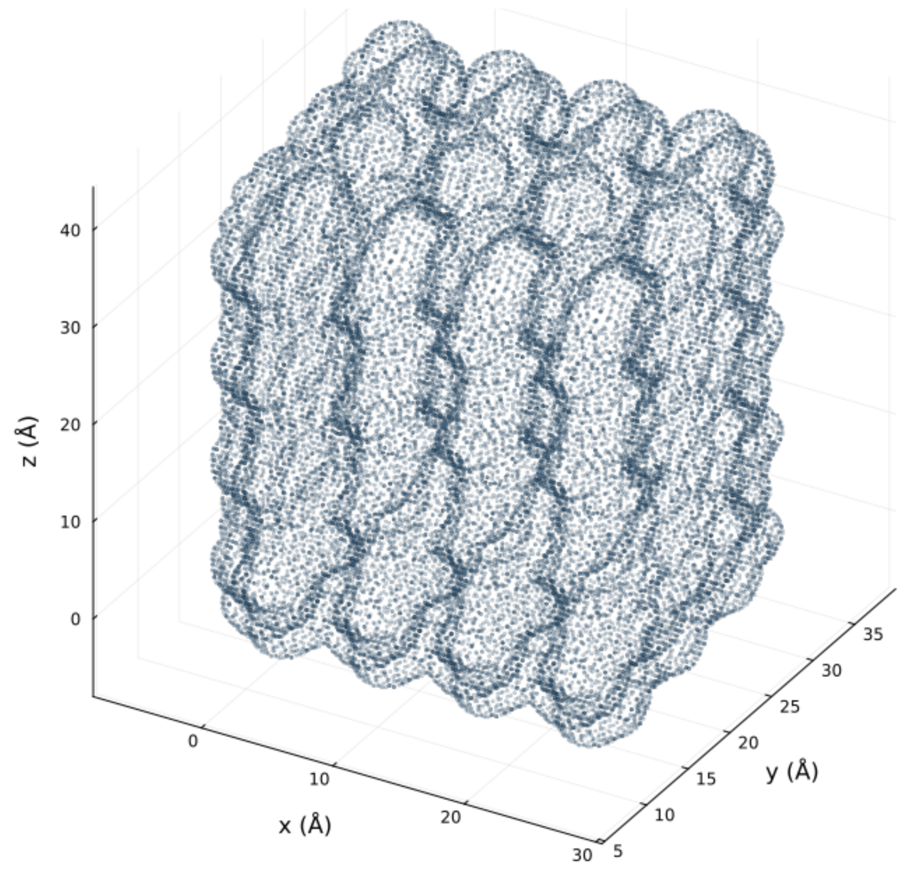

**Figure S17.** Volmap-based analysis of cellulose fibril size distributions. The volmap tool enables higher-resolution determination of cellulose fibril size distributions by exporting isosurfaces with increased surface data density. An example is shown for the 33333333-chain habit, illustrating the enhanced sampling of points around the fibril surface.

#### 5. Nanofibrillation degree, surface charges, and colloidal stability

##### *5.1 Degree of nanofibrillation of hCNFs*

The degree of nanofibrillation of the holocellulose nanofibril (hCNF) suspensions was quantified by comparing the dry mass of hCNFs present in the supernatant after centrifugation with the dry mass of the corresponding polydisperse suspension prior to centrifugation (see Figure S5 for the separation scheme).

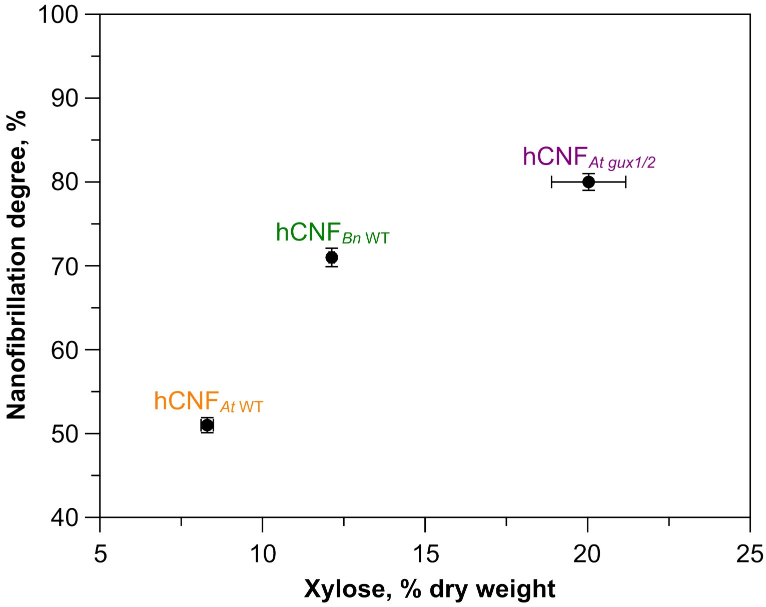

**Figure S18.** Relationship between xylan content and nanofibrillation degree in different hCNFs. Correlation illustrating how changes in xylan content influence fibrillation efficiency. Data are presented as mean ± SD (n = 3–4).

##### *5.2 Carboxylate content*

Carboxylate content was determined by conductometric titration. hCNF dispersions (0.056 wt%) were adjusted to pH 3.5 using 0.1 M HCl and titrated with 0.01 M NaOH. Conductivity and pH were recorded continuously during titration. The carboxylate content was calculated from the plateau corresponding to weak-acid neutralization. Two independent titrations were performed for each sample.

##### *5.3 Zeta potential*

Zeta-potential measurements were carried out using a Zetasizer Nano S (Malvern Panalytical, Malvern, UK) equipped with disposable folded capillary cells (DTS1070). Aqueous hCNF dispersions were analyzed at 25 °C. Zeta potentials were calculated from electrophoretic mobility using the Henry equation with the Smoluchowski approximation. Each reported value represents the average of twenty consecutive runs, with three independent measurements per sample.

*5.4 Light transmittance of hCNF suspensions*

Optical transmittance of hCNF suspensions was measured using a UV–vis spectrophotometer (Shimadzu UV–vis spectrophotometer (UV-1800)) and quartz cuvettes over a wavelength range of 200–800 nm. A water-filled cuvette was used as the blank. For each hCNF system, two to three independently prepared formulations were analyzed, with three technical replicates per formulation. Each replicate consisted of three consecutive measurements.

*5.5 Redispersibility of hCNFs*

Redispersibility of hCNFs was assessed by freeze-drying a 0.1 wt% aqueous suspension, followed by redispersion in water at the original concentration. The redispersed material was mechanically dispersed for 2 min using a Vitamix A3500i blender. Redispersibility was evaluated by comparing the optical transmittance of the suspensions before freeze-drying and after redispersion, as described above.

*5.6 Supporting data for colloidal stability and redispersibility*

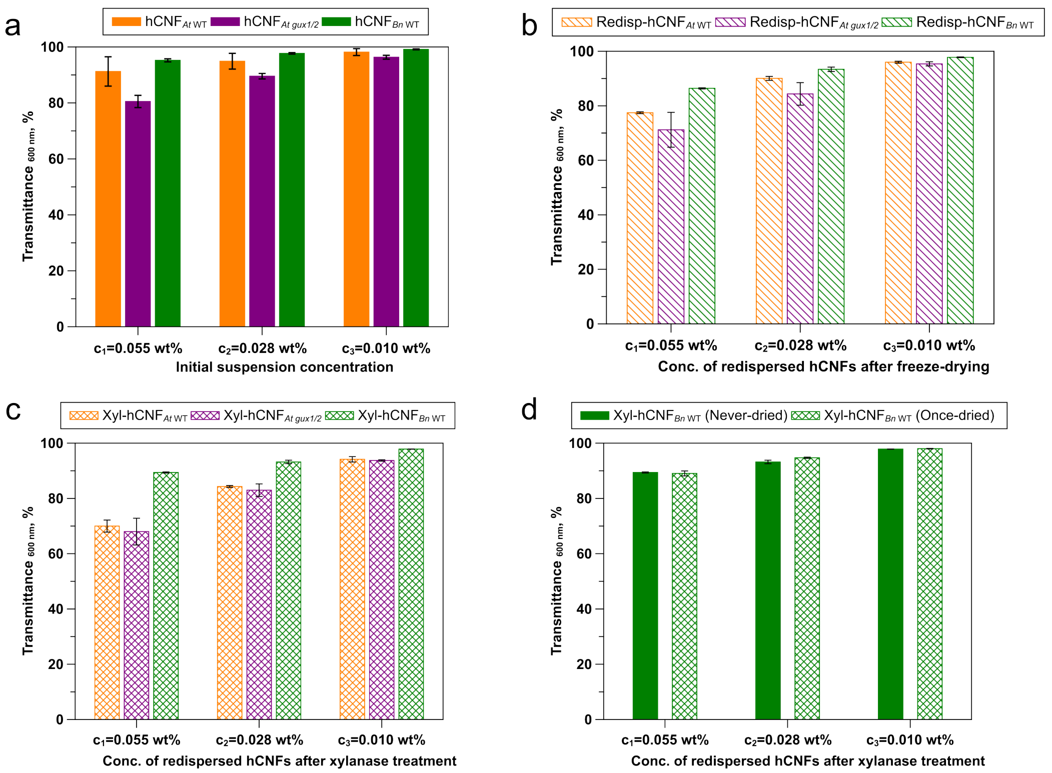

**Figure S19.** Optical transmittance of hCNF and xyl-hCNF suspensions. a-c) Transmittance as a function of suspension concentration for initial hCNFs, redispersed hCNFs, and xylanase-treated hCNFs (xyl-hCNFs), respectively. d) Transmittance of freeze-dried and never dried hCNF*Bn* WT suspensions after xylanase treatment, showing no significant difference and indicating that freeze-drying does not affect the glucuronidated substrate. Data are presented as mean ± SD from 2–3 formulations, each measured with three technical replicates and three measurements per replicate.

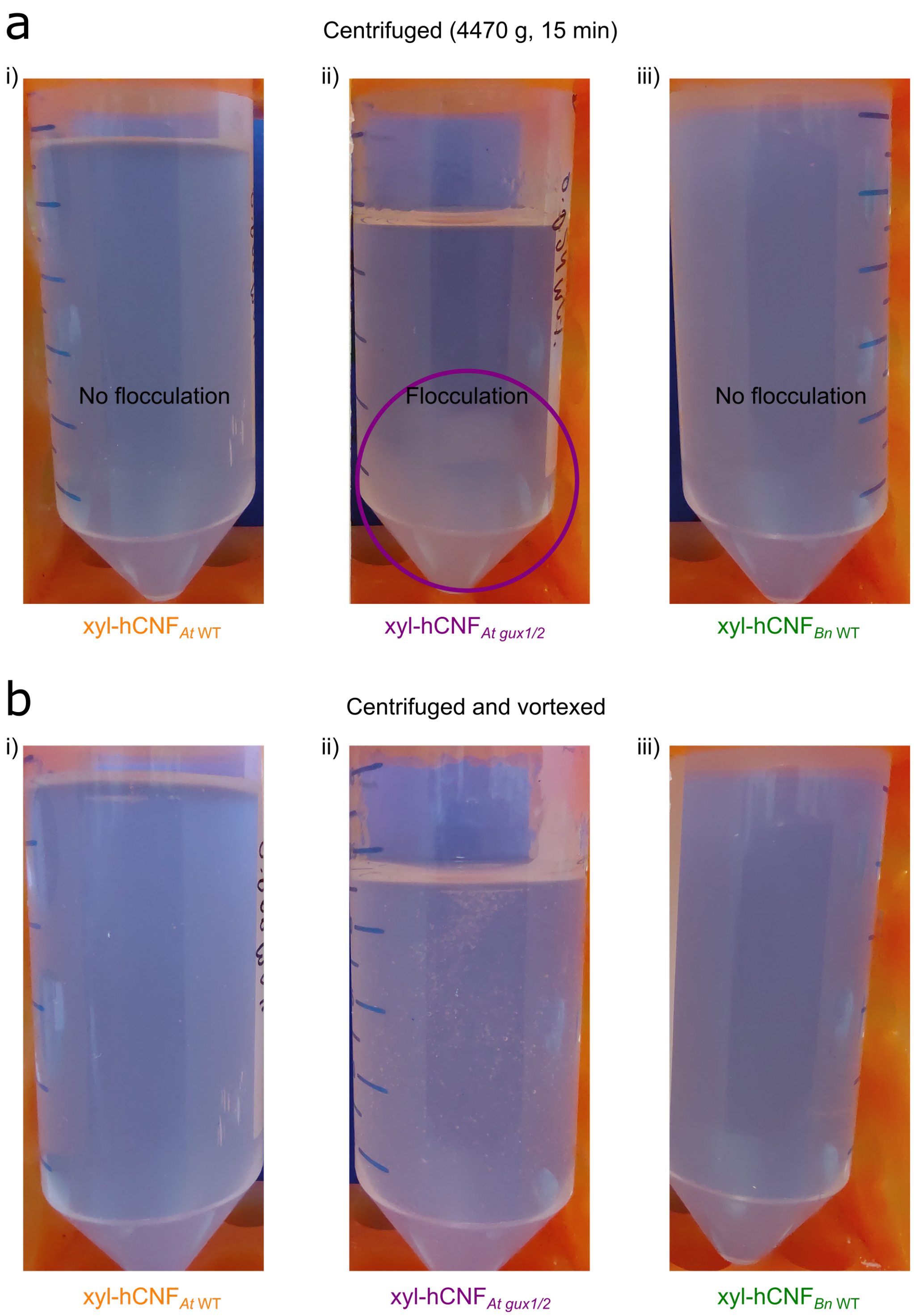

**Figure S20.** Colloidal behavior of xylanase-treated hCNF (xyl-hCNF) suspensions under centrifugation and vortexing. Following centrifugation and subsequent vortexing, the xyl-hCNF$_{At\ gux1/2}$ suspension exhibited pronounced flocculation without sediment formation after centrifugation but cleared upon vortexing. In contrast, xyl-hCNFs containing glucuronidated xylan showed minimal flocculation after centrifugation and similarly return to clear suspensions after vortexing.

#### 6. Moisture sorption and thermal stability of hCNFs

##### *6.1 Dynamic vapor sorption*

Water vapor sorption behavior was investigated using a gravimetric dynamic vapor sorption (DVS) analyzer (Surface Measurement Systems, UK) thermostated at 25 °C. Mass changes were monitored using an electronic Cahn D200 microbalance with a mass resolution of 0.1 µg under a controlled atmosphere. The DVS system records sorption kinetics by continuously measuring he change in mass as a function of time and water activity ($a_w$).

Freeze-dried samples with an initial mass of approximately 10–12 mg were used. Owing to the high reproducibility of the instrument, a single measurement was performed for each sample. Prior to sorption measurements, samples were exposed to a dry nitrogen flow until a constant mass was reached, corresponding to the dry mass ($m_d$). Water vapor was introduced at controlled partial pressures by mixing dry and saturated nitrogen streams using electronic mass flow controllers. Water activity ($a_w$) was increased stepwise from 0 to 0.9 in increments of 0.1, and mass uptake was monitored until equilibrium was reached at each step.

Sorption kinetics and equilibrium sorption isotherms were constructed from the recorded data. The equilibrium water uptake ($\Delta M_{eq}$), expressed as mass percent (grams of water per 100 g of dry sample), was calculated according to:

$$\Delta M_{eq} = \frac{m_{eq} - m_d}{m_d} * 100 \tag{4}$$

where $m_d$ and $m_{eq}$ represent the dry mass and the equilibrium mass, respectively. The uncertainty in equilibrium mass uptake was estimated to be below 2%

The sorption isotherms were modeled using the Park model,[32] which combines dual-mode sorption (Henry-type dissolution and Langmuir-type site-specific sorption) with water-clustering effects, according to:

$$C = \frac{C'_h . b . a}{(1 + b . a)} + k_d . a + n . K_a . k_d^n . a^n \tag{5}$$

Where $C'_h$ and b are the mean concentration and affinity constant of Langmuir sorption sites, $k_d$ is the Henry's law constant describing dissolution of water in the polymer matrix, $K_a$ is the equilibrium constant for water clustering, and $n$ is the mean number of water molecules per cluster.

The quality of the model fit was assessed using the mean deviation modulus (MDM), which provides a measure of agreement between the experimental and predicted values. The Park model was considered to adequately describe the sorption behavior when MDM values were below 10% (Figure S21).[33] The MDM was calculated as:

$$MDM = \frac{100}{N} . \sum_{i=1}^{N} \frac{|m_i - m_{pi}|}{m_i} \tag{6}$$

where $m_i$ is the experimental value, $m_{pi}$ is the corresponding predicted value, and $N$ is the total number of experimental data points.

*6.2 Supporting data for dynamic vapor sorption*

**Table S5.** Park model parameters obtained by fitting the water sorption isotherms of hCNFs.

| hCNF | Langmuir's sorption | | Henry sorption | Water clustering | |
|---|---|---|---|---|---|
| | Mean concentration $C'_h$, g/g | Affinity constant, b | Henry constant $k_d$, g/g | Equilibrium Constant $K_a$ ($x10^5$) | Mean number of $H_2O$, n |
| hCNF$_{At\ WT}$ | 0.026 | 50.00 | 0.15 | 26.7 | 8.3 |
| hCNF$_{At\ gux1/2}$ | 0.027 | 21.83 | 0.12 | 19.9 | 7.9 |
| hCNF$_{Bn\ WT}$ | 0.026 | 34.35 | 0.15 | 12.2 | 6.9 |

Langmuir-type sorption, Henry's-law sorption, and water clustering predominated in the low ($a_W$= 0–0.2), medium ($a_W$=0.2–0.7), and high ($a_W$=0.7–0.9) water-activity regimes, respectively. The data presented are derived from a single measurement, owing to the high reproducibility of the instrument.

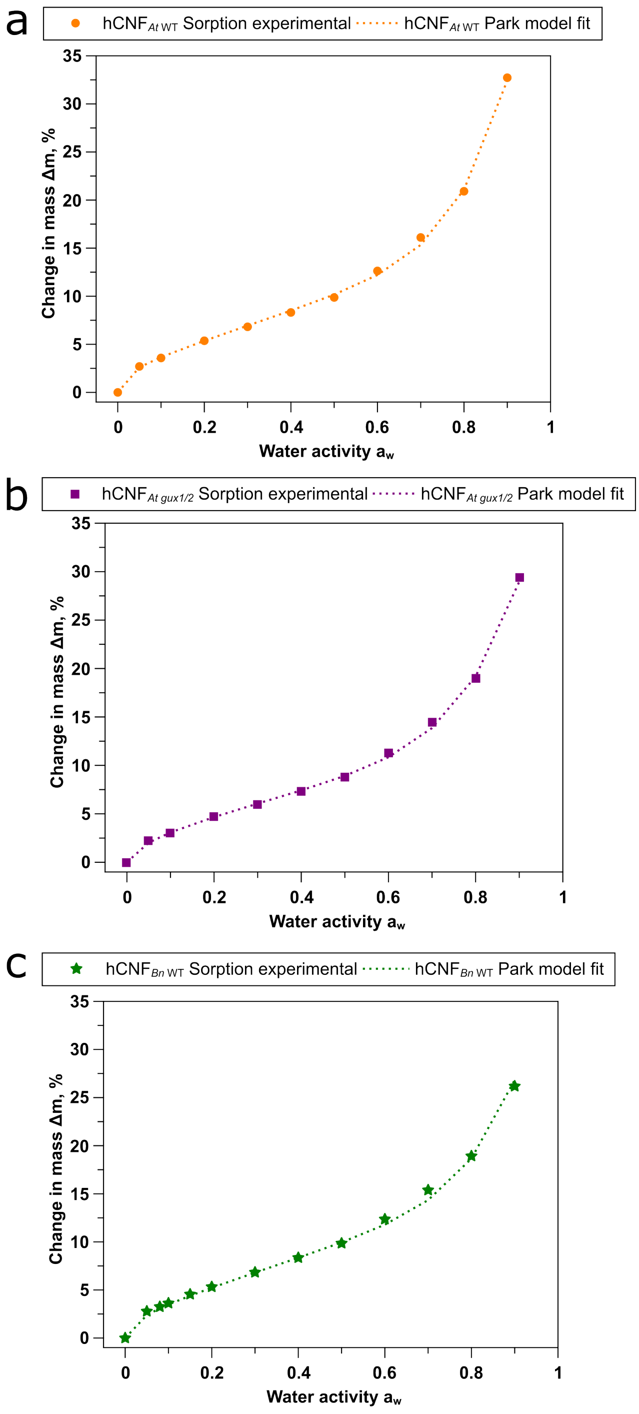

**Figure S21.** Fit of the Park model to water sorption isotherm data for hCNFs. Comparison of the experimental water-sorption isotherms of hCNFs with the fitted Park model, demonstrating the model's ability to describe the observed sorption behavior.

*6.3 Thermogravimetric analysis (TGA)*

Thermogravimetric analysis (TGA) was carried out under an argon atmosphere using a Discovery SDT 650 instrument (TA Instruments) in the Department of Materials Science and Metallurgy, University of Cambridge. Samples were heated from room temperature to 600 °C at a heating rate of 20 °C min$^{-1}$ under a constant argon flow of 50 mL min$^{-1}$. All measurements were performed in triplicate.

*6.4 Supporting data for thermogravimetric analysis*

**Table S6.** Thermogravimetric analysis data for LCs, hCs, hCNFs, and sediments, obtained under an argon atmosphere.

| Sample | Temperature$_{onset}$ ($T_{onset}$), °C | Temperature$_{50\%}$ ($T_{50\%}$), °C | Temperature$_{endset}$ ($T_{endset}$), °C | Final residue, % |
|---|---|---|---|---|
| Lignocellulose LC | | | | |
| *At* WT | 296.9 ± 7.7 | 352.3 ± 11.0 | 387.8 ± 2.0 | 17.9 ± 2.3 |
| *At gux1/2* | 305.3 ± 0.2 | 353.9 ± 1.2 | 385.6 ± 7.1 | 8.4 ± 2.1 |
| *Bn* WT | 290.9 ± 0.1 | 345.8 ± 2.0 | 380.9 ± 0.6 | 1.6 ± 0.9 |
| Holocellulose hC | | | | |
| $hC_{\mathit{At}\ WT}$ | 302.1 ± 3.0 | 344.6 ± 2.1 | 372.3 ± 1.1 | 6.5 ± 3.5 |
| $hC_{\mathit{At\ gux1/2}}$ | 303.9 ± 3.0 | 354.7 ± 0.8 | 378.3 ± 2.0 | 11.5 ± 4.1 |
| $hC_{\mathit{Bn}\ WT}$ | 301.4 ± 4.9 | 347.7 ± 7.3 | 367.2 ± 1.6 | 25.2 ± 3.3 |
| Holocellulose nanofibrils hCNFs | | | | |
| $hCNF_{\mathit{At}\ WT}$ | 273.6 ± 2.5 | 336.7 ± 7.3 | 349.7 ± 2.6 | 27.3 ± 4.0 |
| $hCNF_{\mathit{At\ gux1/2}}$ | 246.7 ± 1.4 | 313.3 ± 0.0 | 345.0 ± 1.6 | 17.8 ± 0.6 |
| $hCNF_{\mathit{Bn}\ WT}$ | 253.6 ± 1.9 | 301.8 ± 11.8 | 328.4 ± 0.7 | 21.4 ± 8.4 |
| Sediments | | | | |
| $Sediment_{\mathit{At}\ WT}$ | 318.4 ± 0.8 | 351.3 ± 12.7 | 375.2 ± 0.9 | 25.8 ± 0.9 |
| $Sediment_{\mathit{At\ gux1/2}}$ | 328.6 ± 0.5 | 366.5 ± 0.7 | 379.0 ± 0.4 | 28.0 ± 2.3 |
| $Sediment_{\mathit{Bn}\ WT}$ | 247.3 ± 0.1 | 360.3 ± 0.2 | 364.3 ± 1.3 | 42.2 ± 2.5 |

Data are reported as mean ± standard deviation (n = 2–3). The onset temperature ($T_{onset}$) denotes the temperature at which mass loss begins; the 50% temperature ($T_{50\%}$) corresponds to the temperature at which 50% of the initial mass is lost; and the endset temperature ($T_{endset}$) indicates the temperature at which mass loss is complete. Sediments are the pellets collected after centrifugation (see scheme in Figure S5).

#### 7. hCNF film preparation and mechanical properties

##### *7.1 Nanopaper film preparation*

Nanopaper films for mechanical testing were prepared by vacuum filtration of degassed 0.1 wt% aqueous dispersions through 0.45 µm PVDF membranes. The resulting wet cake was covered with a second PVDF membrane and sandwiched between filter papers (two sheets on each side). Films were dried under vacuum at 80 °C for 10 min using a gel dryer.

Films with thicknesses of 16–18 µm were prepared from hCNF$_{\textit{At}\ \mathrm{WT}}$, hCNF$_{\textit{At gux1/2}}$, and hCNF$_{\textit{Bn}\ \mathrm{WT}}$ suspensions for both dry and wet tensile testing. In addition, thicker films (27 µm) were prepared from hCNF$_{\textit{Bn}\ \mathrm{WT}}$ to enable direct comparison of tensile properties with values reported in the literature.

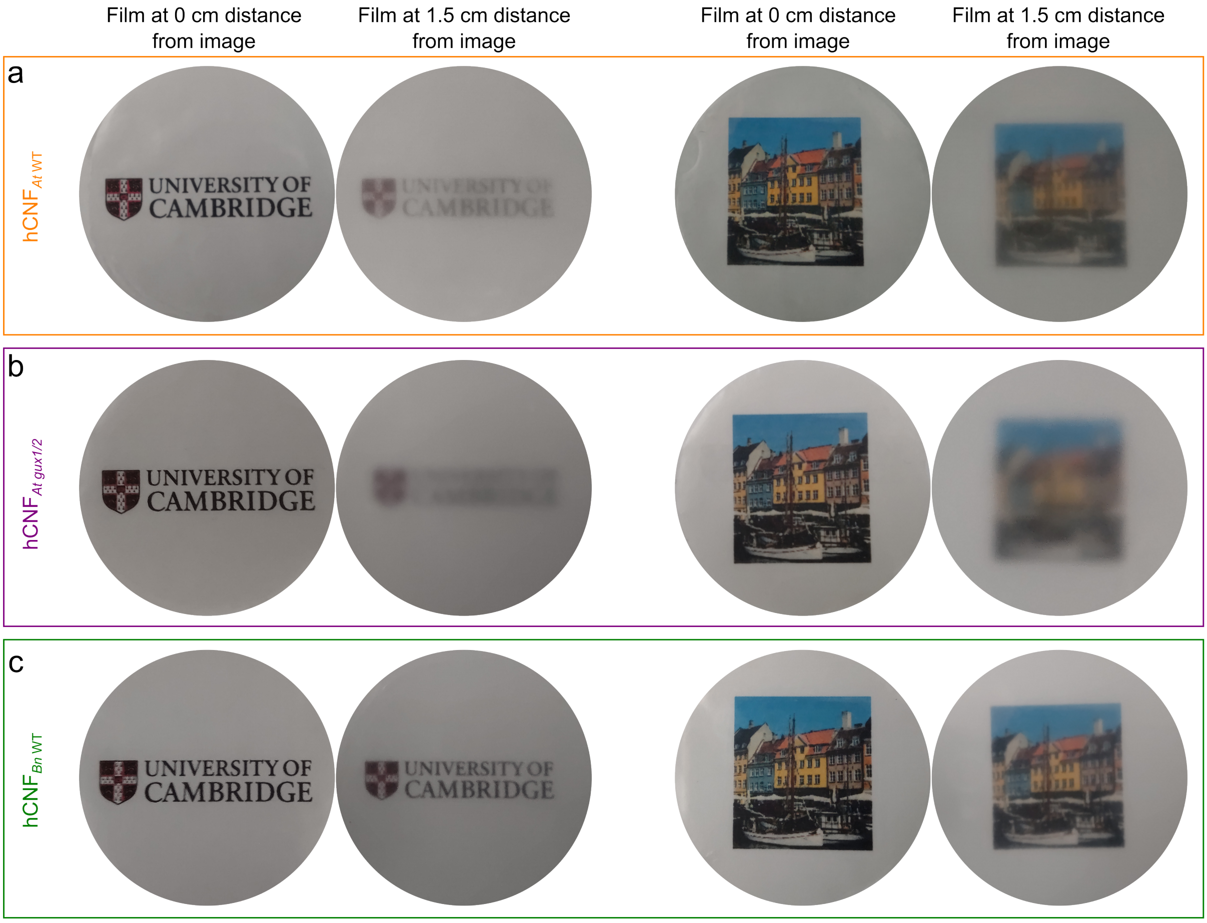

**Figure S22.** Additional images showing the optical transparency of hCNF films. Photographs of hCNF films placed on a printed logo or image at distances of 0 and 1.5 cm.

*7.2 Mechanical properties of films*

Uniaxial tensile tests were performed under both dry and wet conditions. For dry tensile testing, rectangular specimens (30 mm length × 4.3 mm width × 17–18 µm thickness) were cut from nanopaper films prepared from hCNF$_{At\ \mathrm{WT}}$, hCNF$_{At\ gux1/2}$, and hCNF$_{Bn\ \mathrm{WT}}$. For wet tensile testing, rectangular specimens with dimensions of 10 mm length × 2 mm width × 15–16 µm thickness were prepared from the same nanopaper films. In addition, thicker specimens (30 mm length × 4.3 mm width × 27 µm thickness) were prepared from hCNF$_{Bn\ \mathrm{WT}}$ nanopaper films for comparison with literature data.

Dry Tensile Testing: Dry tensile tests were conducted using a universal testing machine (1ST, Tinius Olsen, USA) equipped with a 1 kN load cell. Tests were performed at a crosshead speed of 1 mm min$^{-1}$ with a gauge length of 10 mm at ambient temperature. Prior to testing, specimens were conditioned at 23 °C and 50% relative humidity for at least 24 h. For 18-µm thick films, a minimum of 15 specimens per sample were tested; after exclusion of outliers, at least 9 measurements were retained for data analysis. For the 27-µm thick hCNF$_{Bn\ \mathrm{WT}}$ films, 7 specimens were tested, and 4 measurements were retained after outlier removal.

Wet Tensile Testing: Wet tensile tests were carried out using a custom-built extensometer[34] operated at a crosshead speed of 1.5 mm min$^{-1}$ and a gauge length of 3 mm at ambient temperature. Prior to testing, specimens were immersed in water at 25 °C for 5 min. Between 9 and 12 specimens were tested for each sample; following outlier removal, at least 8 measurements per sample were used for reporting.

*7.3 Supporting data for mechanical properties of hCNF films*

**Table S7.** Mechanical properties of hCNF films measured under dry and wet conditions.

| Film | Max. tensile strength $\sigma_t$, MPa | Tensile strain at break $\varepsilon_b$, % | Young's modulus E, GPa |
|---|---|---|---|
| Dry state films | | | |
| hCNF$_{At\ \mathrm{WT}}$ | 173.2 ± 15.0 | 6.6 ± 1.0 | 4.21 ± 0.74 |
| hCNF$_{At\ gux1/2}$ | 180.6 ± 5.6 | 5.3 ± 0.6 | 4.49 ± 0.85 |
| *hCNF*$_{Bn\ \mathrm{WT}}$ | 190.6 ± 10.4 | 6.9 ± 1.1 | 4.29 ± 0.57 |
| Wet state films | | | |
| hCNF$_{At\ \mathrm{WT}}$ | 15.0 ± 1.6 | 66.4 ± 6.6 | 0.0540 ± 0.0100 |
| hCNF$_{At\ gux1/2}$ | 2.4 ± 0.4 | 16.4 ± 2.4 | 0.0278 ± 0.0070 |
| *hCNF*$_{Bn\ \mathrm{WT}}$ | 15.3 ± 0.6 | 69.4 ± 4.8 | 0.0414 ± 0.0036 |

Data are presented as mean ± SD (n = 8–9).

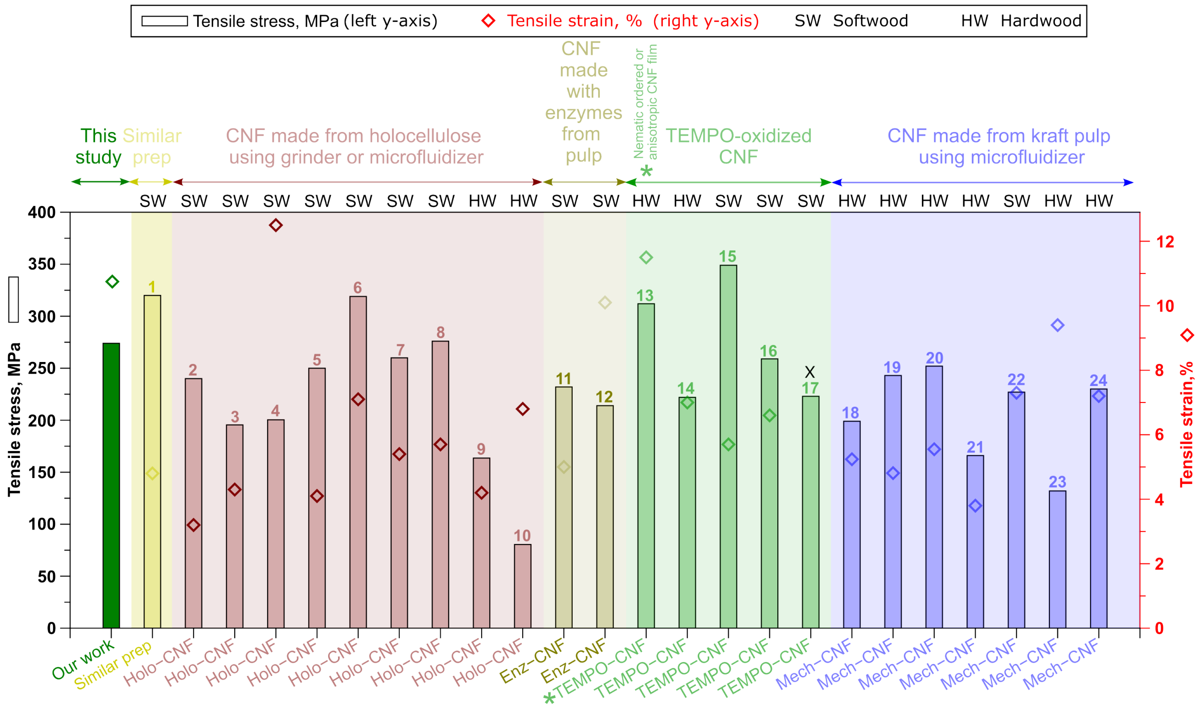

**Figure S23.** Benchmarking tensile properties of hCNF$_{Bn\ \mathrm{WT}}$ films against literature reports. Comparison of the tensile strength (left y-axis) and tensile strain (right y-axis) of hCNF$_{Bn\ \mathrm{WT}}$ films derived from *Brassica napus* (a commercially relevant lignocellulosic feedstock in the UK) with previously reported cellulose-based films. Among the references, only data point 13 exhibited a combined tensile strength exceeding 250 MPa and strain greater than 10%, surpassing the present work; this performance has been attributed to nematic or anisotropic ordering within the film, enabling both high strength and enhanced extensibility. "X" on data point 17 denotes strain data were not available. Literature references correspond to: 1: [7]; 2: [35]; 3-4: [19]; 5-8: [20]; 9-10: [19]; 11: [36]; 12: [37]; 13-14: [22]; 15-16: [38]; 17: [39]; 18-21: [18]; 22: [21]; 23: [40]; 24: [41]

#### 8. Statistical analysis

Statistical analyses were performed using QtiPlot. Data are reported as mean ± standard deviation (SD). Statistical significance was assessed by one-way analysis of variance (ANOVA) followed by Tukey's *post hoc* test for multiple comparisons. Pairwise comparison results are presented using either compact letter displays (CLDs) or p values. Means that do not share a common letter are significantly different according to Tukey's test at the 5% significance level. Differences were considered statistically significant at $p < 0.05$.

#### 9. Language editing

QuillBot was used to improve the readability and clarity of the text in a prefinal version of the manuscript. The text was subsequently reviewed and edited for preparation of the final version.

**10. References**

1. J. C. Mortimer, G. P. Miles, D. M. Brown, et al., "Absence of Branches from Xylan in Arabidopsis Gux Mutants Reveals Potential for Simplification of Lignocellulosic Biomass," *Proceedings of the National Academy of Sciences 107*, no. 40 (2010): 17409–17414, https://doi.org/10.1073/pnas.1005456107.
2. T. Tryfona, Y. Pankratova, D. Petrik, et al., "Altering the Substitution and Cross-Linking of Glucuronoarabinoxylans Affects Cell Wall Architecture in Brachypodium Distachyon," *New Phytologist 242*, no. 2 (2024): 524–543, https://doi.org/10.1111/nph.19624.
3. T. Tryfona, H.-C. Liang, T. Kotake, Y. Tsumuraya, E. Stephens, and P. Dupree, "Structural Characterization of Arabidopsis Leaf Arabinogalactan Polysaccharides," *Plant Physiology 160*, no. 2 (2012): 653–666, https://doi.org/10.1104/pp.112.202309.
4. F. Lu, C. Wang, M. Chen, F. Yue, and J. Ralph, "A Facile Spectroscopic Method for Measuring Lignin Content in Lignocellulosic Biomass," *Green Chemistry 23*, no. 14 (2021): 5106–5112, https://doi.org/10.1039/d1gc01507a.
5. A. Sluiter, B. Hames, R. Ruiz, C. Scarlata, J. Sluiter, and D. Templeton, "Determination of Structural Carbohydrates and Lignin in Biomass Determination of Structural Carbohydrates and Lignin in Biomass," *National Renewable Energy Laboratory (NREL) 2011*, no. June (2010), https://doi.org/NREL/TP-510-42618.
6. J. J. Lyczakowski, K. B. Wicher, O. M. Terrett, et al., "Removal of Glucuronic Acid from Xylan Is a Strategy to Improve the Conversion of Plant Biomass to Sugars for Bioenergy," *Biotechnology for Biofuels 10*, no. 1 (2017), https://doi.org/10.1186/s13068-017-0902-1.
7. X. Yang, M. S. Reid, P. Olsén, and L. A. Berglund, "Eco-Friendly Cellulose Nanofibrils Designed by Nature: Effects from Preserving Native State," *ACS Nano 14*, no. 1 (2020): 724–735, https://doi.org/10.1021/acsnano.9b07659.
8. J. Schindelin, I. Arganda-Carreras, E. Frise, et al., "Fiji: An Open-Source Platform for Biological-Image Analysis," *Nature Methods 9*, no. 7 (2012): 676–682, https://doi.org/10.1038/nmeth.2019.
9. T. J. Simmons, J. C. Mortimer, O. D. Bernardinelli, et al., "Folding of Xylan onto Cellulose Fibrils in Plant Cell Walls Revealed by Solid-State NMR," *Nature Communications 7*, no. 1 (2016): 13902, https://doi.org/10.1038/ncomms13902.
10. G. Metz, X. L. Wu, and S. O. Smith, "Ramped-Amplitude Cross Polarization in Magic-Angle-Spinning NMR," *Journal of Magnetic Resonance, Series A 110*, no. 2 (1994): 219–227, https://doi.org/10.1006/jmra.1994.1208.
11. B. M. Fung, A. K. Khitrin, and K. Ermolaev, "An Improved Broadband Decoupling Sequence for Liquid Crystals and Solids," *Journal of Magnetic Resonance 142*, no. 1 (2000): 97–101, https://doi.org/10.1006/jmre.1999.1896.
12. A. Lesage, M. Bardet, and L. Emsley, "Through-Bond Carbon−Carbon Connectivities in Disordered Solids by NMR," *Journal of the American Chemical Society 121*, no. 47 (1999): 10987–10993, https://doi.org/10.1021/ja992272b.
13. F. Fayon, D. Massiot, M. H. Levitt, et al., "Through-Space Contributions to Two-Dimensional Double-Quantum J Correlation NMR Spectra of Magic-Angle-Spinning Solids," *The Journal of Chemical Physics 122*, no. 19 (2005): 194313, https://doi.org/10.1063/1.1898219.
14. K. Takegoshi, S. Nakamura, and T. Terao, "13C–1H Dipolar-Assisted Rotational Resonance in Magic-Angle Spinning NMR," *Chemical Physics Letters 344*, no. 5 (2001): 631–637, https://doi.org/10.1016/S0009-2614(01)00791-6.
15. S. Koskela, S. Wang, D. Xu, et al., "Lytic Polysaccharide Monooxygenase (LPMO) Mediated Production of Ultra-Fine Cellulose Nanofibres from Delignified Softwood Fibres," *Green Chemistry 21*, no. 21 (2019): 5924–5933, https://doi.org/10.1039/C9GC02808K.

16. A. Villares, C. Moreau, C. Bennati-Granier, et al., "Lytic Polysaccharide Monooxygenases Disrupt the Cellulose Fibers Structure," *Scientific Reports 7*, no. 1 (2017): 40262, https://doi.org/10.1038/srep40262.
17. M. Pääkkö, M. Ankerfors, H. Kosonen, et al., "Enzymatic Hydrolysis Combined with Mechanical Shearing and High-Pressure Homogenization for Nanoscale Cellulose Fibrils and Strong Gels," *Biomacromolecules 8*, no. 6 (2007): 1934–1941, https://doi.org/10.1021/bm061215p.
18. S. Arola, J. M. Malho, P. Laaksonen, M. Lille, and M. B. Linder, "The Role of Hemicellulose in Nanofibrillated Cellulose Networks," *Soft Matter 9*, no. 4 (2013): 1319–1326, https://doi.org/10.1039/c2sm26932e.
19. D. M. de Carvalho, C. Moser, M. E. Lindström, and O. Sevastyanova, "Impact of the Chemical Composition of Cellulosic Materials on the Nanofibrillation Process and Nanopaper Properties," *Industrial Crops and Products 127* (2019): 203–211, https://doi.org/10.1016/j.indcrop.2018.10.052.
20. S. Galland, F. Berthold, K. Prakobna, and L. A. Berglund, "Holocellulose Nanofibers of High Molar Mass and Small Diameter for High-Strength Nanopaper," *Biomacromolecules 16*, no. 8 (2015): 2427–2435, https://doi.org/10.1021/acs.biomac.5b00678.
21. K. Prakobna, S. Galland, and L. A. Berglund, "High-Performance and Moisture-Stable Cellulose–Starch Nanocomposites Based on Bioinspired Core–Shell Nanofibers," *Biomacromolecules 16*, no. 3 (2015): 904–912, https://doi.org/10.1021/bm5018194.
22. T. Saito, M. Hirota, N. Tamura, et al., "Individualization of Nano-Sized Plant Cellulose Fibrils by Direct Surface Carboxylation Using TEMPO Catalyst under Neutral Conditions," *Biomacromolecules 10*, no. 7 (2009): 1992–1996, https://doi.org/10.1021/BM900414T.
23. R. Tanaka, T. Saito, T. Hänninen, et al., "Viscoelastic Properties of Core–Shell-Structured, Hemicellulose-Rich Nanofibrillated Cellulose in Dispersion and Wet-Film States," *Biomacromolecules 17*, no. 6 (2016): 2104–2111, https://doi.org/10.1021/acs.biomac.6b00316.
24. Y. Ono, M. Takeuchi, Y. Zhou, and A. Isogai, "Characterization of Cellulose and TEMPO-Oxidized Celluloses Prepared from Eucalyptus Globulus," *Holzforschung 76*, no. 2 (2022): 169–178, https://doi.org/10.1515/hf-2021-0159.
25. Y. Zhou, Y. Ono, M. Takeuchi, and A. Isogai, "Changes to the Contour Length, Molecular Chain Length, and Solid-State Structures of Nanocellulose Resulting from Sonication in Water," *Biomacromolecules 21*, no. 6 (2020): 2346–2355, https://doi.org/10.1021/acs.biomac.0c00281.
26. T. C. F. Gomes, and M. S. Skaf, "Cellulose-Builder: A Toolkit for Building Crystalline Structures of Cellulose," *Journal of Computational Chemistry 33*, no. 14 (2012): 1338–1346, https://doi.org/10.1002/jcc.22959.
27. O. M. Terrett, J. J. Lyczakowski, L. Yu, et al., "Molecular Architecture of Softwood Revealed by Solid-State NMR," *Nature Communications 10*, no. 1 (2019), https://doi.org/10.1038/s41467-019-12979-9.
28. A. N. Fernandes, L. H. Thomas, C. M. Altaner, et al., "Nanostructure of Cellulose Microfibrils in Spruce Wood," *Proceedings of the National Academy of Sciences 108*, no. 47 (2011): E1195–E1203, https://doi.org/10.1073/pnas.1108942108.
29. B. Song, S. Zhao, W. Shen, C. Collings, and S. Y. Ding, "Direct Measurement of Plant Cellulose Microfibril and Bundles in Native Cell Walls," *Frontiers in Plant Science 11* (2020): 479, https://doi.org/10.3389/fpls.2020.00479.
30. W. Humphrey, A. Dalke, and K. Schulten, "VMD: Visual Molecular Dynamics," *Journal of Molecular Graphics 14*, no. 1 (1996): 33–38, https://doi.org/10.1016/0263-7855(96)00018-5.
31. O. Guvench, S. S. Mallajosyula, E. P. Raman, et al., "CHARMM Additive All-Atom Force Field for Carbohydrate Derivatives and Its Utility in Polysaccharide and Carbohydrate–

Protein Modeling," *Journal of Chemical Theory and Computation 7*, no. 10 (2011): 3162–3180, https://doi.org/10.1021/ct200328p.
32. G. S. Park, "Transport Principles—Solution, Diffusion and Permeation in Polymer Membranes," in *Synthetic Membranes: Science, Engineering and Applications*, edited by P. M. Bungay, H. K. Lonsdale, and M. N. de Pinho, Springer Netherlands: Dordrecht 1986, pp. 57–107, https://doi.org/10.1007/978-94-009-4712-2_3.
33. C. J. Lomauro, A. S. Bakshi, and T. P. Labuza, "Evaluation of Food Moisture Sorption Isotherm Equations Part I: Fruit, Vegetable and Meat Products," *LWT - Food Science and Technology 18*, no. 2 (1985): 111–117.
34. T. Zhang, H. Tang, D. Vavylonis, and D. J. Cosgrove, "Disentangling Loosening from Softening: Insights into Primary Cell Wall Structure," *The Plant Journal 100*, no. 6 (2019): 1101–1117, https://doi.org/10.1111/tpj.14519.
35. S. Iwamoto, K. Abe, and H. Yano, "The Effect of Hemicelluloses on Wood Pulp Nanofibrillation and Nanofiber Network Characteristics," *Biomacromolecules 9*, no. 3 (2008): 1022–1026, https://doi.org/10.1021/bm701157n.
36. H. Sehaqui, A. Liu, Q. Zhou, and L. A. Berglund, "Fast Preparation Procedure for Large, Flat Cellulose and Cellulose/Inorganic Nanopaper Structures," *Biomacromolecules 11*, no. 9 (2010): 2195–2198, https://doi.org/10.1021/bm100490s.
37. M. Henriksson, L. A. Berglund, P. Isaksson, T. Lindström, and T. Nishino, "Cellulose Nanopaper Structures of High Toughness," *Biomacromolecules 9*, no. 6 (2008): 1579–1585, https://doi.org/10.1021/bm800038n.
38. M. Zhao, F. Ansari, M. Takeuchi, et al., "Nematic Structuring of Transparent and Multifunctional Nanocellulose Papers," *Nanoscale Horizons 3*, no. 1 (2017): 28–34, https://doi.org/10.1039/C7NH00104E.
39. M. Nogi, S. Iwamoto, A. N. Nakagaito, and H. Yano, "Optically Transparent Nanofiber Paper," *Advanced Materials 21*, no. 16 (2009): 1595–1598, https://doi.org/10.1002/adma.200803174.
40. M. Farooq, T. Zou, G. Riviere, M. H. Sipponen, and M. Österberg, "Strong, Ductile, and Waterproof Cellulose Nanofibril Composite Films with Colloidal Lignin Particles," *Biomacromolecules 20*, no. 2 (2019): 693–704, https://doi.org/10.1021/acs.biomac.8b01364.
41. M. Österberg, J. Vartiainen, J. Lucenius, et al., "A Fast Method to Produce Strong NFC Films as a Platform for Barrier and Functional Materials," *ACS Applied Materials & Interfaces 5*, no. 11 (2013): 4640–4647, https://doi.org/10.1021/am401046x.
